# Multiscale and multimodal reconstruction of cortical structure and function

**DOI:** 10.1101/2020.10.14.338681

**Authors:** Nicholas L. Turner, Thomas Macrina, J. Alexander Bae, Runzhe Yang, Alyssa M. Wilson, Casey Schneider-Mizell, Kisuk Lee, Ran Lu, Jingpeng Wu, Agnes L. Bodor, Adam A. Bleckert, Derrick Brittain, Emmanouil Froudarakis, Sven Dorkenwald, Forrest Collman, Nico Kemnitz, Dodam Ih, William M. Silversmith, Jonathan Zung, Aleksandar Zlateski, Ignacio Tartavull, Szi-chieh Yu, Sergiy Popovych, Shang Mu, William Wong, Chris S. Jordan, Manuel Castro, JoAnn Buchanan, Daniel J. Bumbarger, Marc Takeno, Russel Torres, Gayathri Mahalingam, Leila Elabbady, Yang Li, Erick Cobos, Pengcheng Zhou, Shelby Suckow, Lynne Becker, Liam Paninski, Franck Polleux, Jacob Reimer, Andreas S. Tolias, R. Clay Reid, Nuno Maçarico da Costa, H. Sebastian Seung

## Abstract

We present a semi-automated reconstruction of L2/3 mouse primary visual cortex from 3 million cubic microns of electron microscopic images, including pyramidal and inhibitory neurons, astrocytes, microglia, oligodendrocytes and precursors, pericytes, vasculature, mitochondria, and synapses. Visual responses of a subset of pyramidal cells are included. The data are being made publicly available, along with tools for programmatic and 3D interactive access. The density of synaptic inputs onto inhibitory neurons varies across cell classes and compartments. We uncover a compartment-specific correlation between mitochondrial coverage and synapse density. Frequencies of connectivity motifs in the graph of pyramidal cells are predicted quite accurately from node degrees using the configuration model of random graphs. Cells receiving more connections from nearby cells exhibit stronger and more reliable visual responses. These example findings illustrate the resource’s utility for relating structure and function of cortical circuits as well as for neuronal cell biology.

## Introduction

Connectomic reconstructions of neural tissue using electron microscopy (EM) are becoming more broadly available, and have been used to study ultrastructural features of specific neuronal cell types (Ribak and Anderson, 1980; Spacek and Harris, 1997; Karube, Kubota and Kawaguchi, 2004; Wu *et al.*, 2017), connectivity rules (Kasthuri *et al.*, 2015; Lee *et al.*, 2016), plasticity rules (Bartol *et al.*, 2015; Bloss *et al.*, 2018; Dorkenwald *et al.*, 2019; Kornfeld *et al.*, 2020), circuit development (Wilson *et al.*, 2019), and sensory processing (Kim *et al.*, 2014; Takemura, Nern, *et al.*, 2017; Vishwanathan *et al.*, 2017; Wanner and Friedrich, 2020). Reconstructions have probed circuitry in various parts of the nervous system for multiple species, including songbird (Kornfeld *et al.*, 2017, 2020), zebrafish (Vishwanathan *et al.*, 2017; Wanner and Friedrich, 2020), rodent retina, cortex, and cerebellum (Helmstaedter *et al.*, 2013; Kasthuri *et al.*, 2015; Behrens *et al.*, 2016; Morgan, 2017; Schmidt *et al.*, 2017; Bae *et al.*, 2018; Dorkenwald *et al.*, 2019; Motta *et al.*, 2019; Wilson *et al.*, 2019; Schneider-Mizell *et al.*, 2020), as well as *Drosophila melanogaster* (Takemura, Aso, *et al.*, 2017; Zheng *et al.*, 2018; Dorkenwald *et al.*, 2020; Shan Xu *et al.*, 2020).

The scale of reconstructions has been a main limitation of EM-based connectomics. Here we present a new resource for neuroscientific discovery, a reconstruction of roughly 3 million cubic microns of mouse visual cortex that includes both neuronal and non-neuronal cells, as well as subcellular (mitochondria) and intercellular (synapses) elements. The reconstruction is accompanied by activity recordings of pyramidal cells (PyCs) with 2-photon calcium imaging. Two previous studies combining calcium imaging and electron microscopy in cortex (Bock *et al.*, 2011a; Lee *et al.*, 2016) were limited to only tens of cells. Our automated reconstruction contains hundreds of somas, including those of 363 PyCs and 34 non-pyramidal neurons. The data are being made publicly available as a resource, along with tools for programmatic and 3D interactive access (microns-explorer.org).

The present paper complements two companion reports (Dorkenwald *et al.*, 2019; Schneider-Mizell *et al.*, 2020) that apply this resource to address specific scientific questions. The present work fully describes the resource, and showcases its usefulness through a broad range of scientific analyses.

Our first analysis focuses on the 34 non-pyramidal cells, presumed inhibitory, with somas captured in our EM volume. We categorized these cells into anatomically-defined classes where possible, and measured the density and size of synapses formed on their dendrites and somas. Our comparison of these properties across classes and compartments extends the sparse reports in the literature (Gulyás *et al.*, 1999a; Martina, Vida and Jonas, 2000a; Kameda *et al.*, 2012).

We then analyze properties of ~180,000 mitochondria within a subset of PyCs in our dataset. Building on previous reports that axonal mitochondria tend to be smaller than dendritic mitochondria (Chang, Honick and Reynolds, 2006; Lewis *et al.*, 2018), we show that somatic mitochondria are statistically intermediate in size. We also investigate the correlation between mitochondrial coverage and synapse density (Li *et al.*, 2004; Dickey and Strack, 2011; Bertholet *et al.*, 2013), and show that the correlation exhibits some compartment-specificity.

The above two analyses illustrate the multiscale nature of the resource. The high resolution (nanometers) of the EM images allows synapses and mitochondria to be identified and segmented accurately. The large field of view (hundreds of microns) enables the annotation of these elements with contextual information (cell type and cellular compartment).

Previous studies employing synaptic physiology in cortical brain slices reported “highly nonrandom” connectivity motifs (Song *et al.*, 2005; Perin, Berger and Markram, 2011), high network transitivity, and a common neighbor rule (Perin, Berger and Markram, 2011). Such findings were influential, and interpreted as evidence for various cortical circuit theories, such as Hebbian cell assemblies (Perin, Berger and Markram, 2011), neuron clustering (Litwin-Kumar and Doiron, 2012; Klinshov *et al.*, 2014), and attractor networks (Brunel, 2016; Zhang, Zhang and Stepanyants, 2019).

The claim of “nonrandomness” was relative to the Erdős-Rényi (ER) model of random graphs, which considers connections between all pairs of nodes to be equally probable. Using our PyC graph, we find that the frequencies of two-cell and three-cell motifs can deviate significantly from the ER model, similar to previous reports, but deviations from the “configuration model” are smaller. In the latter model, the random graphs are constrained to have the same degree sequence as the reconstructed PyC graph (Newman, 2018). We use the directed version of the configuration model, which constrains both in-degree, the number of presynaptic partners of a cell, and out-degree, the number of postsynaptic partners of a cell.

We are able to analyze three-cell motifs because our reconstructed PyC graph is large enough to include nonzero counts of even rare motifs. This is a milestone in scale for cortical connectomics, and was attainable because of the large field of view and acceleration of reconstruction by automation. That being said, our motif frequencies are still affected by truncation of arbors by the borders of the reconstructed volume. Comparison with previous studies in brain slices (Song *et al.*, 2005; Perin, Berger and Markram, 2011) is difficult because the geometry of arbor truncation differs from our EM reconstruction. Still, our analysis with the configuration model suggests that “nonrandomness” may be more subtle than previously supposed, and that further research will be needed to draw theoretical conclusions from empirical data about cortical connectivity motifs.

The degree of a node plays a central role in the configuration model, and is a structural property of cortical circuits. We are able to explore the relation of degree with visual function, since many of the reconstructed PyCs come with visual responses from calcium imaging. To estimate the tendency of a PyC to receive local connections from its neighbors, we define the “in-connection density” as the number of its presynaptic partners (“in-degree”) in the PyC graph, divided by the total length of its dendritic arbor. The normalization is intended to compensate for the varying amount by which arbors are cut off by the borders of the EM volume. We find that orientation-tuned cells with higher in-connection density tend to exhibit stronger responses to oriented stimuli, consistent with the hypothesis of local recurrent excitation as a cortical amplifier (Douglas *et al.*, 1995; Lien and Scanziani, 2013; Li *et al.*, 2013). We also find that cells with higher in-connection density tend to respond in a more reliable manner.

Our multimodal resource should make it possible to discover further relationships between neural connectivity and activity, and advances the trend of performing calcium imaging prior to EM (Bock *et al.*, 2011a; Briggman, Helmstaedter and Denk, 2011; Lee *et al.*, 2016; Vishwanathan *et al.*, 2017; Bae et al., 2018; Wanner and Friedrich, 2020), a powerful approach for relating structure and function of neural circuits.

## Results

### Reconstruction of mouse visual cortex

We acquired a 250×140×90 μm^3^ serial section electron microscopy (ssEM) dataset from L2/3 primary visual cortex (V1) of a P36 male mouse. Alignment of serial section images followed by automatic segmentation yielded more than 8 million objects. Computational skeletons and meshes were automatically generated for each object in the volume to provide useful representations for visualization and analysis. We identified all neuronal and non-neuronal cells with somas in the volume using a semi-automated approach (STAR Methods). Neurons included 363 pyramidal cells (PyCs) and 34 nonpyramidal cells, presumed inhibitory (Figure 1A).

**Figure 1:**
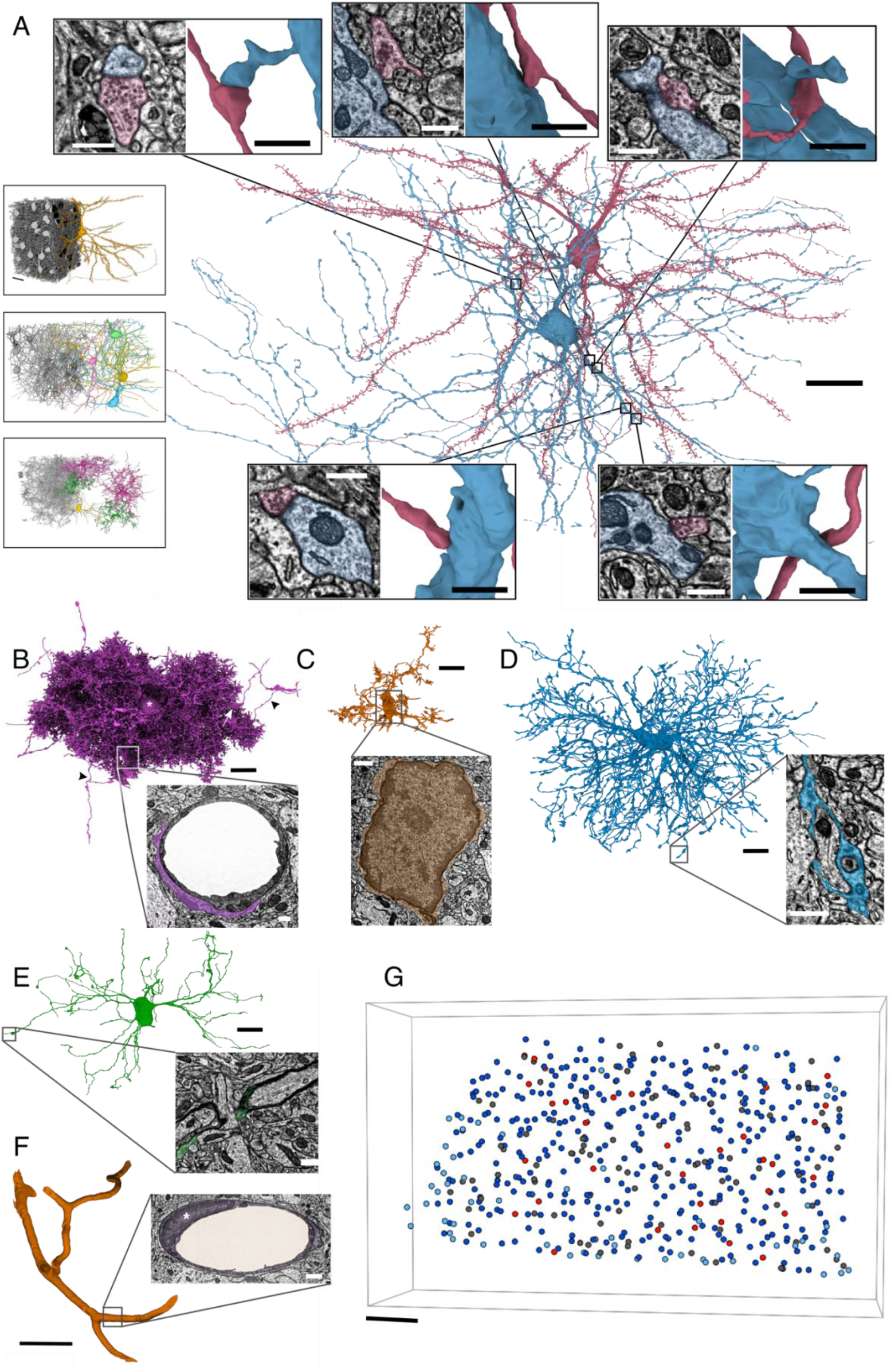
Neuronal and non-neuronal cells. (A) 3D Rendering of pyramidal neuron (red) and basket interneuron (blue) after correction of segmentation errors. Top/Bottom insets: zoomed-in 3D renderings and 2D EM cross-section of 5 synapses formed by the basket cell onto the pyramidal cell. Left insets: all cells of a classified type cut away to reveal a few examples. Top to bottom: PyCs, interneurons, non-astrocytic glia. Four cells were removed from the glia inset due to merge errors that obscure the view (STAR Methods). (B) An astrocyte without proofreading. Extensions onto endfeet (box, inset, arrow) as well as merge errors with axon segments (arrowheads) are visible. Asterisk: soma. (C) A microglia without proofreading. Inset: 2D EM cross-section of microglial soma, showing dark, scarce cytoplasm and dark nucleoplasm. (D) Oligodendrocyte precursor cell (OPC). Inset: EM cross-section showing an OPC process occupying the small space between neurites. (E) Oligodendrocyte. Inset: EM cross-section showing a process transitioning to myelin. (F) Blood vessel after removing split errors (merge errors removed for clarity; see Data S3). Inset: EM cross-section showing blood vessel wrapped by an endothelial cell (purple). Asterisk: endothelial cell nucleus. (G) Locations of all somas in the EM volume, colored by type. Dark blue: PyCs analyzed in this paper; light blue: PyCs at edge of volume that were not analyzed; red: analyzed inhibitory neurons; dark gray: glia. Scale bars: (A) 20 μm, cell type insets, 25 μm, 3D synapse insets, 1 μm, EM insets, 500 nm; (B-E) 10 μm, insets, 750 nm; (F) 20 μm, inset 100 nm; (G) 25 μm. Insets in panel (A) show cells from the back of the dataset (all others show cells from the front). See also Figure S1 and Data S3.

We attempted to classify the 34 inhibitory cells based on axon morphology (STAR Methods). We identified 12 bipolar cells, 4 basket cells, 2 chandelier cells, 1 Martinotti cell, and 1 neurogliaform cell (Figure 2A and Figure S1). There were 14 cells that we could not confidently classify; all of them lacked a significant axon within the bounds of the reconstruction, being severely cut off by the borders of the EM volume (Figure 3A and Data S1).

**Figure 2.**
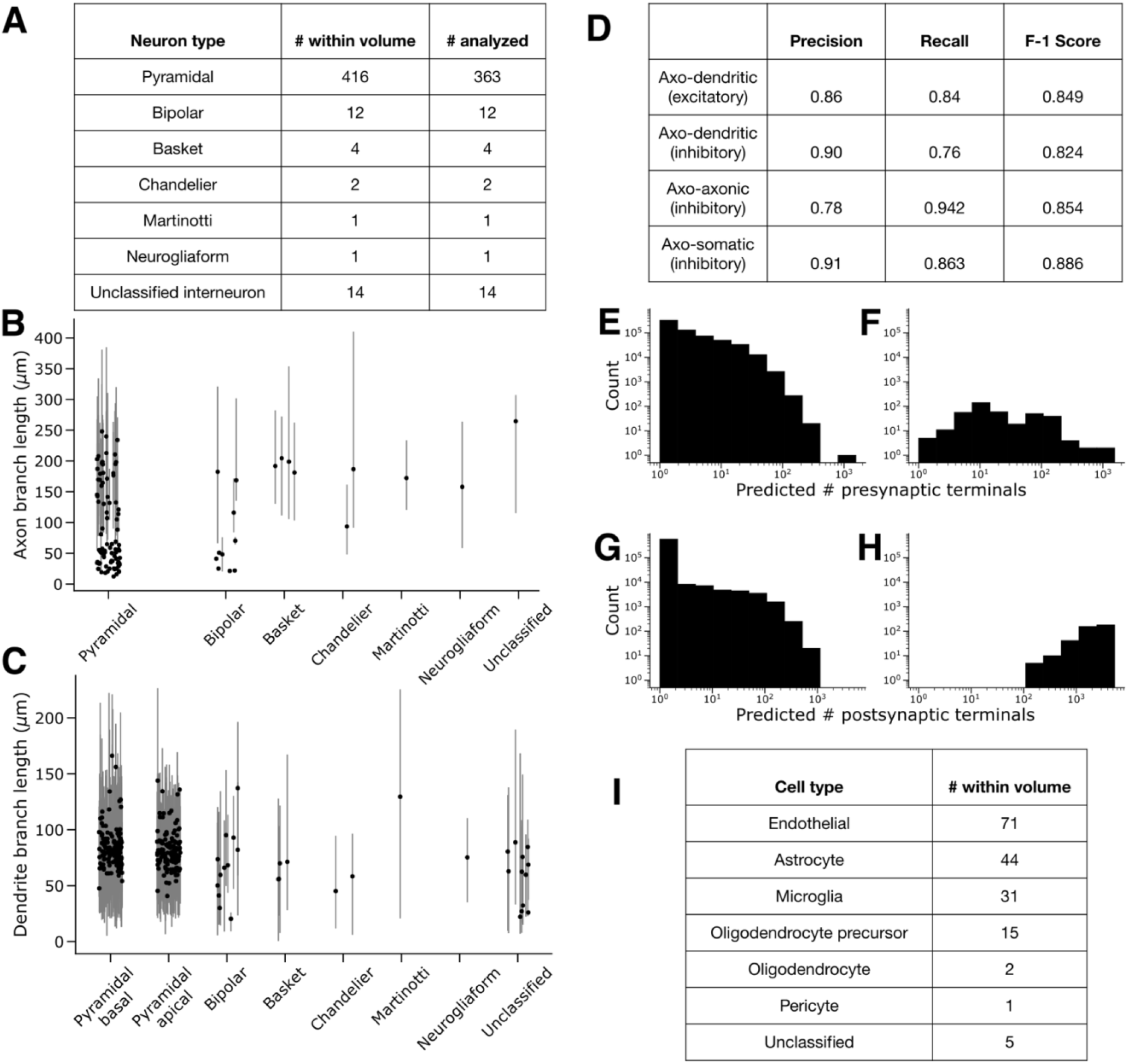
Resource Statistics. (A) Neuron type classifications. (B-C) Captured neurite branch length for proofread cells with soma, measured as path length between each soma and its skeleton leaf nodes (STAR Methods). Points show median branch length for each cell. Lines show 5th and 95th percentiles. Pyramidal cells with ideal compartment labels are shown (*n*=125, STAR Methods). (B) Captured axon branch length per cell (C) Captured dendrite branch length per cell. (D) Estimated synapse detection performance on synapse types (STAR Methods). (E-F) Inferred number of presynaptic terminals per reconstructed segment. (E) Purely-presynaptic orphans (*n*=612201). (F) Cells with a soma in the volume (*n*=397). (G-H) Inferred number of postsynaptic terminals per reconstructed segment. (G) Purely-postsynaptic orphans (*n*=640285). (H) Cells with a soma in the volume (*n*=397). (I) Non-neuronal cell type classifications.

**Figure 3:**
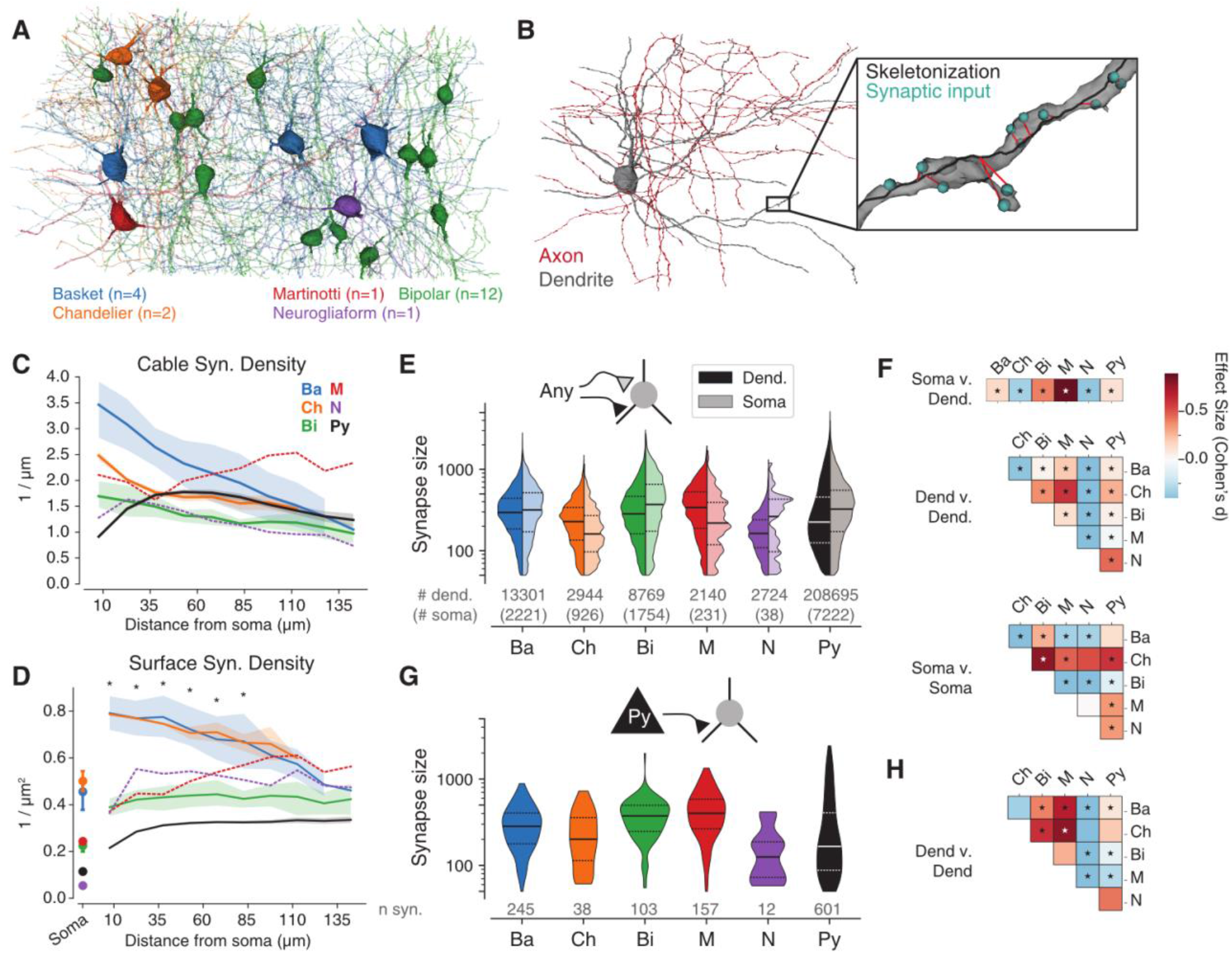
Cell type dependent properties of inhibitory neuron inputs. (A) Classified inhibitory interneurons. (B) Neuronal compartment labels and synapses for an example basket cell. (C) Linear synapse density for each cell type. (D) Surface synapse density for each cell type. Dots indicate value at the soma. Asterisks denote a statistically significant difference between basket and chandelier cells vs. the rest (two-tailed Mann-Whitney U for bins within 85 μm: p<0.001). (E) Synapse size distributions (in voxels) for dendrites and somata of each cell type (ANOVA dendritic synapses p≈0; somatic synapses: p≈0, STAR Methods). (F) Effect size of two-tailed Mann-Whitney U tests between distributions in (E) with Bonferroni correction. (G) Synapse size for inputs from L2/3 pyramidal cells onto dendrites of each cell type. (H) Effect size of two-tailed Mann-Whitney U tests between distributions in (G) with Bonferroni correction. Error bars and shaded regions in (C) and (D) indicate 95% bootstrap intervals. See also Figure S2, Figure S3, and Data S1.

Non-neuronal cells included 71 endothelial cells, 44 astrocytes, 31 microglia, 15 oligodendrocyte precursor cells (OPCs), 2 oligodendrocytes, and 1 pericyte (Figures 1B–1G and Figure 2I). Blood vessels were also reconstructed as a byproduct of the cellular segmentation: there is 2.03 mm of vasculature in the volume (Data S3).

Automatic synapse detection yielded over 3 million synapses throughout the volume. We estimated its performance as 93.0% precision and 90.9% recall within small test volumes (STAR Methods). Axo-dendritic, axo-axonic, and axo-somatic synapses are all included (Figures S1A-C), and we estimated detection performance for these synapse types (Figure 2D and STAR Methods). Synaptically coupled pairs of neurons include PyCs, inhibitory cells, and combinations of these types (Figure 1A, Figure 3A, and Figures S1A-C).

The reconstruction includes over 2.4 million mitochondria with a total path length of over 3.7 m in the EM volume (Figures 4A, 4B, and STAR Methods; see Dorkenwald *et al.*, 2017; Haberl *et al.*, 2018; Xiao et al., 2018 for similar automated methods and Perez *et al.*, 2014; Márquez Neila *et al.*, 2016; Calì *et al.*, 2019; Yuan *et al.*, 2020 for others). Each reconstructed mitochondrion is annotated with an identifier for its host cell. Some dendritic mitochondria have “reticular” morphologies (Popov *et al.*, 2005) that double back on themselves multiple times and contain Y- and H-junctions (Figure 4C). These reticular mitochondria extend for longer distances (>100 μm) in our volume than previously observed. They reside in orphan dendrites with somas outside the EM volume, presumably in deep L2/3, L5 or L6.

**Figure 4:**
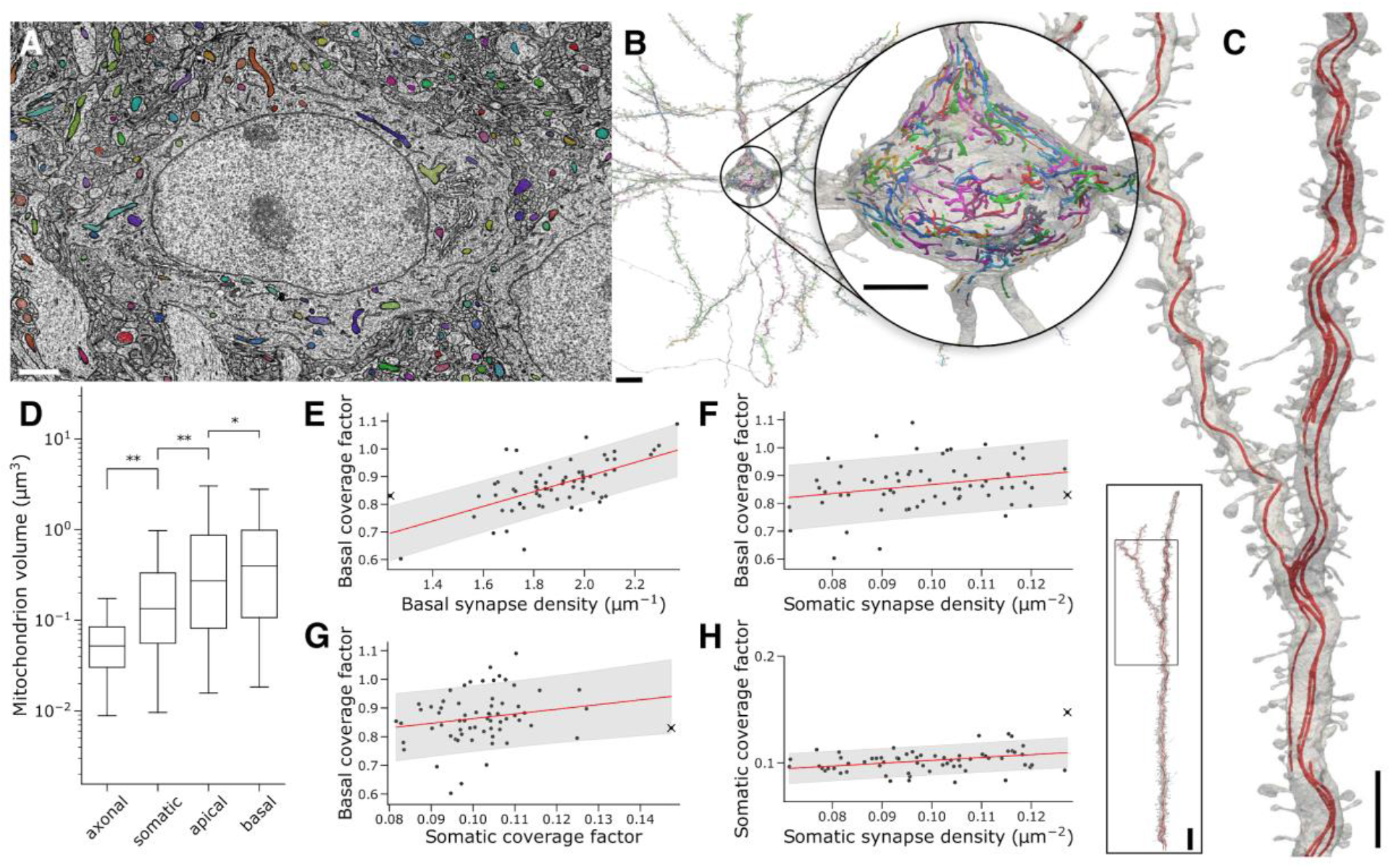
Computational reconstructions of mitochondria in layer 2/3 PyCs. (A) Sample cutout of mitochondrial segmentation. Scale bar 1.5 μm. (B) Single PyC rendered with its overlapping mitochondria. Inset shows its soma. Scale bars: 10 μm, inset 5 μm. (C) Detailed view of a single apical mitochondrion within a pyramidal cell from a deeper cortical layer. Inset shows the morphology of the captured dendrite (grey) and mitochondria (red). Scale bars: 5 μm, inset 10 μm. (D) Comparison of mitochondrial volume by neuronal compartment (One-tailed Mann-Whitney U tests; *: p<2.8 × 10^−26^, **: p≈0). Boxes indicate interquartile ranges, and whiskers indicate the 5th and 95th percentiles. (E) Basal mitochondrial coverage (mitochondrial length divided by neurite length) is correlated with basal synapse density (Pearson’s *r*=0.62, p<3.3 × 10^−8^). (F) Basal mitochondrial coverage factor is only weakly correlated with somatic synapse density (*r*=0.26, p=0.036). (G) No significant correlation between mitochondrial coverage factor in basal dendrites and soma. (*r*=0.20, p=0.104). (H) Somatic synapse density is only weakly correlated with somatic mitochondrial coverage (mitochondrial volume divided by somatic cytosol volume). (*r*=0.34, p=0.005). Outlier cell marked as an ‘X’ within (E-H). Our findings are robust to removal of this cell (Data S2). Shaded regions in (E-H) show the 80% prediction interval. See also Figure S4 and Data S2.

We made several improvements to the automated reconstruction to facilitate its analysis. Segmentation errors in the 363 PyCs and 34 inhibitory cells with somas were corrected using an interactive proofreading system that enabled human experts to split and merge objects. This proofreading allows the study of synapses that involve the soma, as well as their axons and dendrites that each extend up to 400 μm in path length from the soma (Figures 2B, 2C, 2F and 2H). Most PyCs have over half of their expected basal dendrite path length for cells in this region (Gilman, Medalla and Luebke, 2016), and many of these basal dendrites are completely reconstructed. Synapses between PyCs with somas were also proofread. This yielded a highly accurate map of connectivity between PyCs with somas in the EM volume, consisting of 1981 synapses within 1752 PyC-PyC connections. This map can be used to analyze properties of cortical circuits (Figure 5; Dorkenwald *et al.*, 2019).

**Figure 5:**
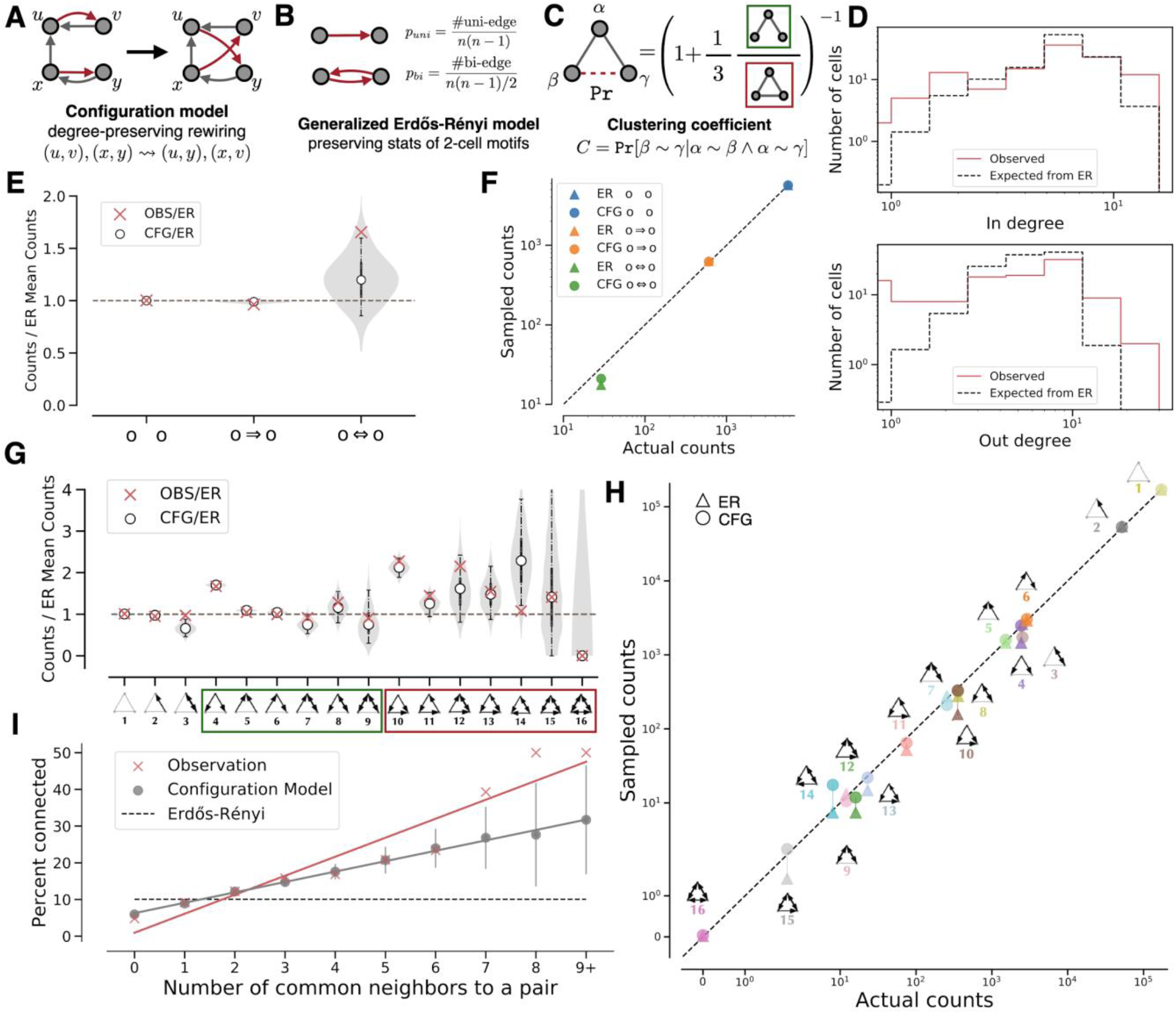
Connectivity motif frequencies can be predicted from degree sequences. (A) Sampling from the configuration model (see Artzy-Randrup and Stone 2005). In and out degrees for each neuron are preserved by exchanging connected partners. (B) Illustration of the generalized Erdős–Rényi (ER) model, where the frequency of bi-directional connections is held (*p*_*uni*_ ≈ 4.80 × 10^−2^, *p*_*bi*_ ≈ 4.58 × 10^−3^). (C) Illustration of the relationship between the clustering coefficient and 3-cell motif frequencies. (D) In and out degree distributions in the actual network and the expected distribution of in and out degrees in a regular ER model (edge probability = 0.0526). Red: observed data; Black, dashed: ER model. Histograms are calculated with 8 bins of the same widths on a log scale. The expected degree distributions are estimated from 100 samples from the ER model. (E) 2-cell motif frequencies in the actual network and a configuration model relative to the ER model. The shaded region shows the smoothed distribution of motif counts sampled from the configuration model. White points indicate medians, vertical lines indicate quartiles and the 95% confidence interval for 1,000 samples. (F) Comparison of 2-cell motif counts in the ER model and configuration model. Circles indicate mean counts sampled from the configuration model, and triangles indicate mean counts sampled from the ER model. (G) 3-cell motif frequencies in the actual network and the configuration model relative to a generalized ER model. The violin plot shows the smoothed distribution of the motif counts sampled from the configuration model. White points indicate medians, and lines indicate the 95% confidence interval for 1,000 samples. (H) Comparison of 3-cell motif counts in the generalized ER model and configuration model. Circles and triangles indicate mean counts sampled from the configuration model and the ER model, respectively. (I) The common neighbor rule is significantly more prominent in the pyramidal cell network than in generalized ER random networks. Gray: configuration model; Black, dashed: generalized ER random networks; Red: observed data. (red slope: r^2^ = 0.91, p=2.2e-5; gray: r^2^ ≈ 1.00, p=1.0e–10; z-test). Lines indicate standard deviation of 100 samples.See also Figure S5, Data S4, Data S5, Tables S1, and Table S2.

Neurites without a soma, termed “orphans”, may also be useful to study synaptic connectivity at a smaller spatial scale (Mishchenko *et al.*, 2010; Bloss *et al.*, 2018). Over 96,000 orphans possess at least 10 predicted synapses (Figures 2E and 2G), where each orphan’s synapses are uniformly presynaptic or postsynaptic. Sampling 200 of these orphans, we estimate that roughly 93% are free of significant merge errors.

Prior to EM imaging of the volume, responses of PyCs to visual stimuli were recorded in an overlapping volume by calcium imaging (Figure 6A and STAR Methods). Two types of visual stimuli were presented. A pink noise stimulus was shown to characterize spatiotemporal receptive fields (Figure 6B). Oriented stimuli moving in 16 directions were shown to characterize direction and orientation selectivity (Figures 6B and 6D). Each direction was shown once for each of 30 trials. Each trial presented the directions in a pseudo-random order, interspersed with the pink noise stimulus (Figure 6B and STAR Methods). The calcium videos were co-registered to the EM volume and activity traces were extracted for 112 of the 363 PyCs. Activity traces were extracted using the EASE algorithm (Zhou *et al.*, 2020), which uses reconstructed cells for initialization, so traces for only a subset of reconstructed cells that overlap with calcium imaging volume were acquired (STAR Methods).

**Figure 6.**
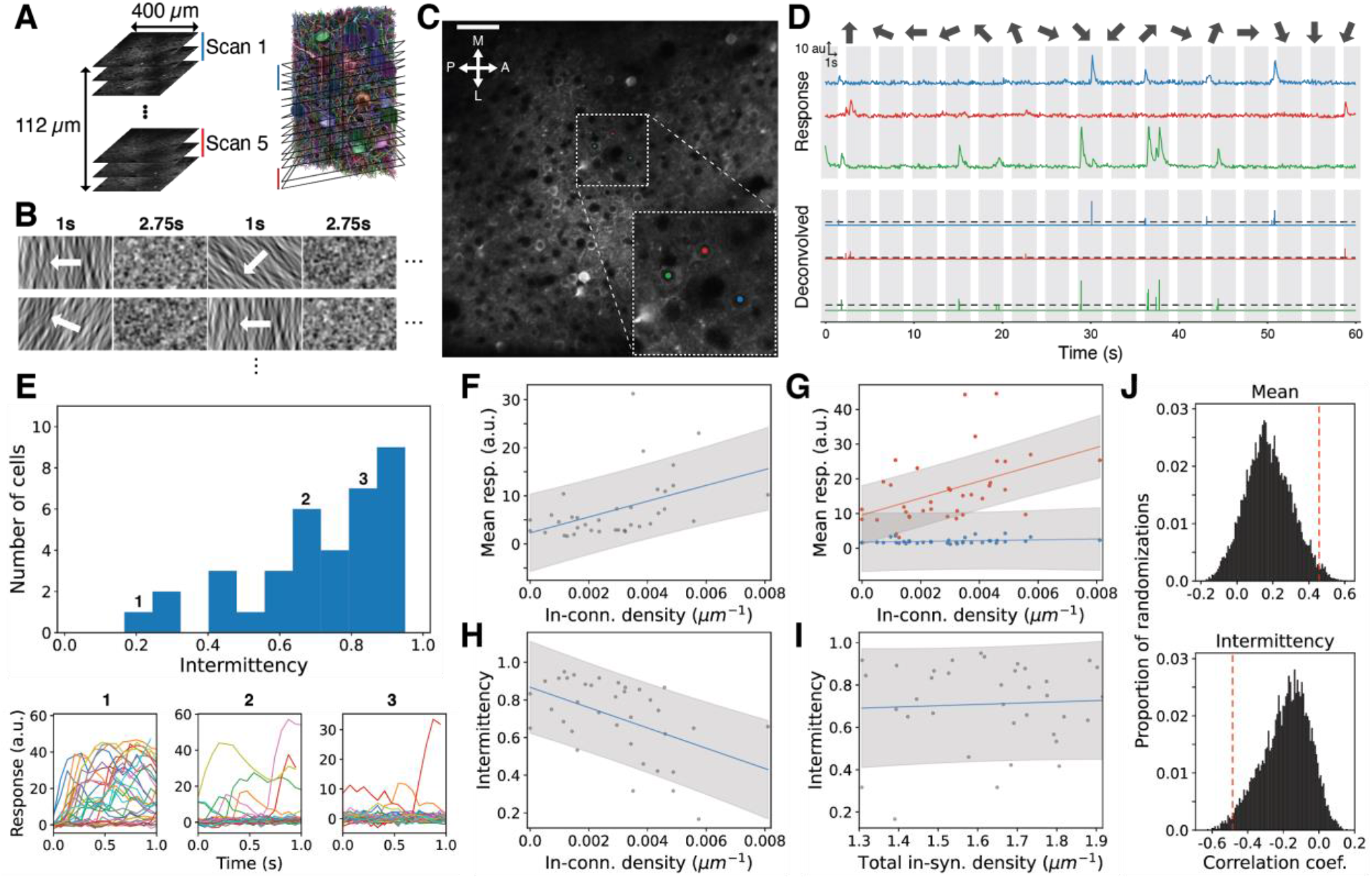
In-connection density is related to visual function. (A) 2-photon calcium recording overview and image plane coregistration. Black squares on the right are the image planes, cropped to only indicate the vertical location. (B) Visual stimulus presented during recording. Oriented stimuli moving in 16 directions were presented, interspersed with the pink noise stimulus, for 30 trials (row: single trial). (C) Example slice of 2-photon recording (A: Anterior, P: Posterior, M: Medial, L: Lateral) with the central area (white dashed box) zoomed in on the lower right. (D) Noise-normalized dF/F traces and deconvolved traces (dashed: activity threshold for “active” trials) for example cells marked in (C) during a single trial (white shade: directional stimulus shown, gray shade: noise stimulus shown). Directions are ordered pseudo-randomly in each trial. (E) Top: Distribution of intermittency, fraction of non-active trials, for orientation tuned cells. Bottom: Example cells’ responses to 30 trials of their preferred stimulus. Example cells are chosen from the bins labeled by numbers. (F) Mean response to preferred directions is positively correlated with in-connection density (*n*=36, Pearson’s r=0.46, p=0.005). (G) In-connection density has a greater effect on mean active responses (red, a=1, *n*=36) than mean inactive responses (blue, a=0, *n*=36, p=0.006). (H) In-connection density is negatively correlated with intermittency (*n*=36, r=-0.49, p=0.003). (I) Intermittency is not correlated with total in-synapse density (*n*=36, r=0.05, p=0.757). (J) Spatial location restricted permutation tests (Figure S7, STAR Methods) for correlations in (F) (top, p=0.0178) and (H) (bottom, p=0.018) to explain the correlation is not due to the spatial bias. (B) Scale bar: 50um. (F, G, H, I) Line: linear fit, Shade: 80% prediction interval (J) Red line: observed, All permutation tests used 10,000 iterations each. See also Figure S6 and Figure S7.

We have already made the EM data and reconstructed cells and synapses publicly available at microns-explorer.org, and the reconstructed mitochondria and visual responses will also be released in a timely fashion. We believe that this resource will be highly useful for the field, and publish interactive analysis code as a part of MicronsBinder (Collman, no date) to encourage its use. Below, we provide as evidence several “vignettes” that address diverse scientific questions about the cortex, on topics ranging from organelles to synapses to circuit structure and function. Each vignette is by necessity brief; they have been chosen to illustrate the breadth of science that is enabled by our resource.

### Spatial organization of synapses onto inhibitory cells

It is well-known that certain types of inhibitory neurons prefer to form synapses onto particular compartments of their postsynaptic partners (Kubota, 2014). Less study has been devoted to the spatial organization of synapses received by inhibitory neurons. Our reconstruction captured somas and dendrites of 34 inhibitory cells, and we assigned classical types to 20 of these (Figure 3A, Data S1, and STAR Methods). The somas and dendrites of these classified cells collectively have 55,493 incoming synapses in the reconstruction, and they display broadly similar properties across neurons within each type (Figure S3). Such information is not available from previous cortical EM reconstructions (Bock *et al.*, 2011b; Lee *et al.*, 2016; Motta *et al.*, 2019). In general, detailed study of dendrites of inhibitory cells has been limited to few cell classes across multiple brain regions and rodent species (Gulyás *et al.*, 1999b; Martina, Vida and Jonas, 2000b; Kawaguchi, Karube and Kubota, 2006; Kameda *et al.*, 2012).

The spatial organization of incoming synapses is important for understanding the biophysical mechanisms by which dendrites integrate their synaptic inputs. Compared to the dendritic biophysics of excitatory neurons (Stuart, Spruston and Häusser, 2016), the properties of inhibitory neurons are less well known (Hu and Vervaeke, 2018). Depending on the context, it can be more convenient to formulate passive cable theory (Rall, 1962) using either length-normalized or area-normalized membrane conductances. Both linear and areal densities of synapses are therefore of biological interest, with area-normalized values better accounting for dendritic radius and shape (Bloss *et al.*, 2016; Karimi *et al.*, 2020). We built both a linear (skeleton) and 3D mesh representation of inhibitory neuron somas and dendrites, computed both quantities, and examined their dependences on path length distance from the soma (Figures 3B–D, and Figure S2). We saw large differences in linear synapse density for different inhibitory cell classes, especially in the proximal dendrites. In this region, synapse density was highest for basket cells (3.5 synapses/μm; Figure 3C). For comparison, we also included data from a set of PyCs (*n*=65) that had full cell bodies in the volume (STAR Methods).

Areal synapse density as a function of distance from the soma also varied across cell classes (Figure 3D and Figure S2). The dependences for basket and chandelier cells were strikingly similar, even when cells originally unassigned due to an absence of axonal reconstruction were included as basket or chandelier cells (Figure S2H). Areal synapse densities on perisomatic dendrites in particular were much higher for basket and chandelier cells than for other types (two-tailed Mann-Whitney U basket and chandelier cells versus other interneurons, for bins within 85 μm: p<0.001). For all cell classes, the areal synapse density was lower in the soma than in the dendrites. Basket and chandelier cells possessed the highest somatic density values. Their values were up to 13 times greater than that of the neurogliaform cell, which had the lowest value (Figure 3D, Figure S2, and Figure S3).

Past work has shown that the sizes of synapses reflect their physiological strength and reliability (Bailey and Chen, 1983; Bourne and Harris, 2011; Bailey, Kandel and Harris, 2015; Holler-Rickauer *et al.*, 2019). Studying the sizes of the inhibitory cell inputs within our reconstruction may help to describe their cortical functions. We found that synapse size distributions differed significantly across cell classes (ANOVA for dendritic synapses: p≈0, somatic synapses: p≈0; Figures 3E and 3F). Within each type, dendritic and somatic size distributions differed significantly, although with small effect size in some cases (Two-tailed Mann-Whitney U test with Bonferroni correction: p_basket_<2.4 × 10^−7^, p_chandelier_<2.9 × 10^−25^, p_bipolar_<1.4 × 10^−26^, p_martinotti_=0.082, p_neurogliaform_<6.4 × 10^−10^, p_pyc_≈0; Figure 3F). PyC somatic inputs were larger than dendritic inputs, but this was not always the case for inhibitory cell types—bipolar cells received larger somatic synapses than dendritic synapses, whereas chandelier and Martinotti cells displayed the opposite. Such variation in synapse size distributions may be related to variation in functional properties and connectivity patterns.

The above analysis examined all synapses incoming to each postsynaptic cell type regardless of presynaptic cell identity. To test for further specificity, we restricted the synapses to include only those from confirmed L2/3 PyCs (Figures 3G and 3H). Chandelier cells and the neurogliaform cell generally received smaller synapses from L2/3 PyCs, whereas bipolar and Martinotti cells generally received larger synapses. This suggests that the biological rules governing synapse size are modulated by both pre- and postsynaptic cell identities.

### Mitochondrial size in neuronal compartments and its relation to synapse density

Regulation of mitochondrion size in neurons affects synaptic plasticity (Divakaruni *et al.*, 2018; Rangaraju, Lauterbach and Schuman, 2019), axonal and dendritic branching (Li *et al.*, 2004; Bertholet et al., 2013; Courchet *et al.*, 2013; López-Doménech *et al.*, 2016; Lewis *et al.*, 2018), and synaptic transmission (Sun *et al.*, 2013; Kwon *et al.*, 2018; Lewis *et al.*, 2018). Our EM reconstruction is suitable for characterizing how the mitochondrial architecture varies between different cell classes and within different compartments of a neuron because it includes large fractions of entire cells, unlike previous EM reconstructions of smaller volumes (Kasthuri *et al.*, 2015; Smith *et al.*, 2016; Bloss *et al.*, 2018; Calì et al., 2018).

It is known that axonal mitochondria are small and punctate, whereas dendritic mitochondria are often much longer and tubular (Chang, Honick and Reynolds, 2006; Lewis *et al.*, 2018). To get more detailed information, we used a semi-automated approach to label each mitochondrion in 125 of our PyC reconstructions by the neuronal compartment it occupies (i.e. axon, soma, apical dendrite, or basal dendrite; STAR Methods). Selecting statistical comparisons by ordered median value (see comparisons in Figure 4D), axonal mitochondria were on average smaller than somatic mitochondria (one-tailed Mann-Whitney U test: p≈0), which were smaller than local apical dendritic mitochondria (p≈0), which were in turn smaller than basal dendritic mitochondria (p<2.8 × 10^−26^). Our ordering is consistent with a previous report suggesting that somatic mitochondria are smaller on average than “neurite” mitochondria as the authors noted that the vast majority of their reconstructed mitochondria were dendritic (Calì *et al.*, 2019).

The fact that somatic mitochondria are statistically intermediate in size relative to axonal and dendritic mitochondria could have multiple causes. A filtering mechanism could prevent trafficking of somatic mitochondria of inappropriate size into the neurites, or mitochondrial size could be adjusted by compartment-specific balance between fission and fusion after trafficking into the neurites (Lewis *et al.*, 2018). The latter idea is consistent with our observation of spatial gradients in mitochondrial size in the perisomatic neurites (Figure S4).

Manipulations of cultured PyCs have shown a coupling between dendritic mitochondrial density and synaptic density (Li *et al.*, 2004; Dickey and Strack, 2011; Bertholet *et al.*, 2013). Such a coupling could be functionally significant for the energetic needs (Harris, Jolivet and Attwell, 2012) or calcium dynamics (Berridge, 1998; Augustine, Santamaria and Tanaka, 2003; Lewis *et al.*, 2018) of synapses. It has remained unclear, however, whether this coupling holds for PyCs under natural conditions. It is also unknown whether the coupling is specific to particular cellular compartments or is cell-wide. We are able to probe this issue because we can compare mitochondria in the soma with those from other compartments of the same cell.

Of the 125 PyCs mentioned above, the somas of 65 PyCs were completely contained in the EM volume. For these 65 cells, we computed mitochondria coverage factors for the basal dendrites (fraction of length covered; Figure S4A and STAR Methods; called mitochondrial index by Li *et al.*, 2004) and the soma (fraction of cytosol volume covered). We also computed synapse densities for the basal dendrites (number of synapses per unit length) and soma (number of synapses per unit area).

We found that mitochondrial coverage is well-correlated with synapse density for basal dendrites (Pearson’s *r*=0.62, p<3.3 × 10^−8^; Figure 4E). Other correlations are weaker, such as basal mitochondrial coverage vs. somatic synapse density (*r*=0.26, p=0.036; Figure 4F); basal vs. somatic mitochondrial coverage (*r*=0.20, p=0.104; Figure 4G); and somatic mitochondrial coverage vs. synapse density (*r*=0.34, p=0.005; Figure 4H). Taken together, these results suggest that the co-regulation of mitochondrial coverage and synapse density exhibits some compartment specificity.

### Predicting motif frequencies from degree sequence

Multi-electrode synaptic physiology studies in brain slices have established that bidirectional connections between cortical pyramidal cells are “overrepresented,” i.e., more frequent than would be expected by chance (Markram *et al.*, 1997; Le Bé and Markram, 2006; Wang *et al.*, 2006). Similar reports have been made for certain three-cell connectivity motifs (Song *et al.*, 2005). Both kinds of overrepresentation have been interpreted as evidence for cortical circuit theories, such as Hebbian cell assemblies (Perin, Berger and Markram, 2011), neuron clustering (Klinshov *et al.*, 2014), and attractor networks (Brunel, 2016; Zhang, Zhang and Stepanyants, 2019). A related overrepresentation was also reported for cerebellum (Rieubland, Roth and Häusser, 2014). The large circuit of connected PyCs within this reconstruction provides an opportunity to reexamine claims of motif overrepresentation. Rather than many small circuits of a handful of PyCs mapped using brain slice physiology, we have a single large circuit mapped using EM. We excluded all PyCs with total axonal length less than 100 μm; these were too cut off by the borders of the EM volume to have a chance of making outgoing connections. This left a total of 113 PyCs in the graph to be analyzed.

We first examined two-cell motifs and found that bidirectionally connected pairs are about 1.66 times more frequent (Figure 5E) than predicted by an Erdős-Renyi (ER) model, where connections are randomized among cells with a single probability value. This is qualitatively consistent with many previous brain slice studies (Markram *et al.*, 1997; Song *et al.*, 2005; Le Bé and Markram, 2006; Wang et al., 2006).

We decided to explore an alternative model of random graphs, the configuration model, which has been applied to many biological networks including the *C. elegans* connectome (Milo *et al.*, 2002). We became interested in the configuration model because the node degrees in our PyC graph exhibit more variation than in random graphs drawn from the ER model (Figure 5D). Here we define the in-degree of a cell by the number of its presynaptic partner cells in our 113-node graph, and the out-degree of a cell by the number of its postsynaptic partner cells. The configuration model constrains the degree sequence of random graphs to exactly match the observed graph. We hypothesized that the degree constraint alone would lead to more accurate predictions of motif frequencies, as previously proposed but never empirically tested in the cortex (Song *et al.*, 2005; Gal *et al.*, 2017; Vegué, Perin and Roxin, 2017).

We sampled from the set of graphs with the same degree sequence as our cortical circuit by applying degree-preserving edge swaps (Figure 5A), including the “hold” technique to ensure random sampling of graphs with uniform probability (Artzy-Randrup and Stone, 2005). The equilibrium distribution of this sampling procedure is known to yield correct motif frequencies (Berger and Müller-Hannemann, 2010), STAR Methods). We found that bidirectional connections are about 1.38 times more frequent than the counts predicted by the configuration model (Figure 5E). The overrepresentation is still statistically significant, but is less than the overrepresentation observed relative to the ER model (Figure 5F and Table S1; configuration model p=0.025, ER model p<0.001).

Moving on to three-cell motifs, we see that highly connected motifs (numbered 10-16 in Figure 5G) are substantially overrepresented relative to the ER model (Figure 5G), qualitatively consistent with previous findings (Song *et al.*, 2005; Perin, Berger and Markram, 2011). Here we use a generalization of the ER model that preserves frequencies of bidirectional connections (Figure 5B; Song *et al.*, 2005). The overrepresentations are reduced, however, if the comparison is relative to the configuration model (Figure 5G and Table S2). A scatter plot compares model predictions of three-cell motif frequencies, and the configuration model is visibly more accurate than the generalized ER model (Figure 5H).

The statistics of three-cell motifs are related to network clustering, or transitivity (Holland and Leinhardt, 1971). In a network with high clustering, if *α* is connected to *β*, and *β* is connected to *γ*, then *α* and *γ* are likely connected. This is quantified by the clustering coefficient:

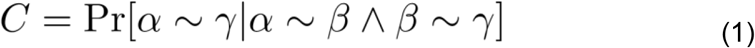

where *α*~*β* means that *α* is connected to *β*, and where the nodes (*α*, *β*, *γ*) are drawn randomly from the network (Figure 5C). It has been claimed that cortical networks are highly clustered, and that such clustering is evidence of Hebbian cell assemblies (Perin, Berger and Markram, 2011). We can check this claim by rewriting the clustering coefficient as

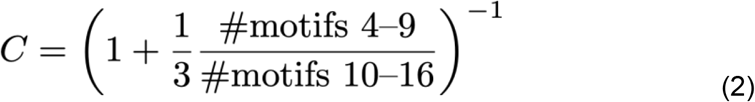

which is a function of the relative frequency of motifs containing two connected pairs (4-9) versus those containing three connected pairs (10-16). The observed clustering coefficient (*C*_OBS_ = 0.161; Table 1) is much higher than that predicted by the generalized ER model (*C*_ER_ = 0.101) but only marginally higher than predicted by the configuration model (*C*_CFG_ = 0.150). Therefore claims about high clustering in cortical networks should be taken with caution.

**Table 1:** Clustering coefficients of all versions of PyC subgraph and null models. Numbers are computed analytically for ER and generalized ER models, and empirically for the configuration model and generalized configuration model by averaging over 1,000 samples. The p-value is computed as the percentile of the observed clustering coefficient in the distribution of clustering coefficient sampled from the configuration model.

|  | OBS | ER | Generalized ER | CFG | Generalized CFG | p-value (OBS vs CFG) |
| --- | --- | --- | --- | --- | --- | --- |
| **Full Graph** | 0.09511 | 0.02646 | 0.026163 | 0.085065 | 0.083758 | 0.000 |
| **> 0 $\mu\text{m}$ axon** | 0.11788 | 0.04783 | 0.04723 | 0.10755 | 0.1059 | 0.005 |
| **> 100 $\mu\text{m}$ axon** | 0.16128 | 0.10248 | 0.10066 | 0.14957 | 0.14626 | 0.027 |
| **> 300 $\mu\text{m}$ axon** | 0.16865 | 0.12171 | 0.1198 | 0.15744 | 0.15416 | 0.040 |

Moving beyond two-cell and three-cell motifs, it has been argued that the connection probability for a pair of cortical pyramidal cells increases with the number of common synaptic partners (Perin, Berger and Markram, 2011). This “common neighbor rule” is a generalization of the idea of high network clustering (Perin, Berger and Markram, 2011), and is inconsistent with the ER model, which predicts that connection probability does not vary with the number of common neighbors. For our reconstructed circuit, the common neighbor rule is largely predicted by the configuration model (Figure 5I). A similar approximate agreement was also reported in *C. elegans* (Azulay, Itskovits and Zaslaver, 2016).

The above analysis excluded PyCs with axon length less than 100 μm. Analyses for the graph consisting of all 363 PyCs, and subgraphs with other threshold values for axon length, are shown in Data S4 and Figure S5. In general, we find that increasing the axon length threshold improves the fit to observed motif frequencies for both models (note the vertical scale in plots), and also reduces differences between the models. Comparisons with other model variants are given in Table 1, Data S5, and Table S2. In particular, the fit was not improved by generalizing the configuration model to preserve frequencies of bidirectional connections.

Given that motif frequencies, the clustering coefficient, and the common neighbor rule are predicted quite well from degree sequence by the configuration model, interpreting these statistics as evidence of synaptic clustering and Hebbian cell assemblies seems more subtle than has commonly been supposed (see Discussion).

### Relation of in-connection density to visual function

Using our PyC graph, we showed above that the frequencies of certain connectivity motifs can be predicted quite accurately from the degree sequence. For 112 of these same PyCs, calcium imaging data are available (Figures 6A and 6C), opening the possibility of relating degree to visual function. We define the in-connection density of a node in the PyC graph as in-degree (number of presynaptic partners in the PyC graph) divided by the total dendritic length of the node (i.e. postsynaptic neuron). This normalized quantity indicates a cell’s tendency to receive connections from nearby PyCs, relative to a baseline predicted from its total dendritic length in the EM volume (Figure S6). Some of the variation in total dendritic length arises from the varying extent to which arbors are cut off by the borders of the EM volume.

In-connection density quantifies local recurrent excitation, since it involves only nearby PyCs. When L4 cortical cells are activated by visual stimuli, their responses are thought to be amplified by local recurrent excitation from neighboring cortical cells (Douglas *et al.*, 1995; Lien and Scanziani, 2013; Li *et al.*, 2013). A similar amplification may also happen in L2/3 (Cossell *et al.*, 2015). We therefore hypothesized that L2/3 PyCs with higher in-connection density would exhibit stronger responses to visual stimuli.

For each calcium-imaged cell, we extracted a direction tuning curve and computed a direction selectivity index (DSi) and an orientation selectivity index (OSi) (Methods). Since we are considering the responses to directional stimuli (Figure 6B), further analysis was restricted to cells with statistically significant DSi or OSi values as evaluated using a permutation test (Ecker *et al.*, 2014; Baden *et al.*, 2016). As hypothesized, we found that the response of a cell to its preferred direction or orientation averaged across trials (trial-averaged response) tended to be larger for cells with higher in-connection density (Pearson’s *r*=0.46, p=0.005; Figure 6F).

We further hypothesized that L2/3 PyCs with higher in-connection density would exhibit more reliable responses to visual stimuli. Trial-to-trial variability (Stein, Gossen and Jones, 2005) is illustrated for example cells at the bottom of Figure 6E. Cells 2 and 3 exhibited little or no response for many trials. Nevertheless, the cells responded strongly to their preferred stimuli in several trials and did not respond to non-preferred stimuli. Accordingly, their DSi and OSi values were statistically significant. The examples show that a cell can be well-tuned for direction or orientation and yet respond only intermittently. Note that defining tuned cells by statistical significance excludes cells that have a large DSi or OSi merely by chance, and is different from imposing a simple threshold on DSi or OSi (Baden *et al.*, 2016).

For each trial, we thresholded the response of a neuron to its preferred stimulus (STAR Methods). The threshold was set at three standard deviations above the baseline (Figures 6D). A cell’s *intermittency* index was defined as the fraction of trials for which a cell’s response did not exceed the threshold. Most cells have intermittency values exceeding 50% (Figure 6E). These values are likely overestimates, since some spikes may have evaded detection (Huang *et al.*, 2020), so we focused our analysis on relative differences between cells, rather than absolute intermittency values.

We indeed found that in-connection density is negatively correlated with intermittency (Pearson’s *r*=−0.49, p=0.003; Figure 6H). We also found that intermittency is not correlated with “total in-synapse density”, i.e. the number of incoming synapses from all sources (with or without somas in the volume) normalized by total dendritic length (Pearson’s *r*=0.05, p=0.757; Figure 6I).

Other things being equal, more intermittent cells are expected to have lower trial-averaged responses. To try to separate the effects of these two variables, we fit a linear model of the form:

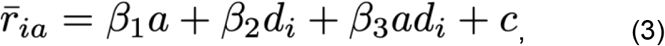

where *a* is a binary variable that captures whether or not responses surpassed the intermittency threshold, *d*_*i*_ represents the in-connection density for cell *i*, and 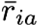 is the mean response for cell given *a*. We find a significant interaction term *β*_3_ (p=0.006; Figure 6G), suggesting a relationship between in-connection density and responses over the activity threshold.

In-connection density is weakly correlated with cortical depth, presumably because our normalization by dendritic length does not compensate for all effects of neuronal geometry (Figures S7A-C). Therefore we investigated the possibility that cortical depth or more generally 3D location could be a confounding variable in the observed correlations of mean response and intermittency with in-connection density. For a permutation test, we computed the same correlations after shuffling cells with similar 3D locations in the volume (Figures S7D and S7E). The observed correlations in our data were still stronger than in the permutations (mean response p=0.0178, intermittency p=0.018; Figure 6J). Taken together, these results suggest that cells with higher in-connection density tend to be less intermittent in their responses and exhibit stronger mean responses.

## Discussion

The current EM reconstruction is large enough to enable analysis of three-cell motifs, including the highly connected ones (numbered 10-16 in Figure 5G). This is a milestone in scale for cortical connectomics. That being said, motif statistics are still affected by truncation of neuronal arbors, especially axons. To probe the effects of truncation, we have applied our motif analysis to the full PyC graph (Data S4) and also the subgraphs obtained by excluding PyCs with highly truncated axons (Figure 5 and Figure S5). For all graphs, the motif counts, the clustering coefficient, and the common neighbor rule are all fairly well-predicted by the configuration model. These statistics deviate strongly from the ER model for the full PyC graph, but not as strongly for the subgraphs of less truncated PyCs. The theoretical possibility that nonuniform degree could explain large deviations from the ER model was noted previously (Song *et al.*, 2005; Gal *et al.*, 2017; Vegué, Perin and Roxin, 2017). We have explored this possibility using empirical data from the cortex.

One should be cautious about comparing the current findings based on EM reconstruction with older work based on synaptic physiology in brain slices (Song *et al.*, 2005; Perin, Berger and Markram, 2011). Both methods truncate neuronal arbors, but the geometry of truncation differs. While brain slices are thicker than our reconstructed volume, neurons recorded in brain slices can be affected by truncation because they are close to the surface of the slice, typically within tens of microns. Forthcoming EM reconstructions of much larger volumes (Yin *et al.*, 2019) should make it possible to eliminate the effects of truncation from motif statistics.

A major motivation for studying motif counts and other cortical connectivity statistics is to constrain cortical theories. Our work suggests a number of ways to approach such constraints. Given that motif statistics are predicted quite well by the configuration model, one approach is to extract predictions about degree sequence from cortical theories and compare these predictions with reconstructed circuits. For example, (Brunel, 2016) theoretically analyzed attractor models of associative memory to predict that cortical neurons should have broad out-degree but not in-degree distributions. Alternatively, a long-tailed degree distribution may be a consequence of neuronal geometry (Gal *et al.*, 2017), which by itself has been shown to yield large deviations of motif frequencies from the ER model (Gal *et al.*, 2020).

Another theoretical approach would be to interpret those remaining deviations from the configuration model that are statistically significant. The overrepresentation of bidirectional connections is statistically significant for the full PyC graph (Data S4B) and for the subgraphs of less truncated PyCs (Figure 5E and Figures S5F-G). For the doubly- and triply-connected motifs (green and red boxes in Figure 5G), there are some statistically significant deviations from the configuration model in Figure 5G. None of these deviations, however, remain significant for a smaller subgraph created with a larger axonal length threshold (Figure S5G and Figure S5I). We also note that we have not applied any correction for testing multiple hypotheses. The clustering coefficient, an overall measure depending on the ratio of triply-connected to doubly-connected motifs (Eq. 2), is slightly larger than predicted by the configuration model, and this deviation is statistically significant for all subgraph versions analyzed (Table 1). It would be interesting to explore the implications of these deviations from the configuration model for cortical theories.

More generally, connectomics opens the possibility of moving beyond statistics of few-cell motifs towards more global models of graphs (Seung, 2009). The configuration model can be viewed as a step in this direction. Many other models of random graphs have been proposed in the literature, and could also be applied to our PyC graph. Such modeling can be carried out by anyone, as the PyC graph has already been made publicly available as part of the resource.

We defined the in-connection density of a PyC as the ratio of in-degree to total dendritic length. This normalized quantity is intended to measure the cell’s preference for connecting with nearby cells, after compensating for truncation. We found that in-connection density is correlated with the intermittency and amplitude of visual response. This suggests that at least some of the variation in degree is related to visual function, and due to biological variations in local connectivity rather than merely truncation artifacts. Future reconstructions of larger cortical volumes will be important for distinguishing biological variation of degree versus truncation artifacts.

We found that basket and chandelier cells exhibited almost identical areal synapse densities in their somas and dendrites (Figure 3D). This commonality may follow from the overlap of both anatomical classes with the molecularly defined class of fast-spiking, parvalbumin-positive interneurons, yet verifying this connection is beyond the scope of our current methods. Our finding of high synapse density on perisomatic dendrites of basket and chandelier cells has a number of possible implications for dendritic biophysics. For example, high input density near the soma could work in conjunction with specialized ion channel distributions (Hu, Martina and Jonas, 2010) and permissive cable properties (Norenberg *et al.*, 2010) to evoke rapid responses to network activity through dense sampling of the local network (Hu, Gan and Jonas, 2014).

We found a correlation between mitochondrial coverage and synapse density that is stronger for basal dendrites than for somas (Figure 4E and 4H). Synapses are important sites of Ca^2+^ signaling in neurons (Berridge, 1998; Augustine, Santamaria and Tanaka, 2003), and also consume a significant amount of ATP (Harris, Jolivet and Attwell, 2012). Mitochondria can rapidly uptake Ca^2+^, and therefore buffer cytoplasmic Ca^2+^, both in dendrites, where they can regulate synaptically-evoked cytoplasmic (MacAskill, Atkin and Kittler, 2010; Kwon *et al.*, 2016) dynamics (Hirabayashi *et al.*, 2017) and synaptic plasticity (Divakaruni *et al.*, 2018), and in axons where they are enriched at presynaptic release sites and regulate synaptic release properties (MacAskill, Atkin and Kittler, 2010; Kwon *et al.*, 2016). A weaker correlation between mitochondria coverage and synaptic density in the soma compared to basal dendrites also might reflect a difference between the largely excitatory axo-dendritic synapses and inhibitory axo-somatic synapses on these systems (Harris, Jolivet and Attwell, 2012). Alternatively, other mitochondrial functions in neurons might affect or determine mitochondrial morphology in dendrites and somas, such as coupling between ATP synthesis and protein translation (Rangaraju, Lauterbach and Schuman, 2019) or reactive oxygen species signaling (Fu *et al.*, 2017). Our analysis of compartment-specificity is a first step, and several more steps are possible using the same data. Coupling functional investigations with a more spatially precise study of synapse and mitochondrion colocalization in specific neuronal subtypes is made possible by our combined reconstruction.

Because the reconstruction is multimodal, combining both electron microscopy and calcium imaging (Figure 6A), it can be used to relate cortical structure and function, as in our relation of intermittency and amplitude of visual response to in-connection density. Our reconstruction is also multiscale, because it spans both the large-scale (up to 400 μm) structure of neurons and small structures like synapses and organelles. The combination of multiple scales was essential for quantifying connectivity motifs, revealing spatial organization of synapses onto inhibitory cells, and quantitatively describing mitochondria across neuronal arbors. These applications demonstrate that our resource will be widely useful for investigating a broad range of questions about cortical structure and function.

## Acknowledgements

Supported by the Intelligence Advanced Research Projects Activity (IARPA) via Department of Interior/ Interior Business Center (DoI/IBC) contract numbers D16PC00003, D16PC00004, and D16PC0005. The U.S. Government is authorized to reproduce and distribute reprints for Governmental purposes notwithstanding any copyright annotation thereon. HSS also acknowledges support from NIH/NINDS U19 NS104648, ARO W911NF-12-1-0594, NIH/NEI R01 EY027036, NIH/NIMH U01 MH114824, NIH/NINDS R01NS104926, NIH/NIMH RF1MH117815, and the Mathers Foundation, as well as assistance from Google, Amazon, and Intel. FP acknowledges support from NIH/NINDS RO1 NS107483. We thank S. Koolman, M. Moore, S. Morejohn, B. Silverman, K. Willie, and R. Willie for their image analyses, G. McGrath for computer system administration, and M. Husseini and L. and J. Jackel for project administration. We are grateful to J. Maitin-Shepard for making neuroglancer freely available. We thank M. Tsodyks, D. Tank, C. Brody, V. Hewitt, A. Wanner, and S. Papadopoulos for helpful discussions and feedback, and A. Foryciarz for preliminary mitochondria analysis. We thank the Allen Institute for Brain Science founder, Paul G. Allen, for his vision, encouragement and support. Disclaimer: The views and conclusions contained herein are those of the authors and should not be interpreted as necessarily representing the official policies or endorsements, either expressed or implied, of IARPA, DoI/IBC, or the U.S. Government.

## Author Contributions

EC built calcium imaging pipeline and EF, JR, and AST performed and oversaw surgery and calcium imaging. ALB, AAB, DB, DJB, JB, and NMC generated the EM dataset after sample preparation by JB, MT, and NMC. RT, GM, and YL stitched and rough aligned block one of the EM images (STAR Methods). DI and TM stitched and rough, coarse, & fine aligned block two of the images. TM coarse & fine aligned all sections. WW built the task manager for distributed alignment. WMS wrote the software for reading and writing data in cloud storage, initially with IT. TM supervised ground truth annotations. SP implemented a framework for CPU-based 3D convolutional net inference with help from AZ. The net for boundary detection was trained by KL with help from JZ, applied by JW, and segmented by RL with help from AZ. The nets for synapse and mitochondria detection were trained by NLT, applied by JW, and segmented by NLT. The net for synaptic partner assignment was trained and applied by NLT. SM trained and applied the net for nucleus detection with ground truth partly generated by LE and FC. SD, NK, and JZ created the proofreading system. SCY, TM, SD, and AMW supervised proofreading and annotation. FC, ALB, NMC, SCY, SD, AMW, and CSM contributed proofreading and annotations. AMW manually verified nucleus detections in the volume and performed cell typing with JB, NMC, and ALB. NK and MC built the front end for proofreading and annotations and generated neuron meshes with WMS. CSJ built the early data sharing system. SD, FC, and CSM created the connectome versioning system. FC, WMS, and CSM skeletonized cells with help from JAB, NLT, and JW. NK and AMW rendered example cells. AMW and NLT compiled resource statistics. CSM analyzed synapses onto inhibitory cells with input from FC, LE and NMC. NLT analyzed mitochondria with help from AMW, FP, CSM, FC, and RY. TM and RY analyzed motifs in the PyC graph with input from NLT. JAB co-registered calcium and EM images using correspondences annotated by NMC, AAB, and JR. JAB extracted activity traces using the approach developed by PZ and LP, and related these traces to in-degree with input from JR, AST, FC, and NMC. FC created the microns-explorer web site with contributions from CSM and NMC. HSS, NLT, AMW, JAB, CSM, and RY wrote the paper, with contributions from FC, NK, MT, NMC, FP, and LP. SS and LB managed the project at the Allen Institute. TM managed the reconstruction team. HSS, RCR, NMC, JR, and AST managed the multi-institution collaboration.

## STAR ⋆ METHODS

### METHOD DETAILS

#### Mouse

All procedures were in accordance with the Institutional Animal Care and Use Committees at the Baylor College of Medicine and the Allen Institute for Brain Science.

#### Mouse line

Functional imaging was performed in a transgenic mouse expressing fluorescent GCaMP6f. For this data set, the mouse we used was a triple-heterozygote for the following three genes: (1) Cre driver: CamKIIa-Cre (Jax: 005359<https://www.jax.org/strain/005359>), (2) tTA driver: B6;CBA-Tg(Camk2a-tTA)1Mmay/J (Jax: 003010<https://www.jax.org/strain/003010>), (3) GCaMP6f Reporter: Ai93 (Allen Institute).

#### Cranial window surgery

Anesthesia was induced with 3% isoflurane and maintained with 1.5% to 2% isoflurane during the surgical procedure. Mice were injected with 5-10 mg/kg ketoprofen subcutaneously at the start of the surgery. Anesthetized mice were placed in a stereotaxic head holder (Kopf Instruments) and their body temperature was maintained at 37°C throughout the surgery using a homeothermic blanket system (Harvard Instruments). After shaving the scalp, bupivicane (0.05 cc, 0.5%, Marcaine) was applied subcutaneously, and after 10-20 minutes an approximately 1 cm^2^ area of skin was removed above the skull and the underlying fascia was scraped and removed. The wound margins were sealed with a thin layer of surgical glue (VetBond, 3M), and a 13mm stainless-steel washer clamped in the headbar was attached with dental cement (Dentsply Grip Cement). At this point, the mouse was removed from the stereotax and the skull was held stationary on a small platform by means of the newly attached headbar. Using a surgical drill and HP 1/2 burr, a 3 mm craniotomy was made centered on the primary visual cortex (V1; 2.7mm lateral of the midline, contacting the lambda suture), and the exposed cortex was washed with ACSF (125mM NaCl, 5mM KCl, 10mM Glucose, 10mM HEPES, 2mM CaCl_2_, 2mM MgSO_4_). The cortical window was then sealed with a 3 mm coverslip (Warner Instruments), using cyanoacrylate glue (VetBond). The mouse was allowed to recover for 1-2 hours prior to the imaging session. After imaging, the washer was released from the headbar and the mouse was returned to the home cage.

#### Widefield imaging

Prior to two-photon imaging, we acquired a low-magnification image of the 3mm craniotomy under standard illumination.

#### Two-Photon imaging

Imaging for candidate mice was performed in V1, in a 400 × 400 × 200 μm^3^ volume with the superficial surface of the volume at the border of L1 and L2/3, approximately 100um below the pia. Laser excitation was at 920nm at 25-45mW depending on depth. The objective used was a 25× Nikon objective with a numerical aperture of 1.1, and the imaging point-spread function was measured with 500 nm beads and was approximately 0.5 × 0.5 × 3 μm^3^ in x, y, and z. Pixel dimensions of each imaging frame were 256 × 256.

#### Functional imaging scans

Imaging data was collected with a resonant scanning microscope (ThorLabs) and software (Scanimage 5.1, Vidrio). Nine scans were collected in total, starting superficially and moving deeper into the cortex with each subsequent scan. During each 30-minute scan, a piezo controlled manipulator (PI-726, Physik Instruments) moved the microscope objective between three different z-planes (“focal planes”). These three focal planes were separated by an average of about 8 μm by the piezo, and each focal plane was imaged at 14.8313 frames per second. We refer to each trio of sequential focal plane as one imaging “volume”. Since we collected 9 volumes with 3 focal planes/volume, we had 27 focal planes in total to span the 200 μm depth of the overall 400×400×200 μm^3^ imaging volume. Two color channels were recorded: Channel one was GCaMP6 calcium imaging and channel two was blood vessels labeled with red dye (Sulfarhodamine 101). After the data was collected, we learned that the piezo command functionality of ScanImage 5.1 was not fully optimized, so the movement of the objective between focal planes was slower than expected. As a result, the posterior edge of the field of view of each focal plane was curved in the Z dimension by several microns. This curvature was a deterministic function of the piezo command and can also be recovered from the high resolution (1 μm spacing) structural stack collected with each functional scan, and we do not expect it to greatly impact the alignment with the EM data. All scans were raster- and motion-corrected. Motion correction in X and Y was performed by sub-pixel cross-correlation. Focal plane alignment into the structural stack was performed by 3D cross-correlation.

#### High resolution structural stack

To facilitate alignment with EM, at the beginning of the experiment we collected a high-resolution structural stack of the imaging volume with the same field of view and (x,y) location as the functional scans. This stack began 310 μm deep and ended at the cortical surface, in one micron steps.

#### Behavioral monitoring

The mouse was head-restrained but could walk on a treadmill during imaging. While we were imaging, we collected treadmill speed at 200 Hz and we recorded a movie of the mouse’s eye at 20 Hz. Pupil size and (x,y) position were extracted from this movie using custom MATLAB and Python code. Both treadmill traces and pupil traces were subsequently synchronized and interpolated to the imaging clock (27300 samples synchronized to the first focal plane of each imaging volume).

#### Visual stimulus

Visual stimuli were presented at 60 fps, and synchronized to imaging and behavioral data via a photodiode which recorded the timing of each stimulus frame. For each 30-minute 40-second scan (27300 volumes at 14.8313 volumes per second) we presented 30 one-minute trials of a colored-noise stimulus (Niell and Stryker, 2008) interspersed with periods of coherent motion of oriented noise. Each one-minute trial contained 16 stationary moving stationary blocks, with a different direction presented in each block, pseudo-randomly ordered. The stimulus was interpolated and aligned to the calcium imaging frame times and stored as a 90×160×27300 matrix of images. First 200 frames were black screen and were used for extracting calcium traces but were not used for the analysis.

#### Calcium trace extraction and deconvolution

Calcium traces have been extracted with the EASE algorithm (Zhou *et al.*, 2020). All the traces used in the analysis were signals from the somas. To extract the signals from somas, a corresponding scan for each cell has been identified by finding the closest scan from the soma location. EASE algorithm uses reconstructed cells for initialization so cells with severely cut-off somas were excluded from the analysis (*n*=3). Also, calcium traces were extracted only for a subset of reconstructed cells that overlap with the calcium recording volume as EASE uses reconstructed cells for initialization. Activity traces of cells that have low signal-to-noise ratio were neglected from the extraction. Spike inference was performed with the modified OASIS algorithm (Friedrich, Zhou and Paninski, 2017), which is implemented within the EASE package.

#### Co-registration of EM and high-resolution 2-photon structural stack

Centers of cell bodies have been annotated manually for both the EM and 2-photon high resolution stacks. A total of 512 points have been labeled in each dataset and the corresponding cells have been determined manually by comparing the relative locations of the cell bodies in both datasets. Based on this list of 512 correspondence points, an affine transformation was estimated and this transformation was applied to the entire EM volume to coregister the volume onto the high-resolution 2-photon structural stack.

#### Tissue preparation and staining

The protocol of Hua et al. (Hua, Laserstein and Helmstaedter, 2015) was combined with the protocol of Tapia et al. (Tapia *et al.*, 2012) to accommodate a smaller tissue size and to improve TEM contrast. Mice were transcardially perfused with 2.5% paraformaldehyde and 1.25% glutaraldehyde. After dissection, 200 μm-thick coronal slices were cut with a vibratome and post-fixed for 12-48 hours. Following several washes in 0.1 M cacodylate buffer, pH 7.4 (CB), the slices were fixed with 2% osmium tetroxide in CB for 90 minutes, immersed in 2.5% potassium ferricyanide in CB for 90 minutes, washed with deionized (DI) water for 2 × 30 minutes, and treated with freshly made and filtered 1% aqueous thiocarbohydrazide at 40°C for 10 minutes. The slices were washed 2 × 30 minutes with DI water and treated again with 2% osmium tetroxide in water for 30 minutes. Double washes in DI water for 30 minutes each were followed by immersion in 1% aqueous uranyl acetate overnight at 4°C. The next morning, the slices in the same solution were placed in a heat block to raise the temperature to 50°C for 2 hours. The slices were washed twice in DI water for 30 minutes each, and then incubated in Walton’s lead aspartate, pH 5.0, for 2 hours at 50°C in the heat block. After another double wash in DI water for 30 minutes each, the slices were dehydrated in an ascending ethanol series (50%, 70%, 90%, 100% × 3) for 10 minutes each and two transition fluid steps of 100% acetonitrile for 20 minutes each. Infiltration with acetonitrile:resin dilutions (2p:1p, 1p:1p and 2p:1p) were performed on a gyratory shaker overnight for 4 days. Slices were placed in 100% resin for 24 hours followed by embedding in Hard Plus resin (EMS, Hatfield, PA). Slices were cured in a 60°C oven for 96 hours. The best slice based on tissue quality and overlap with the 2-photon region was selected.

#### Sectioning and collection

A Leica EM UC7 ultramicrotome and a Diatome 35-degree diamond ultra-knife were used for sectioning at a speed of 0.3 mm/sec. Eight to ten serial sections were cut at 40 −nm thickness to form a ribbon, after which the microtome thickness setting was set to 0 in order to release the ribbon from the knife. Using an eyelash probe, pairs of ribbons were collected onto copper grids covered by 50-nm thick LUXEL film.

#### Transmission electron microscopy

We made several custom modifications to a JEOL-1200EXII 120-kV transmission electron microscope (Yin *et al.*, 2019). A column extension and scintillator magnified the nominal field of view by tenfold with negligible loss of resolution. A high-resolution, large-format camera allowed fields-of-view as large as (13 μm)^2^ at 3.58-nm resolution. Magnification reduced the electron density at the phosphor, so a high-sensitivity sCMOS camera was selected and the scintillator composition tuned to generate high quality EM images with exposure times of 90-200 ms. Sections were acquired as a grid of 3840 × 3840-px^2^ images (“tiles”) with 15% overlap.

#### Alignment in two blocks

The dataset was divided by sections into two blocks (1216 & 970 sections), with the first block containing substantially more folds. Initial alignment and reconstruction tests proceeded on the second block of the dataset. After achieving satisfactory results, the first block was added, and the whole dataset was further aligned to produce the final 3D image. The alignment process included stitching (assembling all tiles into a single image per section), rough alignment (aligning the set of section images with one affine per section), coarse alignment (nonlinear alignment on lower resolution data), and fine alignment (nonlinear alignment on higher resolution data).

#### Alignment, block one

The tiles of the first block were stitched into one montaged image per section and roughly aligned using a set of customized and automated modules based on the “TrakEM2” (Cardona *et al.*, 2012) and “Render” (Zheng *et al.*, 2018) software packages.

##### Stitching

After acquisition, a multiplicative intensity correction based on average pixel intensity was applied to the images followed by a lens distortion of individual tiles using nonlinear transformations (Kaynig *et al.*, 2010). Once these corrections were applied, correspondences between tiles within a section were computed using SIFT features, and each tile was modeled with a rigid transform.

##### Rough alignment

Using 20x downsampled stitched images, neighboring sections were roughly aligned (Saalfeld *et al.*, 2012). Correspondences were again computed using SIFT features, and each section was modeled with a regularized affine transform (90% affine + 10% rigid), and all correspondences and constraints were used to generate the final model of one affine transform per tile. These models were used to render the final stitched section image into rough alignment with block two.

#### Alignment, block two

The second block was stitched and aligned using the methods of (Saalfeld *et al.*, 2012) as implemented in Alembic (Macrina and Ih, no date).

##### Stitching

For each section, tiles containing tissue without clear image defects were contrast normalized by centering the intensities at the same location in each tile, stretching the overall distribution between the 5th & 95th intensity percentiles. During imaging, a 20x downsampled overview image of the section was also acquired. Each tile was first placed according to stage coordinates, approximately translated based on normalized cross-correlation (NCC) with the overview image, and then finely translated based on NCC with neighboring tiles. Block matching was performed in the regions of overlap between tiles using NCC with 140-px block radius, 400-px search radius, and a spacing of 200 px. Matches were manually inspected with 1x coverage, setting per-tile-pair thresholds for peak of match correlogram, distance between first and second peaks of match correlograms, and correlogram covariance, and less frequently, targeted match removal. A graphical user interface was developed to allow the operator to fine-tune parameters on a section-by-section basis, so that a skilled operator completed inspection in 40 hours. Each tile was modeled as a spring mesh, with nodes located at the center of each blockmatch operation, spring constants 1/100th of the constant for the between-tile springs, and the energy of all spring meshes within a section were minimized to a fractional tolerance of 10^−8^ using nonlinear conjugate gradient. The final render used a piecewise affine model defined by the mesh before and after relaxation, and maximum-intensity blending.

##### Rough alignment

Using 20x downsampled images, block matching between neighboring sections proceeded using NCC with 50-px block radius, 125-px search radius, and 250-px spacing. Matches were computed between nearest neighbor section pairs, then filtered manually in 8 hours. Correspondences were used to develop a regularized affine model per section (90% affine + 10% rigid), which was rendered at full image resolution.

##### Coarse alignment

Using 4x downsampled images, NCC-based block matching proceeded with 300-px block radius, 200-px search radius, and 500-px spacing. Matches were computed between nearest and next-nearest section pairs, then manually filtered by a skilled operator in 24 hours. Each section was modeled as a spring mesh with spring constants 1/10th of the constant for the between-section springs, and the energy of all spring meshes within the block were minimized to a fractional tolerance of 10^−8^ using nonlinear conjugate gradient. The final render used a piecewise affine model defined by the mesh.

##### Fine alignment

Using 2x downsampled images, NCC-based block matching proceeded with 200-px block radius, 113-px search radius, and 100-px spacing. Matches were computed between nearest and next-nearest section pairs, then manually filtered by a skilled operator in 24 hrs. Modeling and rendering proceeded as with coarse alignment using spring constants, which were 1/20th of the constant for the between-section springs.

#### Alignment, whole dataset

Blank sections were inserted manually between sections where the cutting thickness appeared larger than normal (11). The alignment of the whole dataset was further refined using the methods of (Saalfeld *et al.*, 2012) as implemented in Alembic (Macrina and Ih, no date).

##### Coarse alignment

Using 64x downsampled images, NCC-based block matching proceeded with 128-px block radius, 512-px search radius, and 128-px spacing. Matches were computed between neighboring and next nearest neighboring sections, as well as 24 manually identified section pairs with greater separation, then manually inspected in 70 hours. Section spring meshes had spring constants 1/20th of the constant for the between-section springs. Mesh relaxation was completed in blocks of 15 sections, 5 of which were overlapping with the previous block (2 sections fixed), each block relaxing to a fractional tolerance of 10^−8^. Rendering proceeded similarly to as before.

##### Fine alignment

Using 4x downsampled images, NCC-based block matching proceeded with 128-px block radius, 512-px search radius, and 128-px spacing. Matches were computed between the same section pairs as in coarse alignment. Matches were excluded only by heuristics. Modeling and rendering proceeded similarly to coarse alignment, with spring constants 1/100th the constant for the between-section springs. Rendered image intensities were linearly rescaled in each section based on the 5th and 95th percentile pixel values.

#### Image volume estimation

The imaged tissue has a trapezoidal shape in the sectioning plane. Landmark points were placed in the aligned images to measure this shape. We report cuboid dimensions for simplicity and comparison using the trapezoid midsegment length. The original trapezoid has a short base length of 216.9 μm, long base length of 286.2 μm, and height of 138.3 μm. The imaged data has 2176 sections, which measure 87.04 μm with a 40-nm slice thickness.

#### Image defect handling

Cracks, folds, and contaminants were manually annotated as binary masks on 256x downsampled images, dilated by 2 px, then inverted to form a defect mask. A tissue mask was created using nonzero pixels in the 256x downsampled image, then eroded by 2 px to exclude misalignments at the edge of the image. The image mask is the union of the tissue and defect masks, and it was upsampled and applied during the final render to set pixels not included in the mask to zero. We created a segmentation mask by excluding voxels that had been excluded by the image mask for three consecutive sections. The segmentation mask was applied after affinity prediction to set affinities not included in the mask to zero.

#### Affinity prediction

Human experts used <u>VAST</u> (Berger, Seung and Lichtman, 2018) to manually segment multiple subvolumes from the current dataset and a similar dataset from mouse V1. Annotated voxels totaled 1.29 billion at full image resolution.

We trained a 3D convolutional network to generate three nearest neighbors (Turaga *et al.*, 2010) and 13 long-range affinity maps (Lee *et al.*, 2017). Each long-range affinity map was constructed by comparing an equivalence relation (Jain, Sebastian Seung and Turaga, 2010) of pairs of voxels spanned by an “offset” edge (to preceding voxels at distances of 4, 8, 12, and 16 in x and y, and 2, 3, 4 in z). Only the nearest neighbor affinities were used beyond inference time; long-range affinities were used solely for training. The network architecture was modified from the “Residual Symmetric U-Net” of Lee et al. (Lee *et al.*, 2017). We trained on input patches of size 128×128×20 px^3^ at 7.16×7.16×40 nm^3^ resolution. The prediction during training was bilinearly upsampled to full image resolution before calculating the loss. Training utilized synchronous gradient updates computed by four Nvidia Titan X Pascal GPUs each with a different input patch. We used the AMSGrad variant (Reddi, Kale and Kumar, 2019) of the Adam optimizer (Kingma and Ba, 2014), with PyTorch’s default settings except step size parameter α = 0.001. We used the binary cross-entropy loss with “inverse margin” of 0.1 (Huang and Jain, 2013); patch-wise class rebalancing (Lee *et al.*, 2017) to compensate for the lower frequency of boundary voxels; training data augmentation including flip/rotate by 90°, brightness and contrast perturbations, warping distortions, misalignment/missing section simulation, and out-of-focus simulation (Lee *et al.*, 2017); and lastly several new types of data augmentation including the simulation of lost section and co-occurrence of misalignment/missing/lost section.

Distributed computation of affinity maps used chunkflow (Wu *et al.*, 2019). The computation was done with images at 7.16×7.16×40 nm^3^ resolution. The whole volume was divided into 1280×1280×140 chunks overlapping by 128×128×10, and each chunk was processed as a task. The tasks were injected into a queue (Amazon Web Service Simple Queue Service). For 2.5 days, 1000 workers (Google Cloud n1 - highmem-4 with 4 vCPUs and 26 GB RAM, deployed in Docker image using Kubernetes) fetched and executed tasks from the queue as follows. The worker read the corresponding chunk from Google Cloud Storage using CloudVolume (Silversmith and Tartavull, no date), and applied previously computed masks to black out regions with image defects. The chunk was divided into 256×256×20 patches with 50% overlap. Each patch was processed to yield an affinity map using PZNet, a CPU inference framework (Popovych *et al.*, 2020). The overlapping output patches were multiplied by a bump function, which weights the voxels according to the distance from patch center, for smooth blending and then summed. The result was cropped to 1024×1024×120 vx and then previously computed segmentation masks were applied (see Image defect handling above).

#### Watershed and size-dependent single linkage clustering

The affinity map was divided into 514×514×130 chunks that overlapped by 2 voxels in each direction. For each chunk we ran a watershed and clustering algorithm (Zlateski and Seung, 2015) with special handling of chunk boundaries. If the descending flow of watershed terminated prematurely at a chunk boundary, the voxels around the boundary were saved to disk so that domain construction could be completed later on. Decisions about merging boundary domains were delayed, and information was written to disk so decisions could be made later. After the chunks were individually processed, they were stitched together in a hierarchical fashion. Each level of the hierarchy processed the previously delayed domain construction and clustering decisions in chunk interiors. Upon reaching the top of the hierarchy, the chunk encompassed the entire volume, and all previously delayed decisions were completed.

#### Mean affinity agglomeration

The watershed supervoxels and affinity map were divided into 513×513×129 chunks that overlapped by 1 in each direction. Each chunk was processed using mean affinity agglomeration (Lee *et al.*, 2017; Funke *et al.*, 2019). Agglomeration decisions at chunk boundaries were delayed, and information about the decisions was saved to disk. After the chunks were individually processed, they were combined in a hierarchical fashion similar to the watershed process.

#### Synaptic cleft detection

Synaptic clefts were annotated by human annotators within a 310.7 μm^3^ volume, which was split into 203.2 μm^3^ training, 53.7 μm^3^ validation, and 53.7 μm^3^ test sets. We trained a version of the Residual Symmetric U-Net (Lee *et al.*, 2017) with 3 downsampling levels instead of 4, 90 feature maps at the 3rd downsampling instead of 64, and “resize” upsampling rather than strided transposed convolution. Images and labels were downsampled to 7.16×7.16×40-nm^3^ image resolution. To augment the training data, input patches were transformed by (1) introducing misalignments of up to 17 pixels, (2) blacking out up to 5 sections, (3) blurring up to 5 sections, (4) non-linear warping, (5) varying brightness and contrast, and (6) flipping and rotating by multiples of 90 degrees. Training used PyTorch (Adam *et al.*, 2017), and the Adam optimizer (Kingma and Ba, 2014). The learning rate started from 10^−3^, and was manually annealed three times (505k training updates), before adding 67.2 μm^3^ of extra training data for another 670k updates. The extra training data focused on false positive examples from the network’s predictions at 505k training updates, mostly around blood vessels. The trained network achieved 93.0% precision and 90.9% recall in detecting clefts of the test set, using parameters selected to maximize F1.5 score (biasing towards recall). This network was applied to the entire dataset using the same distributed inference setup as affinity map inference. Connected components of the thresholded network output that were at least 50 voxels at 7.16×7.16×40-nm^3^ resolution were retained as predicted synaptic clefts.

#### Synaptic partner assignment

Presynaptic and postsynaptic partners were annotated for 387 clefts, which were split into 196, 100, and 91 examples for training, validation, and test sets. A network was trained to perform synaptic partner assignment via a voxel association task (Turner *et al.*, 2020). Architecture and augmentations were the same as for the synaptic cleft detector. Test set accuracy was 98.9% after 710k training iterations. The volume was separated into non-overlapping chunks of size 7.33 × 7.33 × 42.7 μm^3^ (1024 × 1024 × 1068 voxels), and the net was applied to each cleft in each chunk. This yielded a single prediction for interior clefts. For a cleft that crossed at least one boundary, we chose the prediction from the chunk which contained the most voxels of that cleft. Cleft predictions were merged if they connected the same synaptic partners and their centers-of-mass were within 1 μm. This resulted in 3,556,643 final cleft predictions.

We found a small number of synapses targeting the PyCs whose assignment was incorrect. These synapses were outside of the manually proofread PyC-PyC subgraph, but we implemented a heuristic to remove these errors before further analysis. We mapped each PyC synapse to its closest mesh vertex by euclidean distance, and also mapped each PyC skeleton node to its closest mesh node in a similar manner. We then associated synapses to skeleton nodes by distance across the mesh. Associating synapses and skeleton nodes by means of the mesh, rather than directly, helped to handle cases where neurites passed nearby one another. Having associated synapses to skeleton nodes, we found the 9 “nearest neighbors” to each synapse by its associated skeleton nodes. Assignment errors were selected for manual review if a majority of these neighbors were of a different type than the original synapse (presynaptic vs. postsynaptic). This procedure found 1392 incorrect predictions out of a set of 894528 (~0.16%).

#### Mitochondria detection and assignment

Mitochondria were annotated by human annotators within 1069.8 μm^3^ of image data, which was split into 462.3 μm^3^ training, 57.00 μm^3^ validation, and 550.4 μm^3^ test sets. The mitochondria labels were eroded by 2 voxels in-plane in order to prevent merges from organelle contacts. We trained a version of the Residual Symmetric U-Net (Lee *et al.*, 2017) with 2 downsampling levels instead of 4, “resize” upsampling rather than strided transposed convolution, and [32, 40, 80] feature maps at each downsampling level. Images and labels were downsampled to 7.16×7.16×40-nm^3^ image resolution. To augment the training data, input patches were transformed by (1) introducing misalignments of up to 30 pixels, (2) blacking out up to 3 sections, (3) non-linear warping, (4) varying brightness and contrast, and (5) flipping and rotating by multiples of 90 degrees. Training used PyTorch (Adam *et al.*, 2017), and the Adam optimizer (Kingma and Ba, 2014). The learning rate started from 10−4 and was manually annealed twice (220k training updates). The trained network achieved 93.6% precision and 92.7% recall in detecting mitochondria of the test set. These metrics were defined by overlap, and errors for this network were dominated by mergers and splits of true segments rather than fully erroneous objects or missing object predictions. This network was applied to the entire dataset using the same distributed inference setup as affinity map and synaptic cleft inference. Connected components of the thresholded network output that were at least 300 voxels at 7.16×7.16×40-nm^3^ resolution were retained as predicted mitochondria. Each predicted mitochondrion was associated with the neuronal segmentation object with which it overlapped most within the final proofread segmentation. We further meshed each mitochondrion segment for visualization and further analysis (see below) using zmesh (Zlateski and Silversmith, no date).

#### Cell type assignment

Cells with somas inside the EM volume were classified as either excitatory pyramidal neurons, inhibitory interneurons, or non-neuronal cells based on morphological and synaptic criteria. PyCs (417 in the volume) were identified by the presence of a spiny apical dendrite radiating toward the pia, spiny basal dendrites extending laterally, and a basal axon that extended radially inward towards the white matter. PyCs axons formed asymmetric synapses. Cells that did not fit this description but that were associated with synapses and had dendrites or axons were classified as inhibitory interneurons (34).

Interneurons with identifiable axons inside the volume (20) were also assigned putative subtypes based on a combination of axonal and dendritic morphology as well as synaptic connectivity properties (Kawaguchi, Karube and Kubota, 2006; Kubota, 2014; Kubota *et al.*, 2016). Interneurons without axons were not assigned types. Basket cells (4) were identified by virtue of their highly branched, local-space-filling axonal arbors. They also had a larger number of primary dendrites, and at least 12% of their postsynaptic targets were pyramidal cell somata. Chandelier cells (2) were identified based on the cartridge-like groups of synapses formed by their axons and they targeted almost exclusively axon initial segments of pyramidal cells. One Martinotti cell was identified by its apical axon that targeted mostly dendritic shafts and spines of excitatory cells; consistent with (Kawaguchi, Karube and Kubota, 2006), it had four primary dendrites that were spiny. Bipolar cells represented a very diverse group of neurons that consistently had two to three primary dendrites. The dendrites were usually spiny and showed a vertical bias. Though it was rare for these cells to have an axon in the volume, in the few cases where this occurred there was also diversity, with some bipolar cells forming the majority of their synapses with inhibitory dendrites and another targeting mostly excitatory cells. Finally, one neurogliaform cell was identified. This cell had an axon with boutons that rarely formed synapses; consistent with (Kawaguchi, Karube and Kubota, 2006) it also had a large number of primary dendrites and, as described in the results section, possessed a very different pattern of synaptic inputs when compared with other cell types.

Non-neuronal cells were generally not associated with synapses and could be broadly classified by morphology as either vasculature-related or glial cells. Vasculature-related cells, i.e. endothelial cells and pericytes, wrapped tightly around blood vessels in the volume. Endothelial cells visibly formed the walls of blood vessels and the continuation of their somata into blood vessel walls was readily visible. The one pericyte we identified was wrapped around an endothelial cell, consistent with typical descriptions as in (Brown *et al.*, 2019). Endothelial cells could also be distinguished from pericytes by their dark, electron-dense cytoplasm.

Glial cells were categorized based on their overall morphology and ultrastructural features. Protoplasmic (cortical) astrocytes had angular shaped processes with electron lucent cytoplasm that contained numerous glycogen granules. The processes of the cells frequently filled interstices surrounding synapses and bore endfeet that covered the surface of blood vessels (Figure 1B inset). Microglia had electron-dense cytoplasm; dark, irregularly shaped nuclei; and thick branches with spiky endings (Figure 1C inset). Often, there were large lipofuscin granules (residues of lysosomal digestion) in the cell body, whereas lysosomes and scattered phagolysosomes were present in branches. Microglia had long strands of endoplasmic reticulum (ER) in their granular cytoplasm and few microtubules. These are the smallest of neuroglial cells (they differ from other types because of their hemopoietic origin Ginhoux and Prinz, 2015). Most of the microglia in this dataset were also perineuronal satellites and were located adjacent to their host neurons with an astrocytic process between them (Peters, L. and Webster, 1978). Perivascular microglia were in close association with blood vessels, and some of the processes contacted surfaces of the endothelial cells forming the vessel. We grouped both of the microglial types into a single microglia group. OPCs resembled microglia because of their highly ramified branches and perinuclear satellite position with neurons, however they were much larger, with branches extending up to 50 μm from the cell soma (Peters, 2004; Figure 1D inset) Their nuclei were pleomorphic, with dense heterochromatin patterning and undulating nuclear envelopes. The branches were thick near the soma, diving into smaller processes with terminations that bore growth-cone-like structures and filopodia. The cytoskeleton of the cells was composed of numerous microtubules and there were many organelles throughout. As the OPCs transitioned to oligodendrocytes, morphological changes became evident (Trapp *et al.*, 1997). The nucleus was elongated with a smooth nuclear envelope and the branches appeared thinner and less ramified. The branch terminations wrapped axons, and there was evidence of myelin formation with uncompacted myelin present as well. Mature oligodendrocytes had a compact elongated soma, with scant cytoplasm in the soma. The branches were thin, smooth and straight and contained microtubules. The branches terminated in spiral membrane formations of compact myelin, enwrapping up to 60 axons (Simons and Nave, 2015; Figure 1E inset).

Each cell with a soma in the volume received at least 2x coverage for identification and the consensus is reported here.

#### PyC proofreading

The mean affinity graph of watershed supervoxels was stored in our PyChunkedGraph backend, which uses an octree to provide spatial embedding for fast updates of the connected component sets from local edits. We modified the Neuroglancer frontend (Maitin-Shepard, 2019) to interface with this backend so users directly edit the agglomerations by adding and removing edges in the supervoxel graph (merge and split agglomerations). Connected components of this graph are meshed in chunks of supervoxels, and chunks affected by edits are updated in real-time so users can always see a 3D representation of the current segmentation. Using a keypoint for each object (e.g. soma centroid), objects are assigned the unique ID of the connected component for the supervoxel which contains that location. This provides a means to update the object’s ID as edits are made.

Cell bodies in the EM volume were manually identified. PyCs were identified by morphological features, including density of dendritic spines, presence of apical and basal dendrites, direction of main axon trunk, and cell body shape. A team of annotators used the meshes to detect errors in dendritic trunks and axonal arbors, then to correct those errors with 50,000 manual edits in 1,044 person-hours. After these edits, PyCs were skeletonized, and both the branch and end points of these skeletons were identified automatically (with false negative rates of 1.7% and 1.4%, as estimated by annotators). Human annotators reviewed each point to ensure no merge errors and extend split errors where possible (210 person-hours). Putative broken spines targeted by PyCs were identified by selecting objects that received one or two synapses. Annotators reviewed, and attached these with 174 edits in 24 person-hours. Some difficult mergers came from small axonal boutons merged to dendrites. We identified these cases by inspecting any predicted presynaptic site that resided within 7.5 μm of a postsynaptic site of the same cell, and corrected them with 50 person-hours.

#### PyC-PyC synapse proofreading

Synapses between PyCs were extracted from the automatically detected and assigned synapses. We reviewed these synapses manually with 2x redundancy (1972 correct synapses out of 2433 putative synapses). 2 predicted synapses out of these were “merged” with other synaptic clefts. These cases were excluded from further analysis. 1 synapse was “split” into two predictions, and these predictions were merged for analysis.

#### Nucleus detection

Nucleus segmentation was produced by a convolutional neural net predicting the likelihood of being part of a nucleus for each voxel (at the downsampled 57.28×57.28×40 nm ^3^ voxel resolution), followed by a distributed connected components process merging together voxels with a likelihood above the threshold of 0.5.

The convolutional neural net was trained on image data and labels from two other similar EM datasets to be presented elsewhere. The network architecture was a modified version of the Residual Symmetric U-Net introduced in (Lee *et al.*, 2017). Specifically, in each residual module, the first convolution used a kernel size of 1 and was effectively a simple linear layer; a total of 3 downsampling levels were used instead of 4, with [16, 32, 64, 128] feature maps at each downsampling level. An input patch size of (160, 160, 32) was used in training.

Each nucleus was manually checked to identify all cells with somas in the volume.

#### Skeletonization

We generated two separate sets of skeletons for the neuronal segmentation. The first set performed bulk processing of every object in the volume. These skeletons are useful for measuring the lengths of arbitrary objects, their orientations, and topologies. The second set was only produced for specific objects of interest (neurons with soma), and were intended to better handle two failure modes of bulk skeletonization: discontinuous objects generated by proofreading across image defects and “self-contact” sites. This second set of skeletons was used for the analyses within each vignette, though we release both sets for general use.

The bulk set of skeletons were produced using a modified form of TEASAR (Sato *et al.*, 2000; Bitter, Kaufman and Sato, 2001) that operates on 3D labeled images. The dataset was divided into a grid of 513×513×513 cutouts at 28.64×28.64×40 nm^3^ resolution with an overlap margin of one voxel in each dimension. Conceptually, each 26-connected component within each cutout was extracted as a binary image and we serially extracted a set of centerlines that form a tree. We selected the foreground root as the furthest foreground voxel from an arbitrarily selected foreground voxel except at somata where the root is set as the foreground voxel with maximum distance from the boundary.

We then perform TEASAR with the following modifications: (1) In the PDRF, we set M as max(DBF)^1.01^, the coefficient to 100,000, and lower the exponent to 4 (2) We normalize the DAF to 1 (3) the invalidated region uses cubes instead of spheres (4) we weight the PDRF to zero along already traced paths, which results in better branching behavior and faster tracing when tracing from target to root. (5) We also assigned priority tracing targets from the centers of the cross sections of the shape that touch the border. We then trivially fused all the skeleton fragments for each neuron together at the overlapping voxel, removed detached components smaller than 1 μm, removed loops introduced by the forest fusion, and joined nearby detached components if they were nearer to each other than the EDT radius of the closest two vertices. We make our implementation available through our packages Kimimaro (Silversmith and Bae, no date) and Igneous (Silversmith and Tarvatull, no date). The procedure was set to stop tracing after 50 paths in a given cutout, which tends to exclude glia.

To produce the second set of skeletons, we developed a skeletonization algorithm similar to TEASAR (Sato *et al.*, 2000) that operates on meshes. For each connected component of the mesh graph, we identify a root and find the shortest path to the farthest node. This procedure is repeated after invalidating all mesh nodes within the proximity of the visited nodes until no nodes are left to visit. We make our implementation available through our package MeshParty (https://github.com/sdorkenw/MeshParty). The skeletonization parameters were hand-tuned to largely ignore dendritic spines.

The results of these skeletons were further processed to improve our estimations of path length distance across cells. Each skeleton was smoothed by iteratively shifting vertices towards a local average computed using a vertex’s neighborhood. Given a vertex, the smoothing neighborhood around consists of the set of vertices less than or equal to 2-edges from (excluding itself). After computing the local average from the neighborhood, each vertex is moved to a weighted sum of its original location and its local average 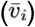 with a blending parameter 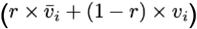.

We used *r*=0.1, and performed 20 iterations of this smoothing. The soma root node, branch point nodes, and leaf nodes were kept fixed throughout this smoothing procedure.

#### Neuronal compartment labeling

Annotators tagged all branches leaving the soma approximately 5 μm away from the cell body and labeled them as one of: “axon”, “basal dendrite”, “apical dendrite”, “ambiguous dendrite”, “ambiguous”. We next mapped these annotations to the closest skeleton nodes in the automatically generated skeletons and moved the label to the closest branch node if one was within 5 μm. Labels were then propagated to all skeleton nodes away from the soma. Imperfect skeletons introduced errors in the label propagation. We qualitatively inspected the resulting compartment labels and designated a conservative set of 125 cells with ideal compartment labels. Out of this set, we manually identified 65 that had their somas fully captured by the EM volume.

## QUANTIFICATION AND STATISTICAL ANALYSIS

### Neurite length calculation

The skeleton for each PyC and inhibitory neuron was split according to its compartment labels to generate individual sub-skeletons for dendrites and for the axon. For each sub-skeleton, the node closest to the soma centroid was chosen as the source and all other leaf nodes (nodes with degree 1) were identified as targets. Each leaf node represented either the biological end of a neurite (e.g. the end of a short dendrite in the volume) or the point at which a longer neurite left the volume. The shortest path length was computed from the source to each target node with Dijsktra’s algorithm. We summarized this distribution between its 5th and 95th percentiles to remove a few large outliers.

### Synapse detection performance estimation

To estimate the precision of synapse detection for each class, we defined a “likely subset” of synapses belonging to that class based on a heuristic, and scored synapses that were of the correct type. For example, for axo-axonic synapses, we extracted the set of synapses received by PyC’s within 10 μm of the axon and soma compartment label boundary. In some cases (mostly axo-somatic), a synaptic cleft would be split into two predictions by an image defect or weak prediction. In those cases, we counted that prediction as 50% true positive and 50% false positive.

To estimate the recall for each class, we randomly sampled a set of cells, manually identified a set of synapses for those cells by traversing their neurites or target domains, and compared the manually identified set to the predicted clefts. Each precision and recall value was estimated with roughly 50 synapses each.

### Synapse/connection degree and synapse/connection density

Using the 334-node PyC graph, we counted incoming and outgoing synapses within the graph are named as in-synapse degree and out-synapse degree respectively. The number of unique presynaptic partners and postsynaptic partners are named as in-connection degree and out-connection degree respectively. To compensate for the truncated effect due to limited size of the volume, we normalized these quantities by axon length and dendrite length. These normalized quantities are synapse/connection density. We also looked at all outgoing synapses on axons and incoming synapses on dendrites, not restricted to the PyC graph, and named these quantities consistent with terms used for PyC graph but added “total” as prefix. For incoming synapses on dendrites, we filtered out synapses which are within 15 μm from the human-labeled soma centroid to remove perisomatic synapses.

### Inhibitory cell input analysis

In order to define compartments for inhibitory neurons, we manually placed labels near the axon initial segment for each cell if it had an axon in the volume. The label was extended to the branch point of the skeleton closest to the cell body. As with pyramidal cells, the soma was defined to include the region within 15 μm of its soma center label. In order to make neuronal meshes and skeletons continuous, links were added to the meshes where proofreading spanned a gap in the segmentation using the MeshParty function “add_link_edges.” Synapses were associated with the mesh vertex closest to the location of the synaptic cleft in the synapse detection and synapses beyond 200 nm from a mesh vertex were omitted. Mesh vertices and their synapses were bidirectionally mapped to the closest skeleton vertex as measured along the surface of the mesh.

Statistical visualization was performed in python using Seaborn (Michael Waskom and the seaborn development team, no date). Confidence intervals for line plots were estimated by bootstrap resampling the density values within each distance bin and type (1000 bootstraps, as implemented by default in Seaborn). For violin plots, synapses were first log_10_-transformed and then plotted with a kernel bandwidth of 0.1. Statistical tests were performed with Scipy 1.5.2 (Virtanen *et al.*, 2020) and multiple comparison correction was done with statsmodels 0.11.1 (Seabold and Perktold, 2010).

### Mitochondrion analysis

We associated each mitochondrion mesh to the set of skeleton nodes that were closest to any mesh vertex of that object within the 125 PyCs with ideal compartment labels. We then assigned a compartment label to each predicted mitochondrion by taking a majority vote of the skeleton node labels associated with that object. Mitochondria with an “ambiguous dendrite” or “ambiguous” label were removed from further analysis.

We computed the path length distance along the computational skeleton to the nearest somatic node of a given cell. The linear mitochondrial coverage factor within a section of neurite was computed by first measuring the path length within this section covered by each present mitochondrion, and then normalizing the sum of these covered lengths by the amount of neurite path length within the section (Figure S4A). When assigning a single location to a mitochondrion (Figure S4C), we took the mean path length value across all skeleton nodes associated with that mitochondrion.

We further restricted the density analysis to the 65 cells whose somata were fully captured by the EM volume. We then computed the linear coverage factor over all skeleton nodes with a basal dendrite label. In order to compute the volume coverage factor, we first measured soma volume by cutting each PyC’s computational mesh at a 15 μm radius from the human-labeled soma center, and measured the volume of the result. We then subtracted the volume of the predicted nucleus segment for this cell to measure cytosolic volume. We finally took the sum of all mitochondrion volumes which were only associated with skeleton nodes within 15 um radius of the soma center, and divided that sum by the cytosolic volume measurement. We computed soma synapse density by normalizing the number of received synapses within a 15 μm radius from the human-labeled soma center by the surface area of the soma mesh cutout. The two-tailed p-value was calculated for each Pearson correlation as implemented in scipy: the null distribution of the r-value is a Beta distribution over [-1, 1] with shape parameters *α*=*β*=n/2-1.

#### Random network models

To identify nonrandom connectivity motifs, we compared the simple subgraph of PyCs (removing self-loops and multiple edges) with different random graphs as null models. When studying two-neuron motifs, we first employed a directed Erdős-Rényi (ER) model (Gilbert, 1959), *G*(*n*, *p*), as a baseline, where a directed edge between any two distinct vertices is uniformly drawn at random with probability p. The expected number of edges of directed ER model*m* = *n*(*n* − 1)*p*. To match the data, for example the subgraph with a 100 μm axon length threshold, we set *n* = 113, *m* = 666, and *p* = *m*/[*n*(*n* − 1)] ≈ 0.0526. We directly calculated the expected number of different two-neuron motifs in this ER model.

We also employed a directed configuration model, 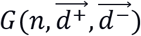, which generates rewired graphs preserving the original degree distribution uniformly at random, where 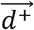 and 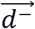 are predefined in- and out-degree sequences of the network. We sampled 1,000 random networks uniformly from a set of graphs with the same degree sequences as our PyCs subgraph by applying the switch-and-hold algorithm (Figure 5A; Artzy-Randrup and Stone, 2005), where we randomly select two edges in each iteration, swap the target endpoints of them if not introducing self-loops or multiple edges (switch), otherwise keep them unchanged (hold). Note that in the original paper, the switch step is implemented by selecting four vertices at random, while our implementation selects two edges at random, since it is more efficient on a sparse network. In general, the Markov chain for our sampling procedure is reducible, so samples are drawn from only one ergodic component of the set of all simple graphs with the same degree sequence (Carstens and Horadam, 2016). However, all ergodic components share the same motif frequencies so the lack of ergodicity has no consequence for our analysis (Berger and Müller-Hannemann, 2010). A more complex Markov chain sampling procedure with true ergodicity is known (Roberts and Coolen, 2012), but as mentioned above would yield the same motif frequencies. To generate one sample, we run the algorithm for 10,000 iterations from the previously sampled network, and the average holding rate is 19.2% for the largest PyC subgraph.

When studying three-neuron motifs, besides the vanilla ER model and configuration model as described above, we also modified the ER model (Figure 5B) and the configuration model to preserve two neuron motif statistics. In the generalized ER model, the probability of a unidirectional edge is 4.80 × 10^−2^, and the probability of a bidirectional edge is 4.58 × 10^−3^. We directly calculated the expected number of different three-neuron motifs in the generalized ER model.

We also modified the switch-and-hold algorithm such that the number of bidirectional edges is also preserved (Data S5A). Sampling from this generalized configuration model involves two phases: mixing and hitting. For each sample, we run the switch-and-hold for 10,000 iterations to ensure the convergence to stationary distribution (mixing), then continue for a few iterations until it reaches a graph with the same number of bidirectional edges as the observed network (hitting). This generalized configuration model efficiently samples graphs with unchanged degree sequences and the statistics of two-neuron motifs without biases. It has the same holding rate and running time as ordinary switch-and-hold, and only increases the required number of iterations by a small constant Θ(log(1/(1-p)), where p is the percentage of graphs with the same freq. of 2-neuron motifs in the entire space. In the experiment, we sampled 1,000 random networks with an average holding rate of 19.2% for the largest PyC subgraph. The average number of iterations in the hitting phase is 3,736.78, which only increases the total number of iterations by 37.4%.

#### Direction and orientation selectivity

For each calcium-imaged cell, we defined a response matrix *R*_*td*_ as the peak amplitude of the response triggered by the stimulus in trial *t* to stimulus direction θ_*d*_. Peak amplitude was measured as the maximum value within the trial. When the maximum value is observed at the end of the trial time window, peak amplitude was measured as the maximum value within the next 15 frames (~1s, which is approximately the same as the duration of the stimulus). When the maximum value is observed at the start of the time window or the value at the start of the time window is greater than half of the peak response, we set the peak amplitude to be the same as the baseline assuming it’s the response decaying from the previous stimulus. The *T* × *D* matrix included response amplitudes in *T* = 30 trials and *D* = 16 directions. We approximated the response matrix using singular value decomposition (Baden *et al.*, 2016).

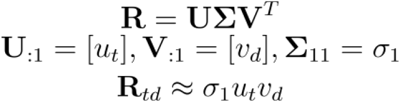

From the direction tuning function *ν*_*d*_, we computed a direction selectivity index (DSi) and orientation selectivity index (OSi) as below.

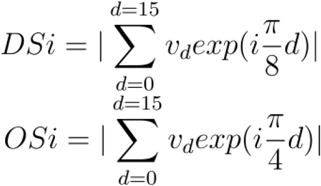

An orientation-tuned cell was defined as one with statistically significant OSi, evaluated using a permutation test (Ecker *et al.*, 2014; Baden *et al.*, 2016). A direction-tuned cell was defined as one with statistically significant DSi but not OSi. Statistical significance was determined by p<0.05 where the p-value indicates the fraction of 10,000 permutations with DSi/OSi greater than that of the observed data. The permutation test effectively excludes cells that have a large DSi or OSi merely by chance, which can happen due to trial-to-trial variability. Restriction by statistical significance is different from imposing a simple threshold on DSi or OSi (Baden *et al.*, 2016).

#### Intermittency

We defined a binary-valued function by thresholding the activity after temporal deconvolution where the threshold was set at three standard deviations greater than the baseline. For example, the value of this function for a directional stimulus with direction *d* and trial *t* is defined as

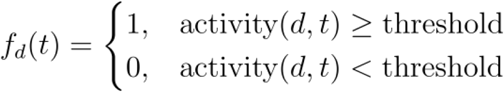

A cell’s intermittency is an inverse measure of how consistently a cell responds to preferred stimuli. Here, preferred stimuli was determined by finding the directions where *v*_*d*_ is greater than the average. Therefore, the intermittency value was defined as

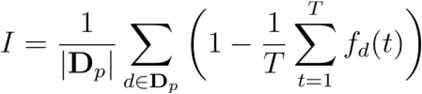

where D_*p*_ is a set of directions in preferred stimuli and is the number of trials for each stimulus. The Pearson correlation coefficient was calculated to measure the correlation between the intermittency and the in-connection density using the scipy package. To test for significance, the two-tailed p-value was calculated; a threshold of 0.05 has been used to reject the null hypothesis, where the null distribution of the r-value is a Beta distribution over [−1, 1] with shape parameters *α*=*β*=n/2-1.

#### Mean response

From the response matrix, a cell’s mean response was computed by taking the average of peak responses in all 30 trials of the preferred stimuli. To test for significance, we performed an analogous permutation test for intermittency.

We fit our linear model combining active responses and in-connection density using ordinary least squares as implemented in statsmodels (Seabold and Perktold, 2010).

#### Spatial location restricted permutation test

To test that significance of correlation is not due to the spatial organization, we performed a permutation test with spatial restriction. We divided the volume into small cubes by dividing the x- and z-dimensions into 2 regions and y-dimension (cortical depth) into 4 regions. Axis along the cortical depth was divided into finer regions because more distinct spatial organization existed along that axis (Figure S7) while other axes had less bias. For each cube, we have located all the cells that have soma centers within the cube and shuffled the in-connection density values among these cells. Then, we computed the correlation coefficient for the randomized distributions. Statistical significance was determined by p<0.05 where the p-value indicates the fraction of 10,000 permutations with correlation coefficient stronger than that of the observed data. We performed this permutation test for mean response and intermittency.

#### Statistical reporting

We report p-values greater than 1 × 10^−4^ as a rounded value, those under 1 × 10^−50^ as approximately 0, and those in between 1 × 10^−50^ and 1 × 10^−4^ as less than the value rounded up to two significant figures. We report p-values for permutation tests as the percentage of permutations whose test statistic value is more extreme than the observed value.

#### Rendering information

The following segment IDs were removed from the insets in Figure 1A to improve their visual clarity: 648518346349526102, 648518346349525545, 648518346349531742, 648518346349508442.

## DATA AND CODE AVAILABILITY

EM image data and cellular segmentation can be viewed at https://microns-explorer.org/. The mitochondria reconstruction will be made available in a timely fashion at the same location. Raw calcium imaging data are available together with the analysis code at https://github.com/seung-lab/MicronsBinder.

The code for generating the cell and organelle reconstructions is available at https://github.com/seung-lab/.

## Supplementary Information

**Figure S1.**
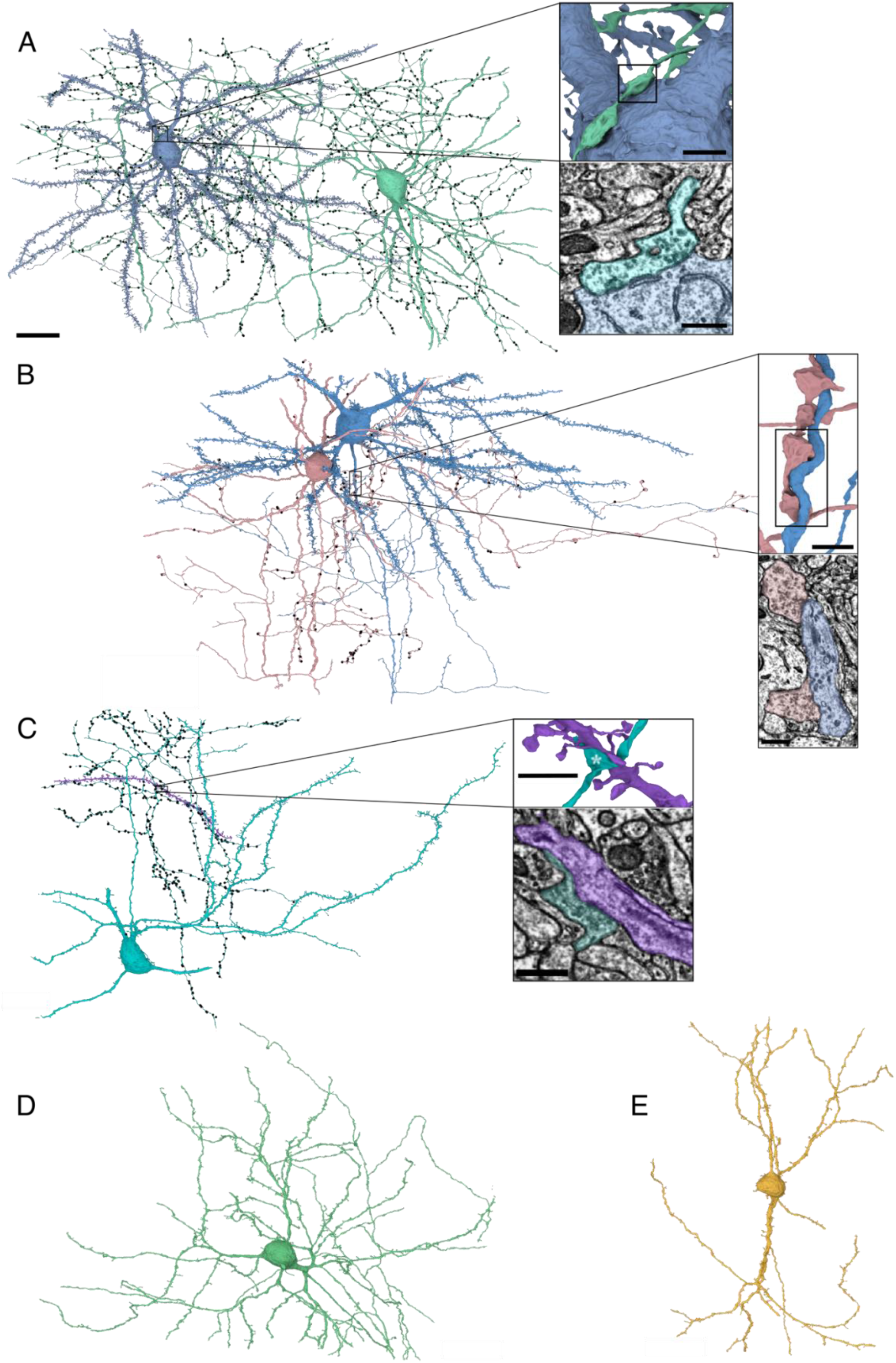
Example inhibitory neurons and the synapses they form, Related to Figure 1. (A) Axosomatic synapse formed by a basket neuron (green) onto a pyramidal cell (blue). Upper inset: view of the direct synaptic contact between axonal bouton and the pyramidal soma (box); lower: EM image of this region showing pleomorphic vesicles and symmetric pre- and postsynaptic density typical of inhibitory synapses. (B) A cartridge of axoaxonic synapses formed by a chandelier neuron (pink) onto a pyramidal neuron (blue). Upper inset: a series of chandelier presynaptic boutons target the axon initial segment of the pyramidal cell (box); lower: EM image of two of these synapses. (C) An axodendritic synapse formed by a Martinotti neuron (cyan) onto a pyramidal dendrite (purple). The dendrite has a natural end in the volume (right) and comes from a pyramidal cell located outside the volume bounds (left). Upper inset: the Martinotti axon forms a presynaptic bouton (asterisk) directly onto the shaft of the spiny pyramidal dendrite; lower: EM image showing pleomorphic vesicles and symmetric postsynaptic density at this synapse. (D) The neurogliaform cell identified in the dataset. (E) A bipolar cell (with basal axon). Scale bars: (A-E) 20 μm, upper insets 2 μm, lower insets 500 nm.

**Figure S2.**
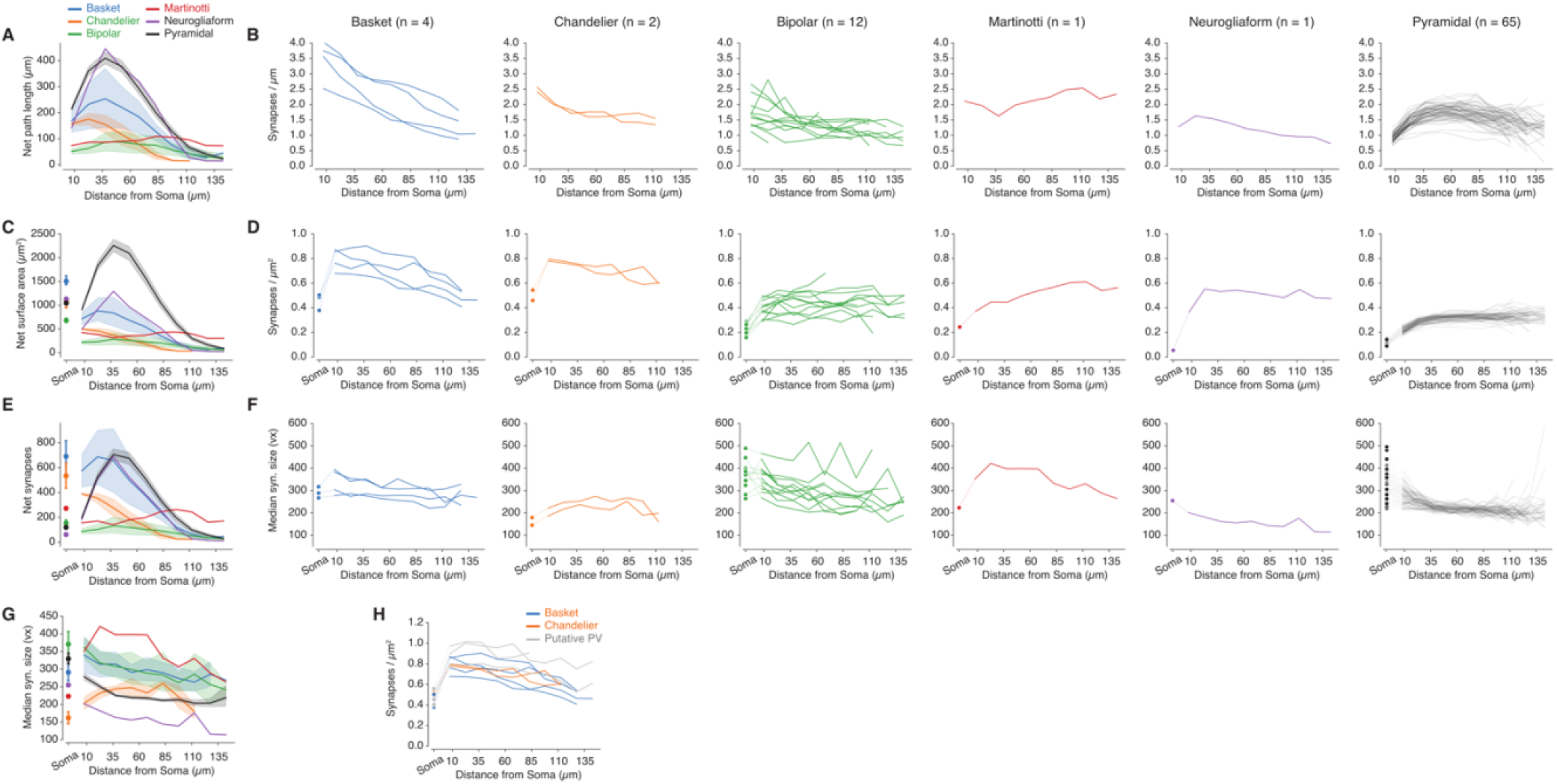
Input synapse density for excitatory and inhibitory cells, Related to Figure 3. (A) Path length captured by the volume for each coarse neuron type as a function of distance from the soma. (B) Linear synapse density for each cell grouped in Figure 3. (C) Neurite surface area for each coarse neuron type. (D) Surface synapse density for each cell grouped in Figure 3. (E) Synapse count for each coarse neuron type. (F) Median synapse size for each cell grouped in Figure 3. (G) Median synapse size grouped by coarse neuron type (H) Surface synapse density for basket and chandelier groups, as well as unclassified cells suspected to be within these types.

**Figure S3.**
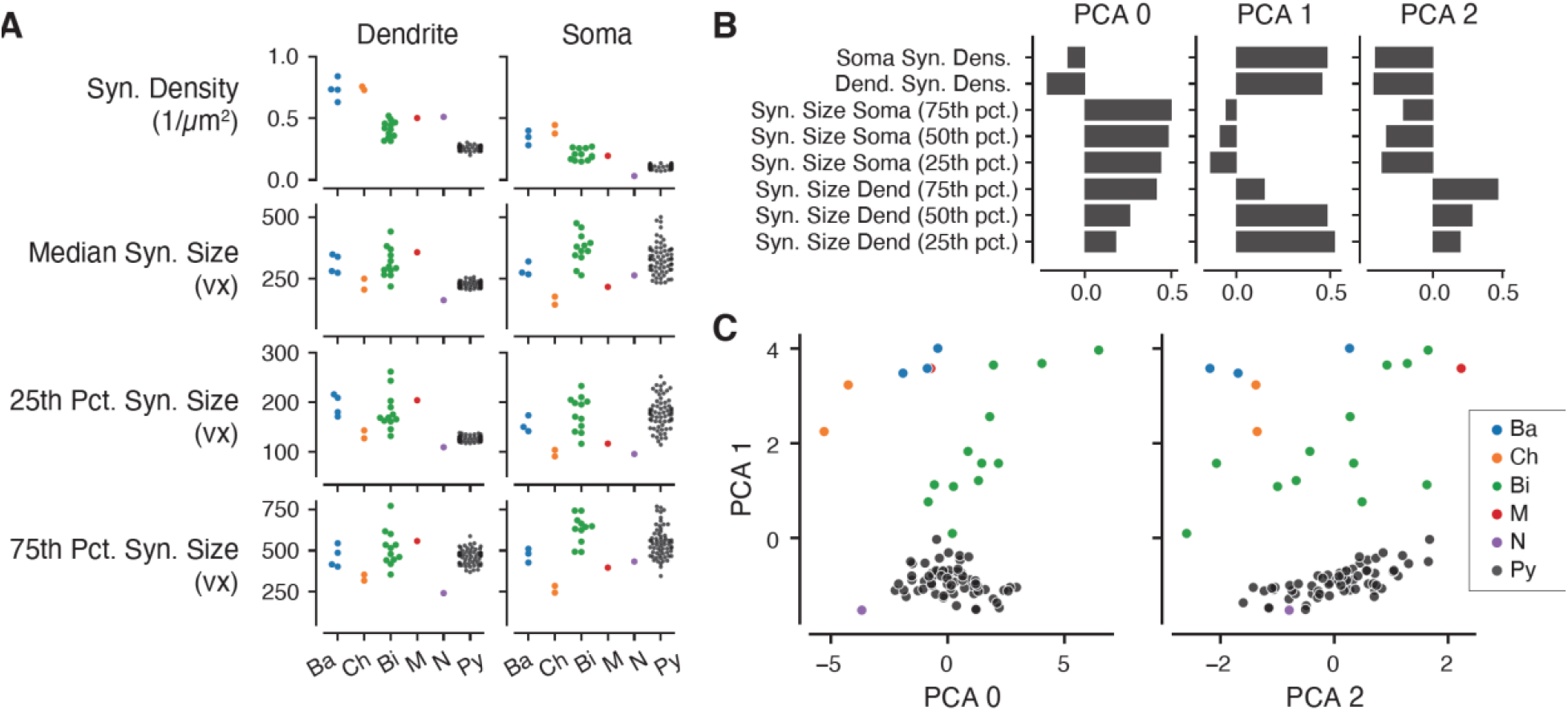
Inhibitory single cell data and grouping, Related to Figure 3. (A) Synapse features for each cell grouped in Figure 3. (B) First three principal components of the features in (A). These components explain 92% of the total variance. (C) PCA plots of cells grouped in Figure 3.

**Figure S4.**
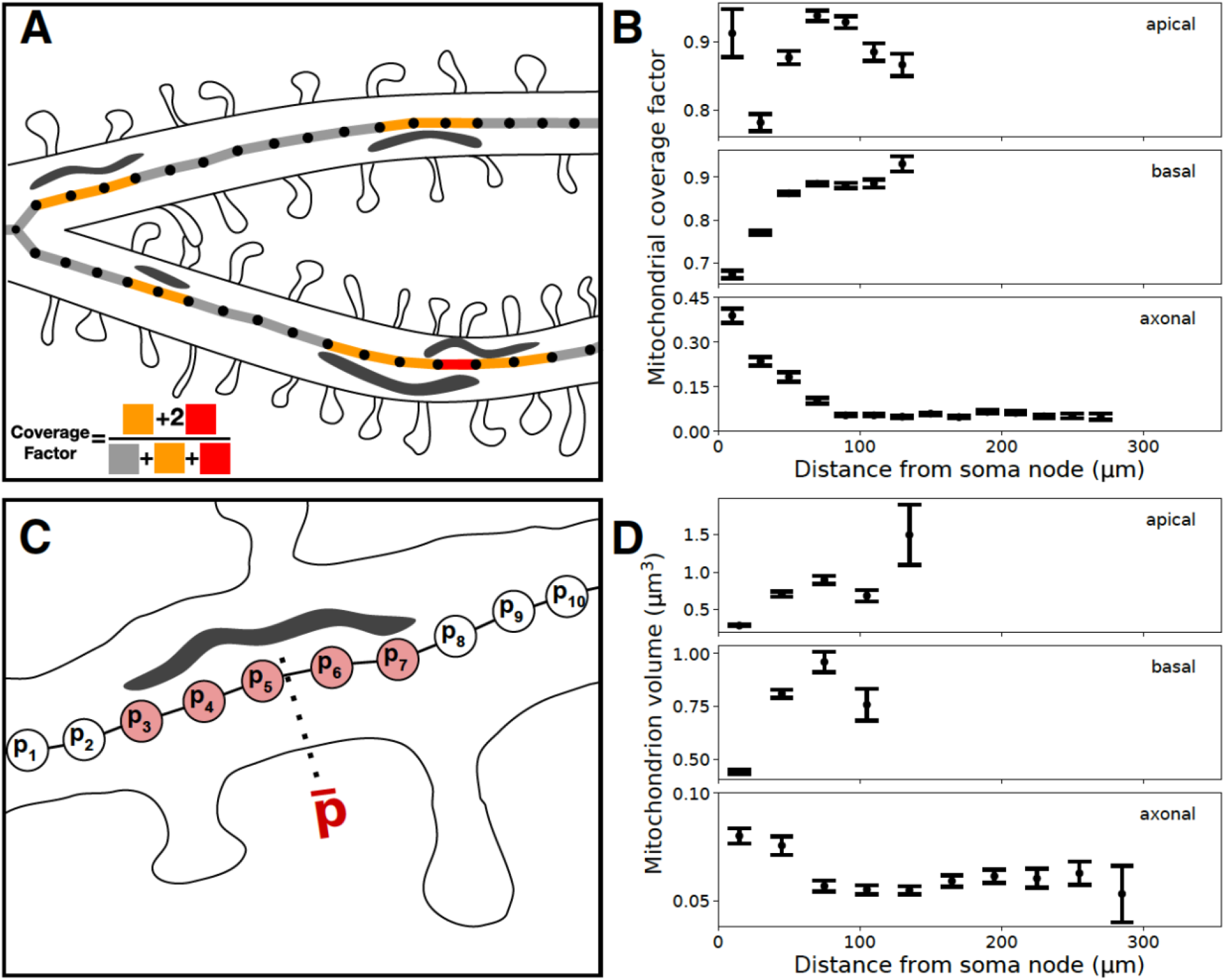
Distribution of mitochondria over pyramidal arbors, Related to Figure 4. (A) Cartoon depiction of mitochondrial index. Skeleton nodes are associated with a mitochondrion by proximity (STAR Methods). Mitochondrial coverage factor is defined as the ratio of mitochondria path length to neurite path length. (B) Mitochondrial coverage factor computed at different distances from a pyramidal neurons soma (20 μm intervals) (C) Cartoon depiction of associating each mitochondrion with a single path length from its soma. Skeleton nodes are associated to each mitochondria by proximity (STAR Methods). Each mitochondrion is labeled with the average path length distance to its soma. (D) Mitochondria volume at different mean path length distances from a cell’s soma (30 μm intervals). Mitochondria at the boundaries of the reconstruction can be truncated or otherwise corrupted.

**Figure S5.**
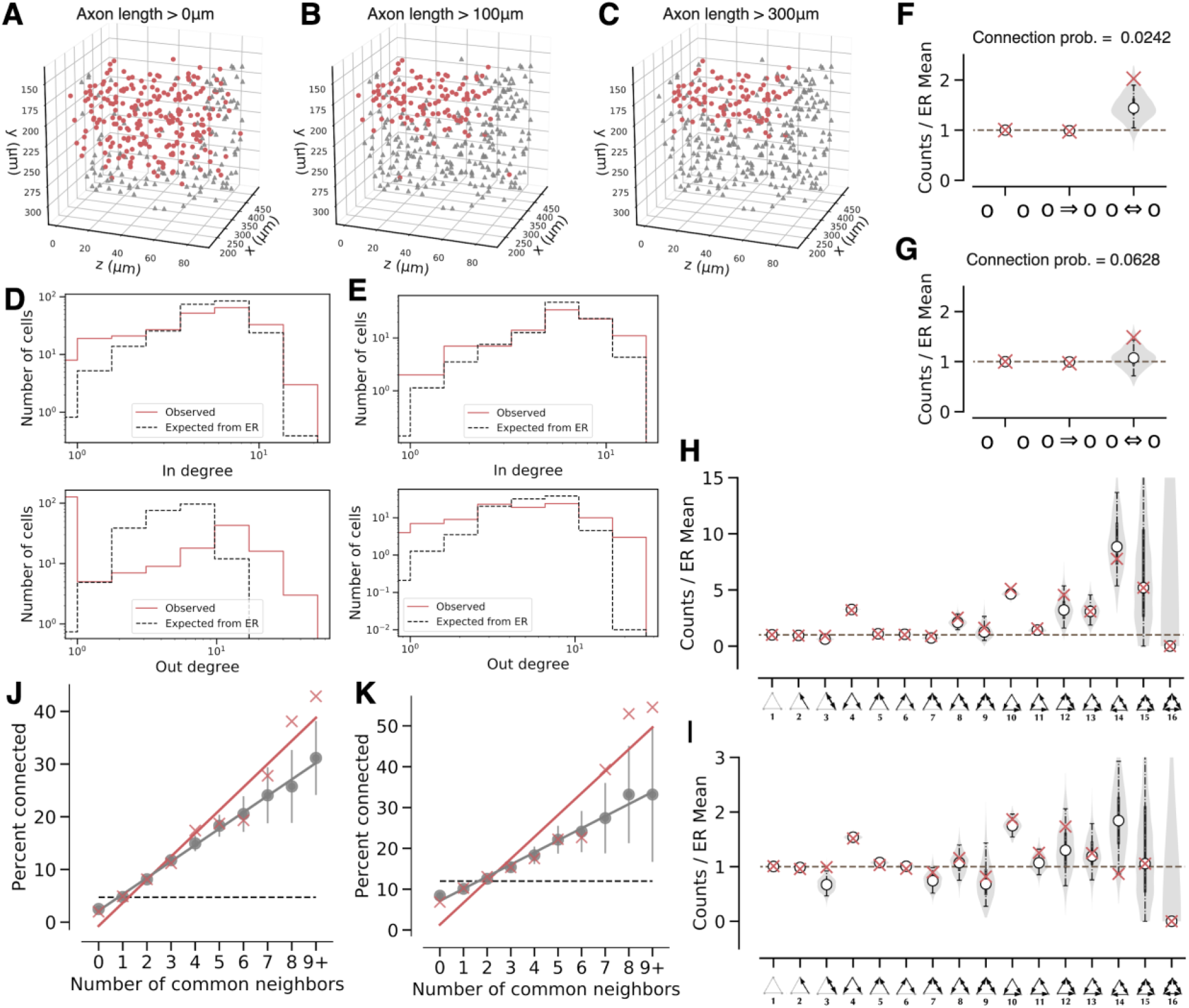
Predicting motif frequencies for PyC subgraphs with different axon length thresholds (0 μm and 300 μm), Related to Figure 5, which is based on a subgraph consisting of PyCs with axon lengths ≥ 100 μm. (A) Spatial location of nodes in PyC subgraphs with axon length threshold = 0 μm. Red: 229 PyCs included in the subgraph. Gray: PyCs excluded due to axon lengths shorter than threshold. (B) Similar to (A) for axon length threshold = 100 μm. The resulting subgraph contains 113 neurons. (C) Similar to (A) for axon length threshold = 300 μm. The resulting subgraph contains 100 neurons. (D) In and out degree distributions in an actual network with threshold = 0 μm and the expected distribution of in and out degrees in a regular ER model (edge probability = 0.0242). Red: observed data; Black, dashed: ER model. Histograms are calculated with 8 bins of the same widths on a log scale. The expected degree distributions are estimated from 100 samples from the ER model. (E) Similar to (D) for axon length threshold = 300 μm. ER edge probability = 0.0628. (F) 2-cell motif frequencies in an actual network with threshold = 0 μm and a configuration model relative to the ER model. The red crosses indicate observed counts. The shaded region shows the smoothed distribution of motif counts sampled from the configuration model. White points indicate medians, vertical lines indicate quartiles and the 95% confidence interval for 1,000 samples. (G) Similar to (F) for axon length threshold = 300 μm. (H) 3-cell motif frequencies in an actual network with threshold = 0 μm and the configuration model relative to a generalized ER model. The red crosses indicate observed counts. The violin plot shows the smoothed distribution of the motif counts sampled from the configuration model. White points indicate medians, vertical lines indicate quartiles and the 95% confidence interval for 1,000 samples. (I) Similar to (H) for axon length threshold = 300 μm. (J) The common neighbor rule for the graph with threshold = 0 μm. Linear fits are guides to the eye (red: observed data, r^2^=0.95, p=2.5e-6; gray: configuration model, r^2^≈1.00, p=1.2e–10; z-test, dashed line: generalized ER model). Error bars indicate standard deviation of 100 samples. (K) Similar to (J) for axon length threshold = 300 μm. Red: r^2^=0.88, p=6.3e-5; Gray: r^2^=0.99, p=7.8e–9; z-test.

**Figure S6.**
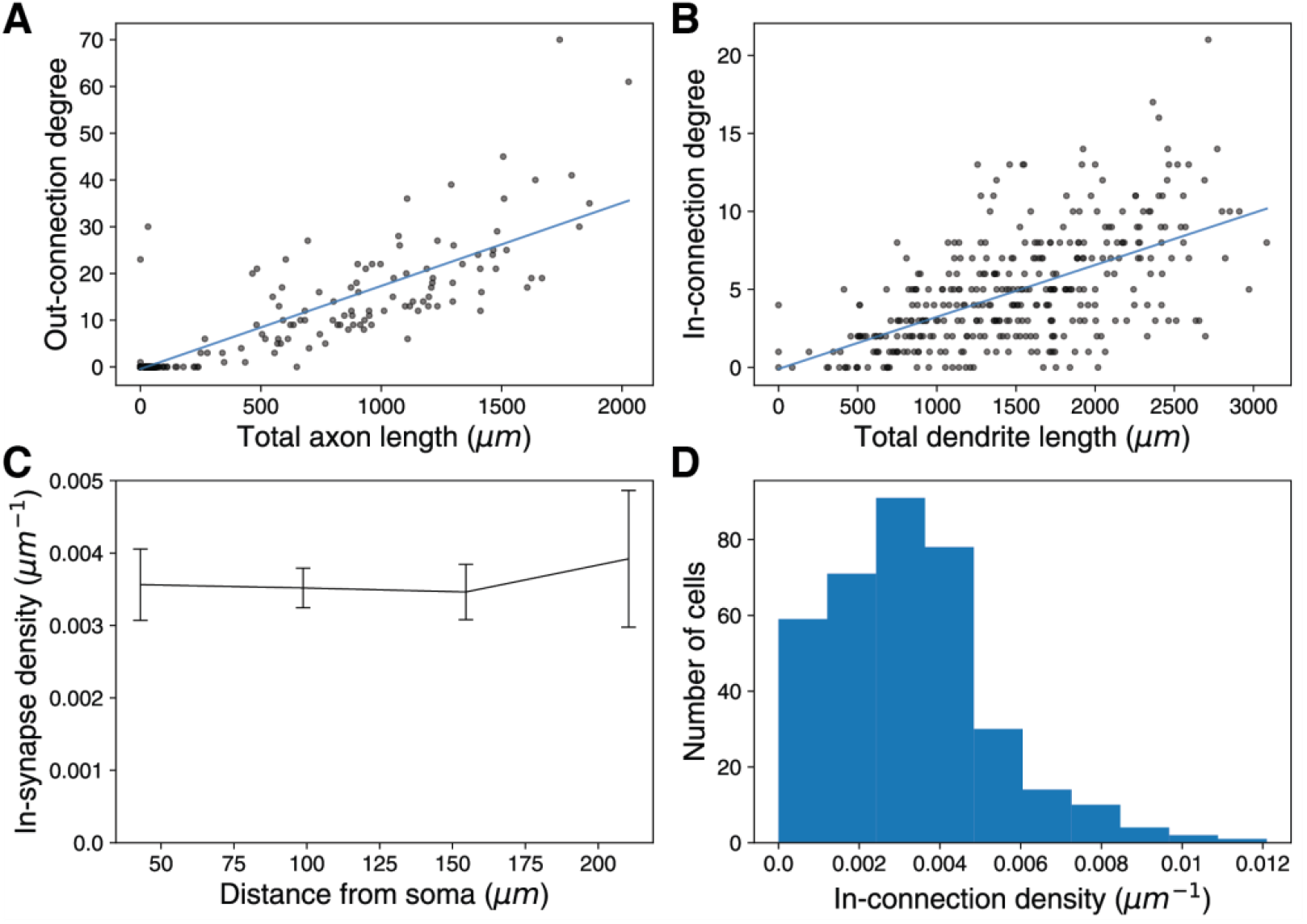
Normalization of in-connection degree to accommodate truncation effect due to the restricted volume, Related to Figure 6. (A) Total axon length is correlated with out-connection degree (*n*=363, Pearson’s r=0.87, p≈0). (B) Total dendrite length is correlated with in-connection degree (*n*=363, Pearson’s r=0.59, p<3.5e-35). (C) In-synapse density is consistent regardless of how far the synapses are located from the soma (error bar: mean±std.err.). (D) In-connection density varies between cells (*n*=363) (A and B) Line: linear fit.

**Figure S7.**
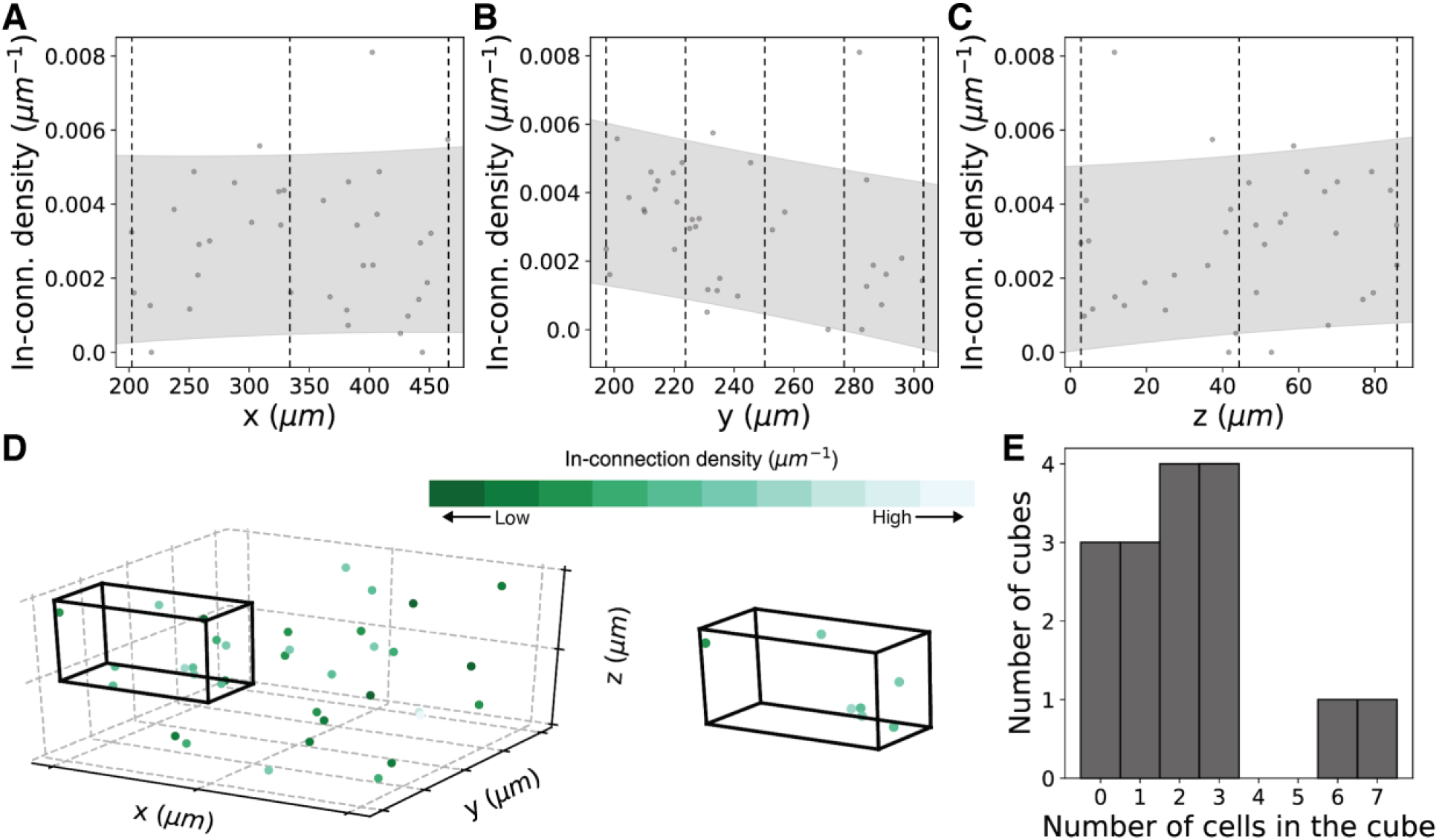
Spatial location restricted permutation test, Related to Figure 6. (A-C) In-connection density distribution along x-dimension (A), y-dimension (depth) (B), and z-dimension (C) of soma location (*n*=36). Dashed line: border boundary for shuffling, Shade: 80% prediction interval. (D) To generate randomized distributions, in-connection densities are shuffled among the cells in the same cube. 3D plot of soma locations (left) and example cube (right) are shown. Dashed lines are consistent with (A-C). (E) Histogram of the number of cells in each cube.

**Data S1. Inhibitory cell profiles (see bioRxiv Supplemental Material)**

**Data S2.**
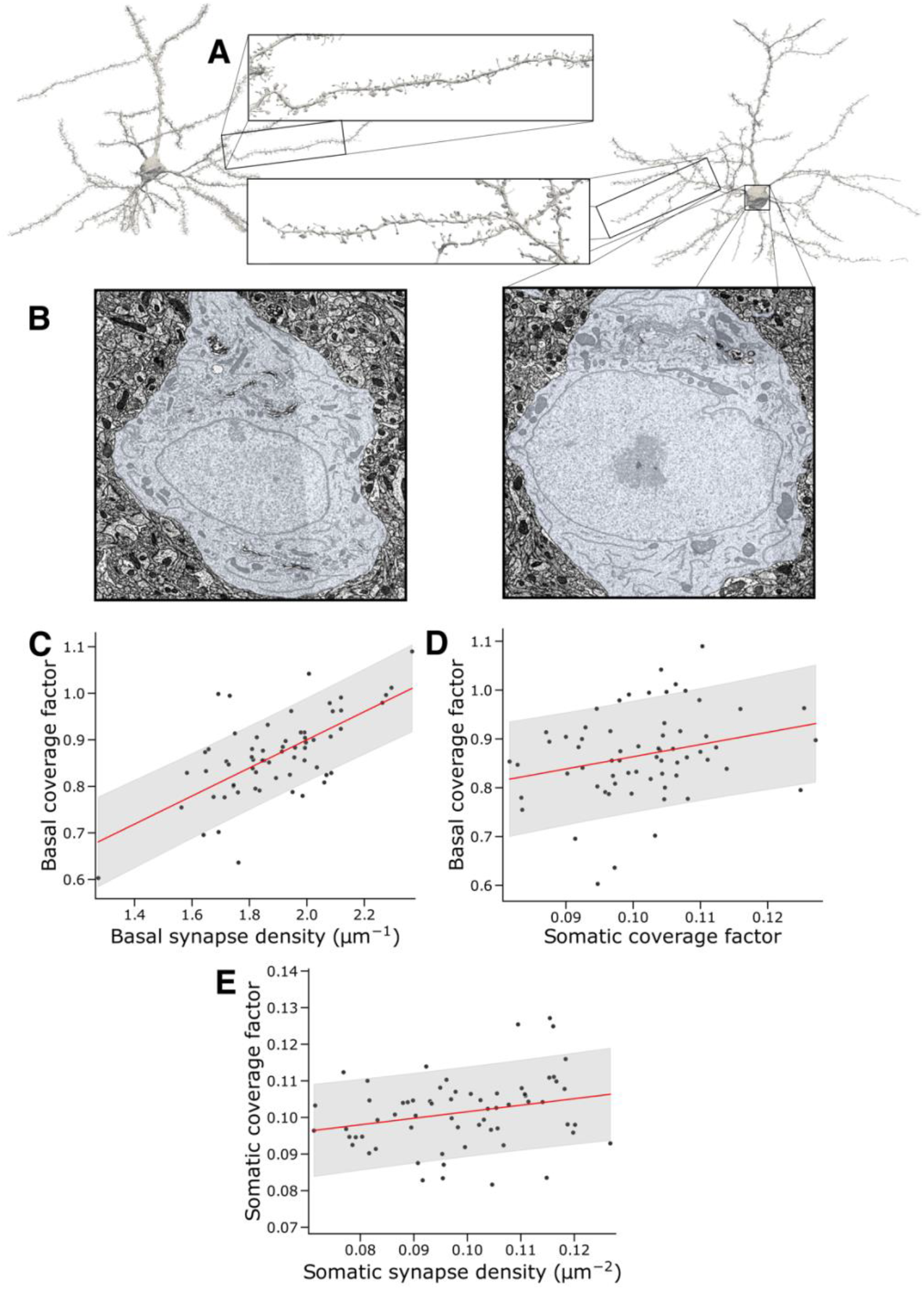
Outlier cell of mitochondria analysis, Related to Figure 4. We can only speculate as to whether this cell is unhealthy, a rare pyramidal type, or a developmental anomaly. (A) Outlier cell (right) spine density compared to PyC with the most similar basal coverage (left). (B) Outlier cell somatic mitochondria (right) compared to PyC with the most similar somatic synapse density (left). (C) Basal coverage and basal synapse density w/ outlier removed (Pearson’s r=0.654, p<4.6e-9). (D) Basal mitochondrial coverage and somatic mitochondrial density w/ outlier removed. (r=0.268, p=0.032). (E) Somatic synapse density and somatic mitochondrial density w/ outlier removed. (r=0.257, p=0.040). Shaded regions in (C-E) show the 80% prediction interval.

**Data S3.**
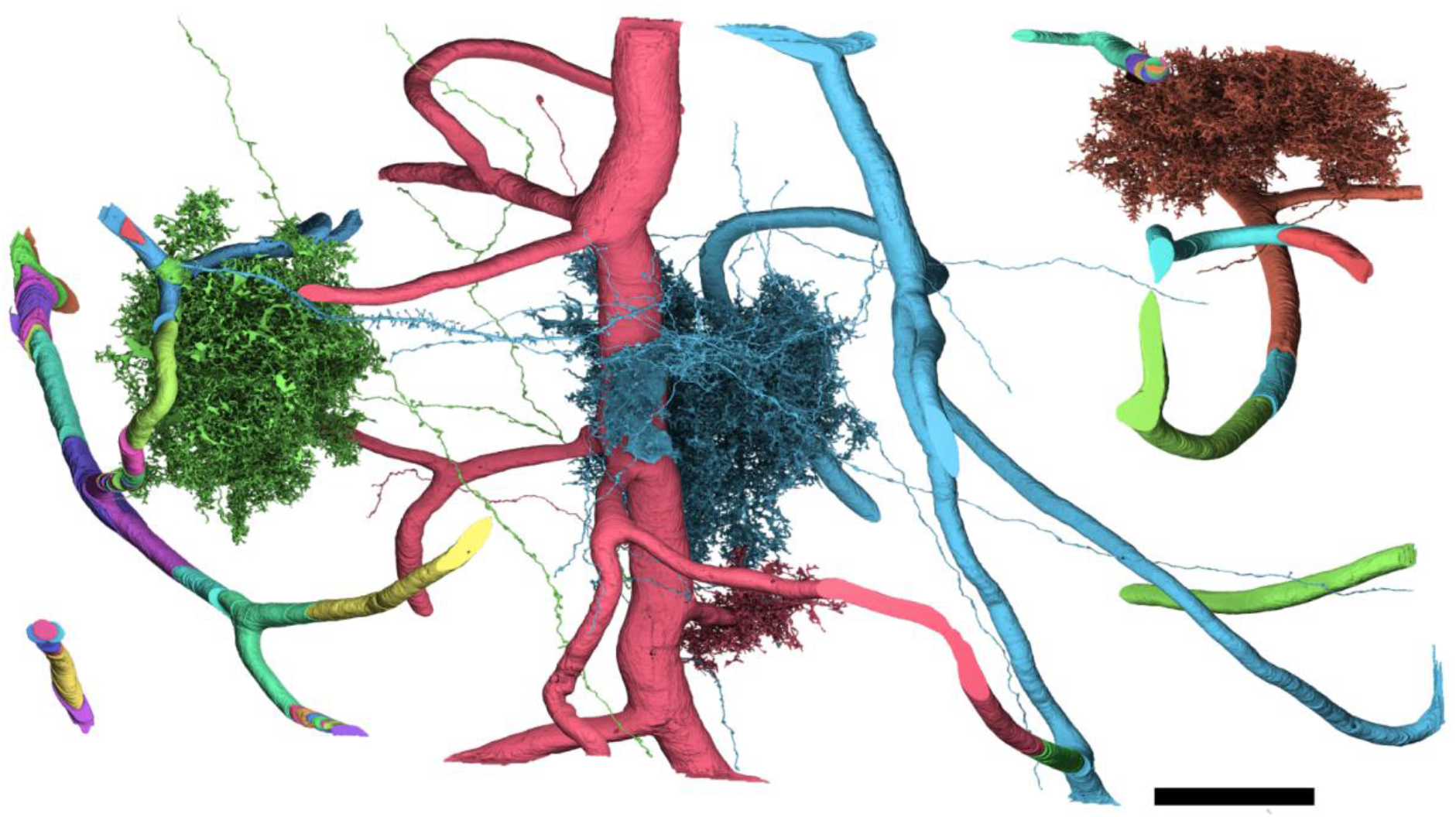
Full vasculature reconstructions with merge errors, Related to Figure 1.

**Data S4.**
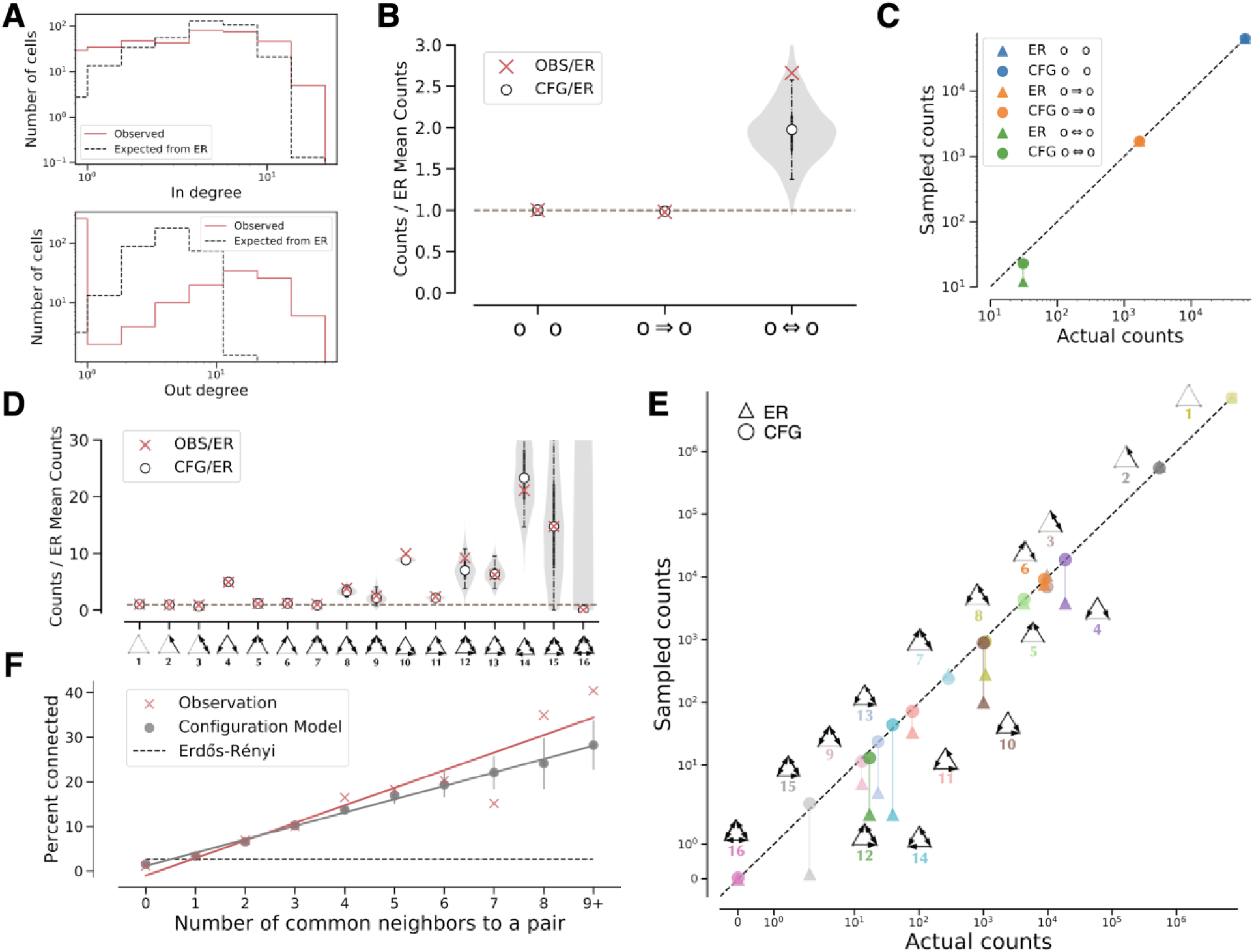
Predicting motif frequencies for all 363 PyCs, Related to Figure 5. (A) In and out degree distributions of all PyCs and the expected distribution of in and out degrees in a regular ER model (edge probability = 0.0113). Red: observed data; Black, dashed: ER model. Histograms are calculated with 8 bins of the same widths on a log scale. The expected degree distributions are estimated from 100 samples from the ER model. (B) 2-cell motif frequencies in the full network and a configuration model relative to the ER model. The shaded region shows the smoothed distribution of motif counts sampled from the configuration model. White points indicate medians, vertical lines indicate quartiles and the 95% confidence interval for 1,000 samples. (C) Comparison of 2-cell motif counts in the ER model and configuration model. Circles indicate mean counts sampled from the configuration model, and triangles indicate mean counts sampled from the ER model. (D) 3-cell motif frequencies in the full network and the configuration model relative to a generalized ER model. The violin plot shows the smoothed distribution of the motif counts sampled from the configuration model. White points indicate medians, and lines indicate the 95% confidence interval for 1,000 samples. (E)Comparison of 3-cell motif counts in the generalized ER model and configuration model. Circles and triangles indicate mean counts sampled from the configuration model and the ER model, respectively. (F) The common neighbor rule is significantly more prominent in the pyramidal cell network than in generalized ER random networks. Gray: configuration model; Black, dashed: generalized ER random networks; Red: observed data. (red slope: r^2^=0.87, p=9.5e-5; gray: r^2^≈1.00, p=8.2e–11; z-test). Lines indicate standard deviation of 100 samples.

**Data S5.**
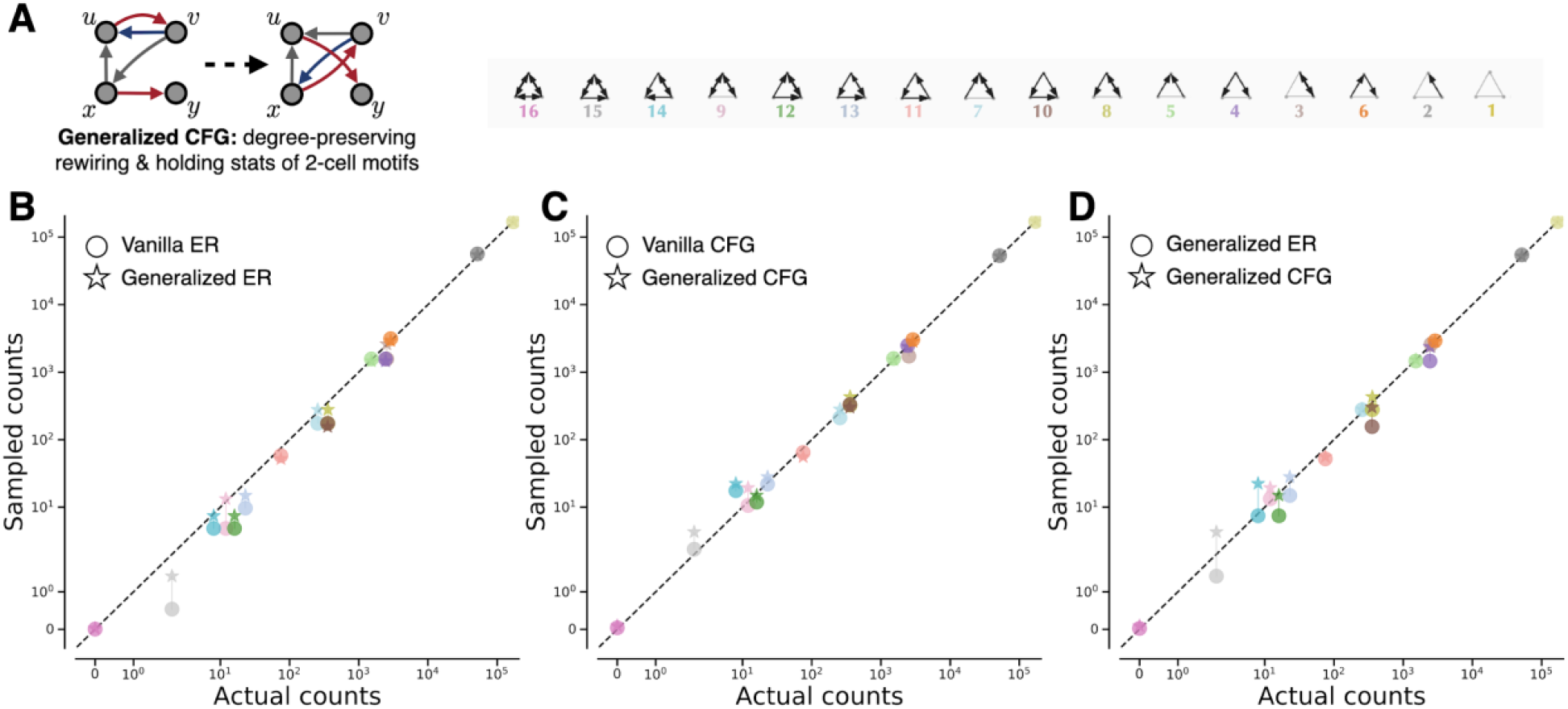
Comparison of the Erdős–Rényi model, configuration model, generalized Erdős–Rényi model, and generalized configuration model for predicting three-cell motifs, Related to Figure 5. (A) Illustration of sampling from a generalized configuration model, where the statistics of two-neuron motifs in the original network are preserved. (B) Comparison of 3-cell motif counts in the ER model and generalized ER model for the subgraph with the axon length threshold = 100 μm. Circles and stars indicate mean counts sampled from the ER model and the generalized ER model, respectively. (C) Similar to (B), circles and stars indicate mean counts sampled from the configuration model and the generalized configuration model, respectively. (D) Similar to (C), circles and stars indicate mean counts sampled from the generalized ER model and the generalized configuration model, respectively.

**Table S1.**
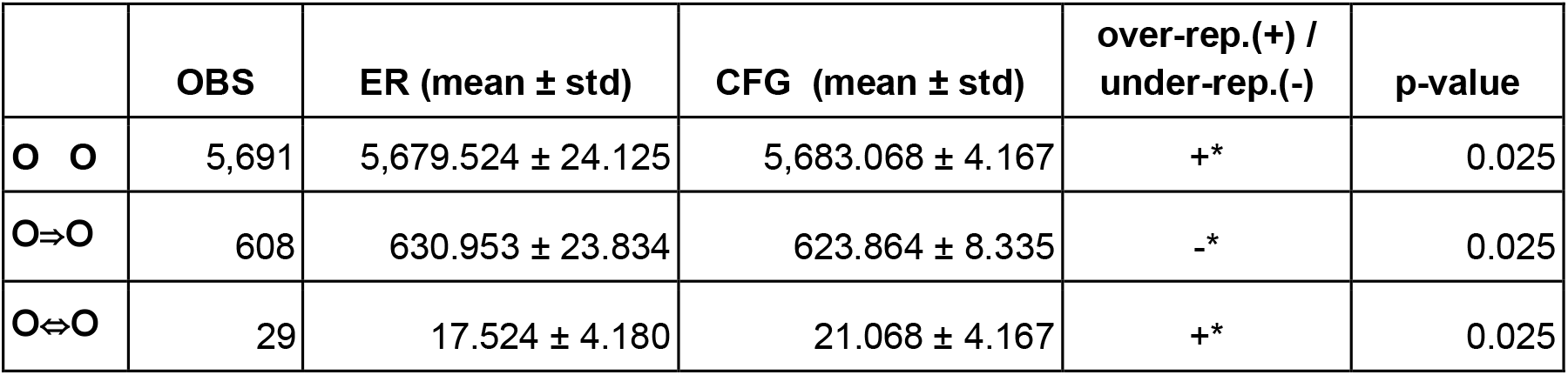
Two-cell motif statistics, Related to Figure 5. Sample size = 1,000. Connection probability for the vanilla Erdős-Rényi model is *p*_*ER*_=0.0526. “+” indicates over-representation, and “-” indicates under-representation with respect to the configuration model. “*” indicates the significant level 0.01 < p ≤ 0.05. The p-value is computed as the percentile of the observed motif counts in the distribution of motif counts sampled from the configuration model.

| | OBS | ER (mean ± std) | CFG (mean ± std) | over-rep.(+) /<br>under-rep.(-) | p-value |
| --- | --- | --- | --- | --- | --- |
| **O O** | 5,691 | 5,679.524 ± 24.125 | 5,683.068 ± 4.167 | +* | 0.025 |
| **O⇒O** | 608 | 630.953 ± 23.834 | 623.864 ± 8.335 | -* | 0.025 |
| **O⇔O** | 29 | 17.524 ± 4.180 | 21.068 ± 4.167 | +* | 0.025 |

**Table S2.**
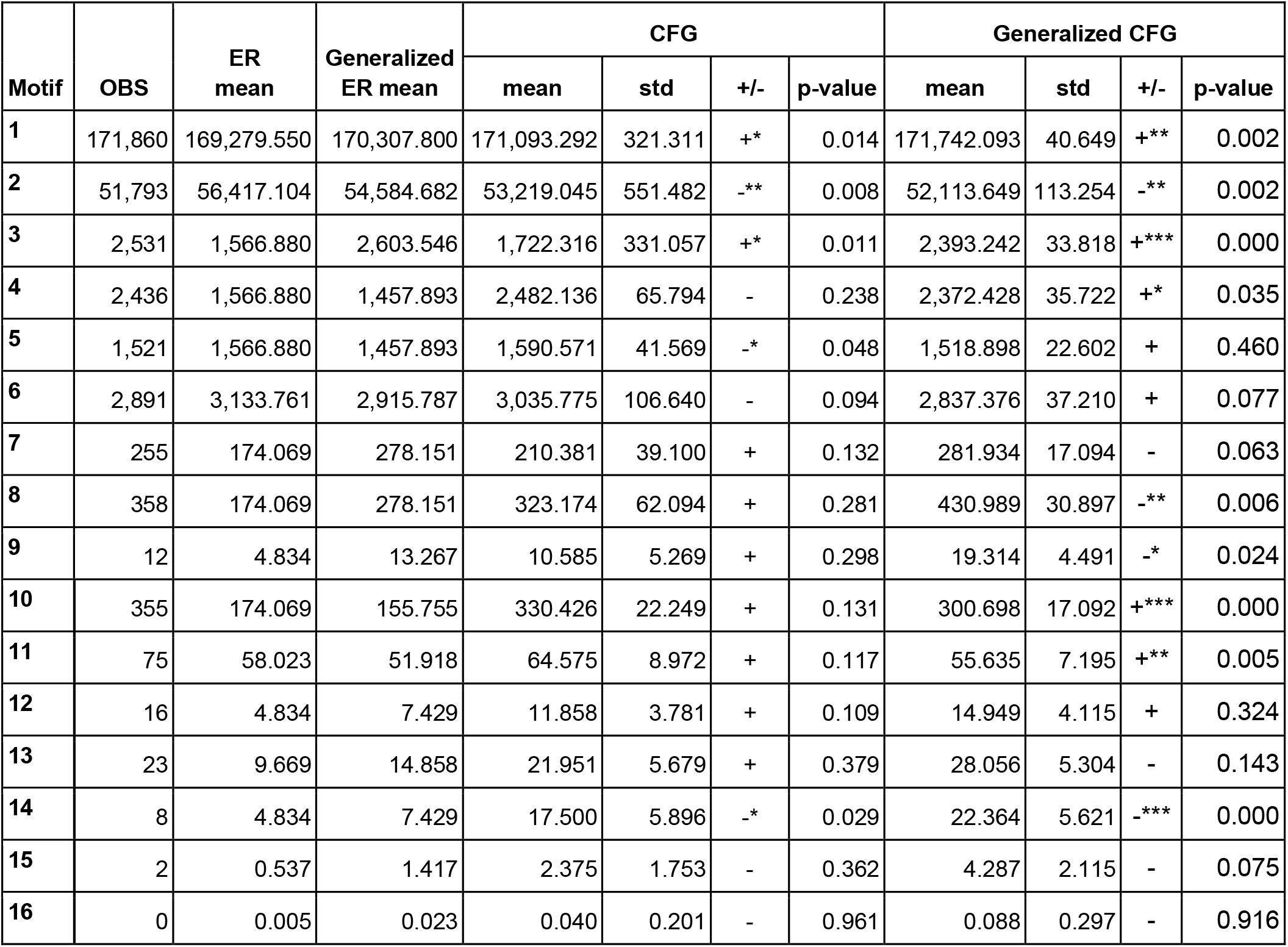
Three-cell motif statistics, Related to Figure 5. Sample size = 1,000. The connection probabilities of the vanilla Erdős-Rényi model is *p* ≈ 5.26 × 10^−2^. Unidirectional and bidirectional connection probabilities in the generalized Erdős-Rényi model are *p*_*uni*_ ≈ 4.80 × 10^−2^and *p*_*ER*_ ≈ 4.58 × 10^−3^. “+” indicates over-representation, and “-” indicates under-representation with respect to the configuration model. “*”, “**”, and “***” indicate the significance levels 0.01 < p ≤ 0.05, 0.001 < p ≤ 0.01, and p ≤ 0.001 respectively. The p-values are computed as the percentile of the observed motif counts in the distribution of motif counts sampled from the configuration model and the generalized configuration model.

