## Supplementary material for "Multiscale and multimodal reconstruction of cortical structure and function": Data S1

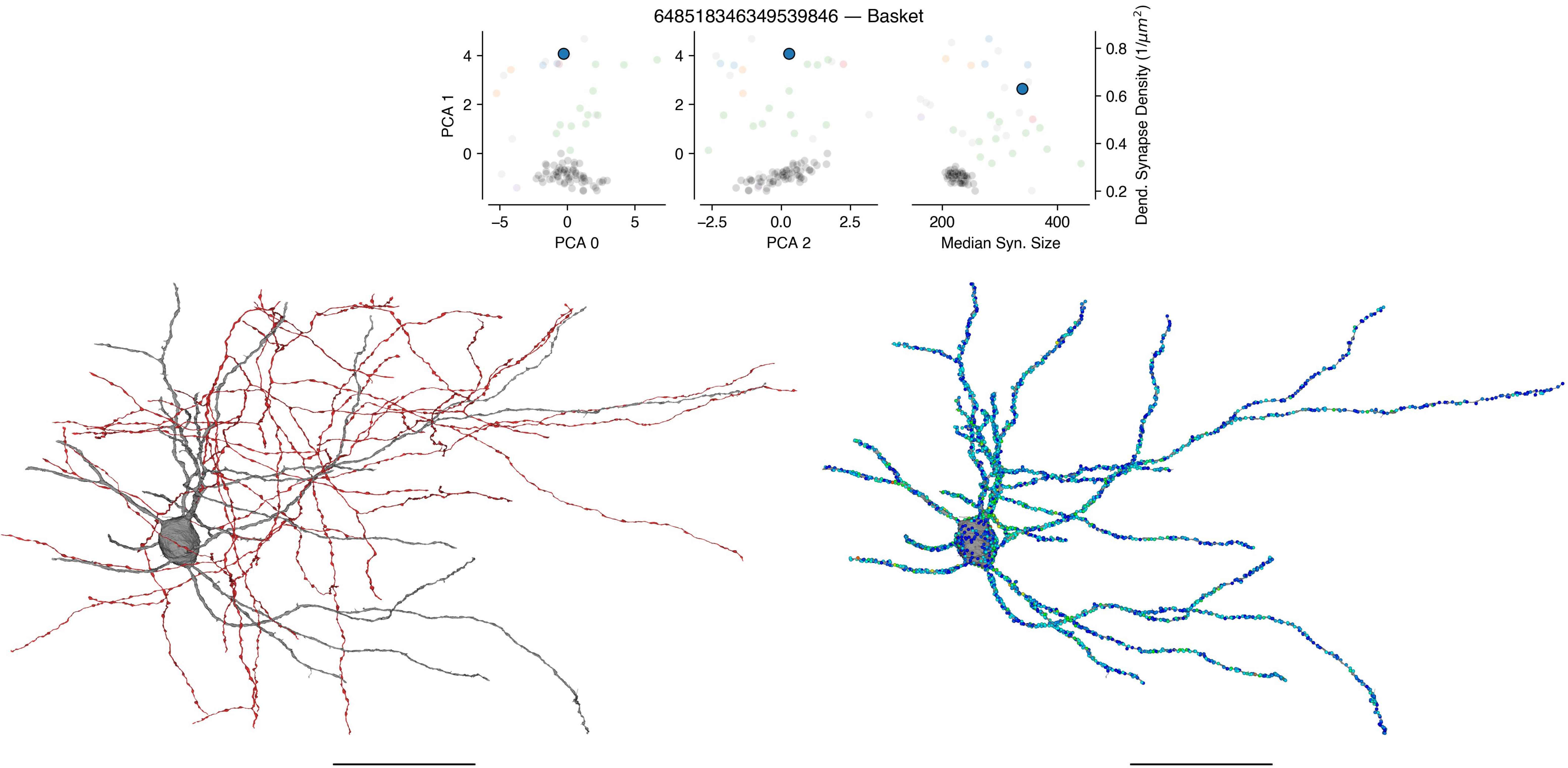
648518346349539846 — Basket
PCA 1
PCA 0
PCA 2
Median Syn. Size
Dend. Synapse Density (1/μm²)

648518346349538791 — Basket

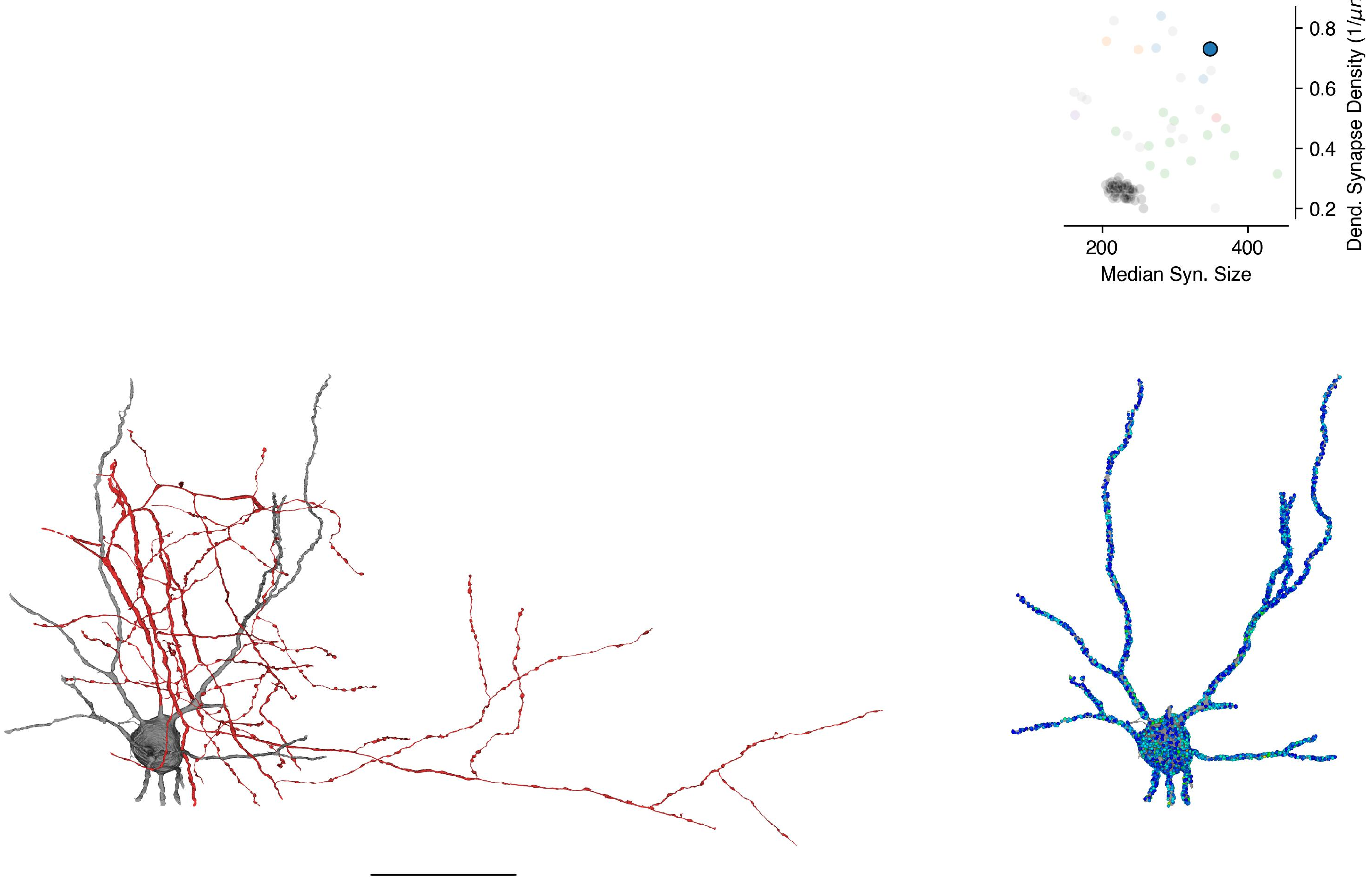

648518346349528994 — Basket

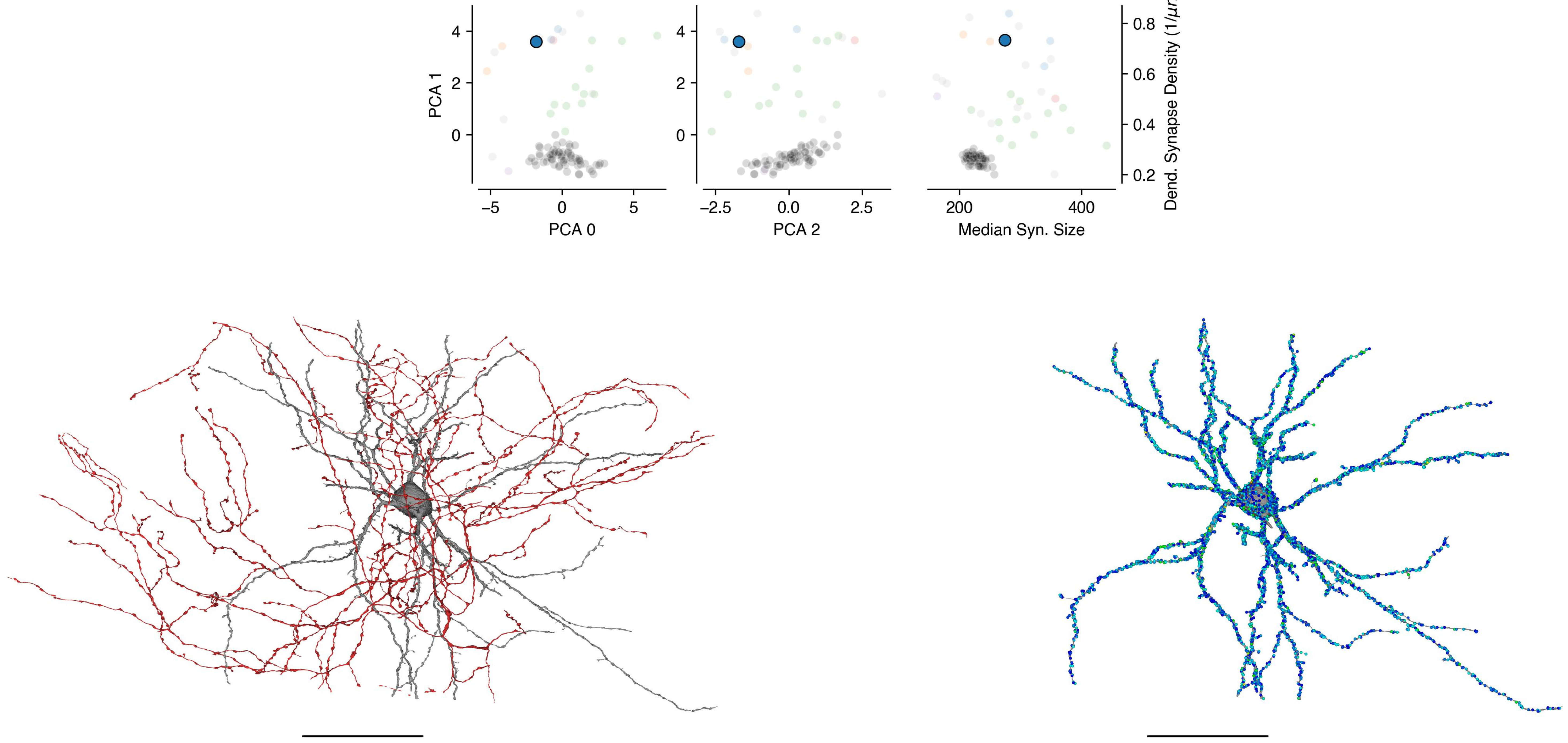

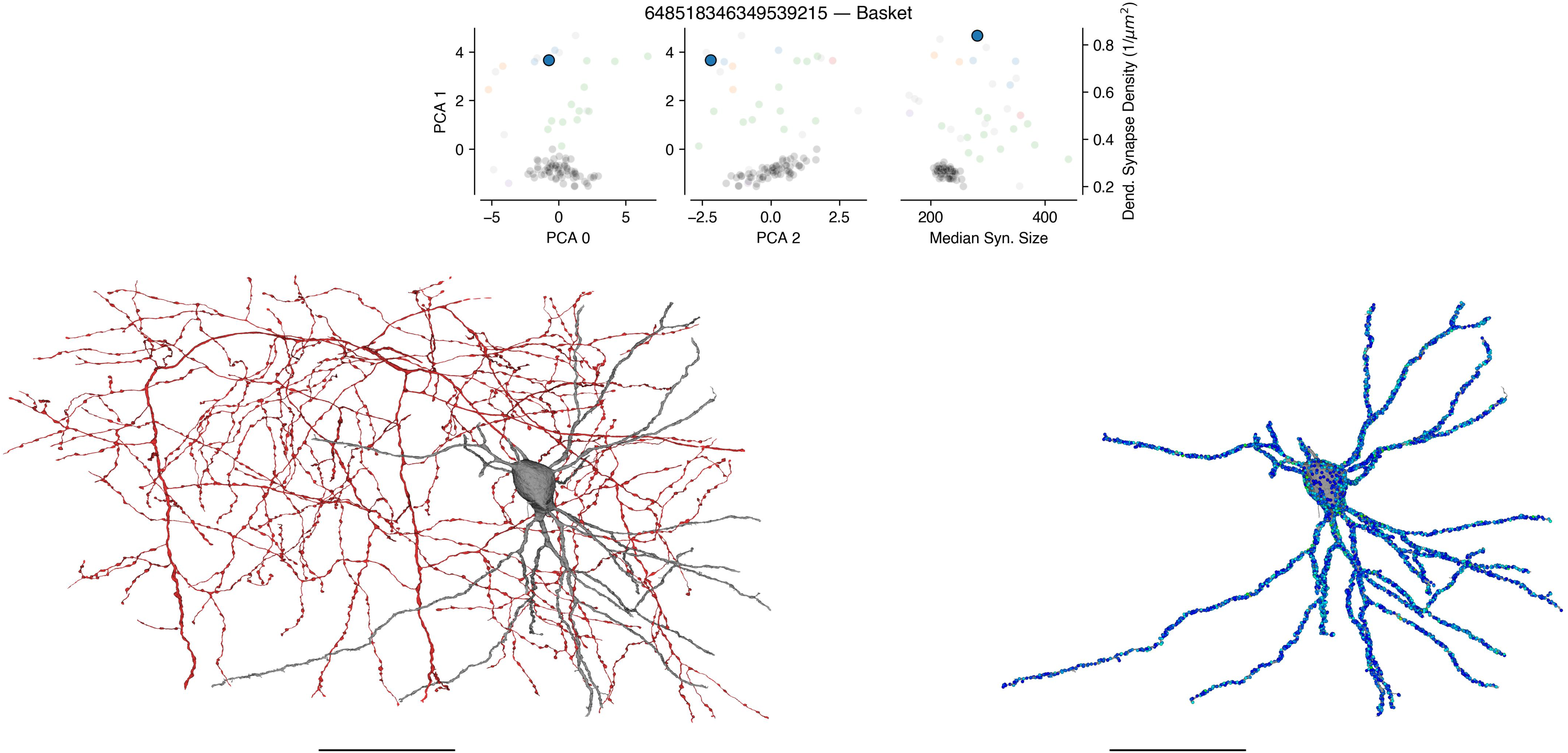

648518346349539215 — Basket
PCA 1
PCA 0
PCA 2
Median Syn. Size
Dend. Synapse Density (1/μm²)

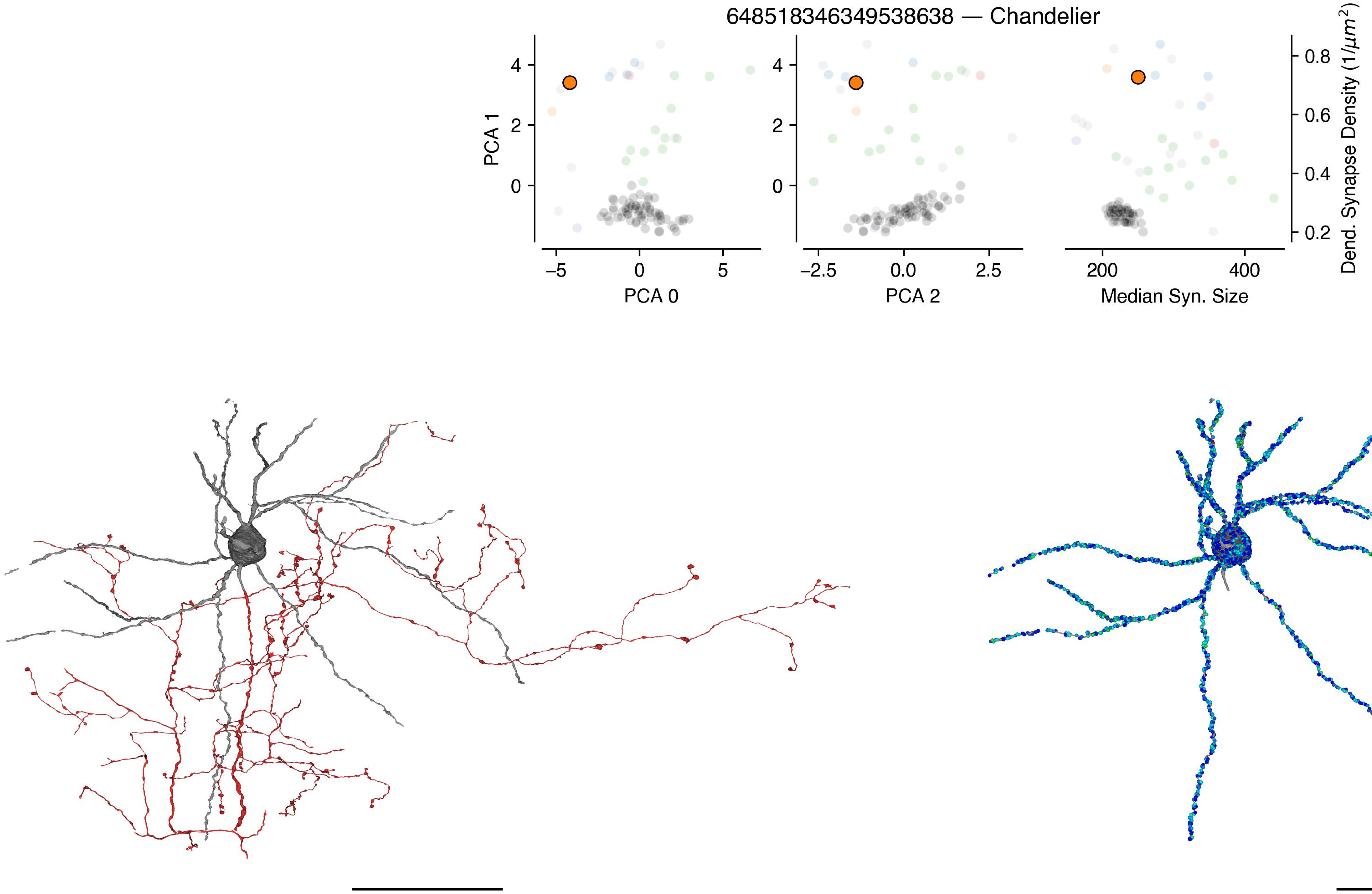

648518346349538638 — Chandelier
PCA 1
PCA 0
PCA 2
Median Syn. Size
Dend. Synapse Density (1/μm²)
4
2
0
–5
0
5
–2.5
0.0
2.5
200
400
0.8
0.6
0.4
0.2

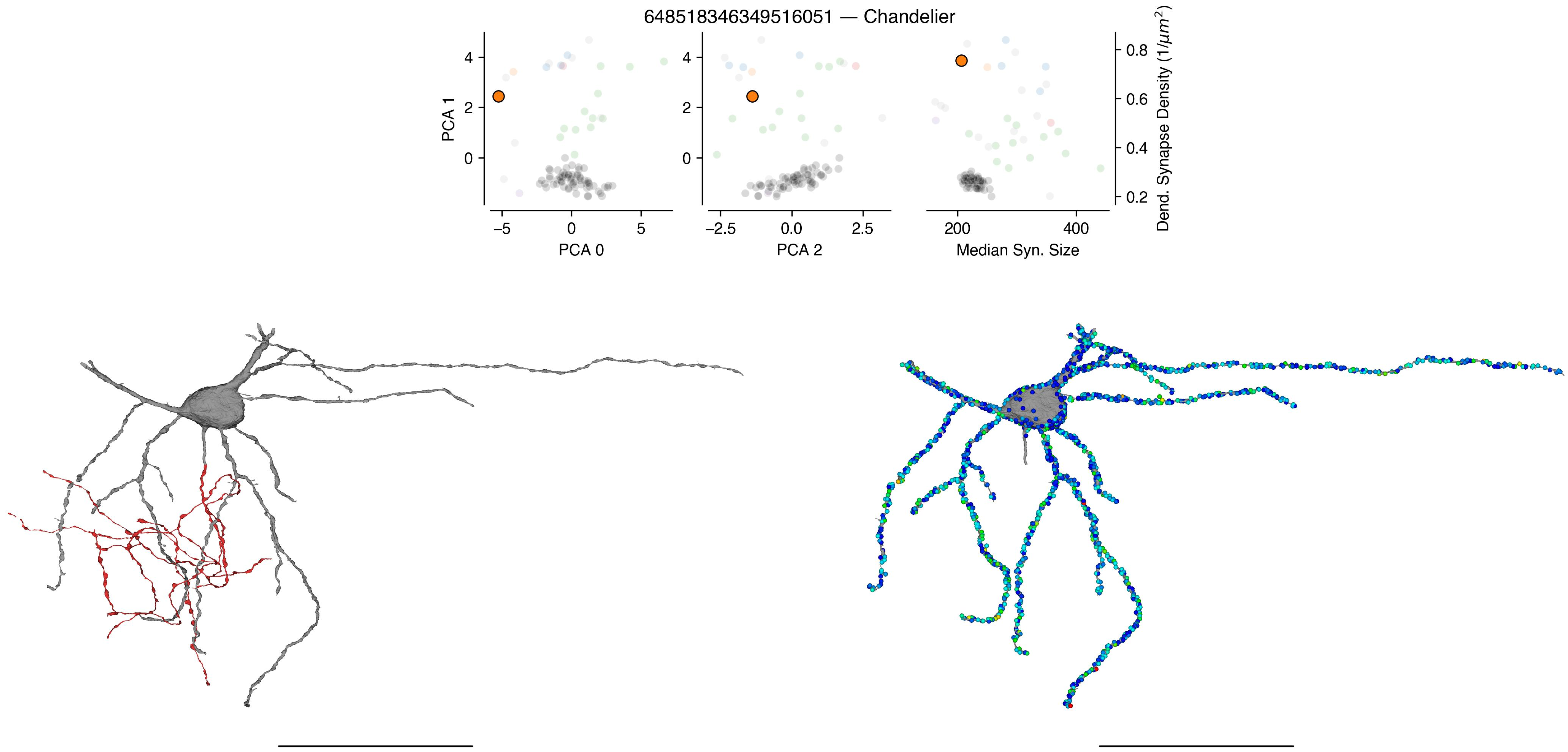
648518346349516051 — Chandelier
PCA 1
PCA 0
PCA 2
Median Syn. Size
Dend. Synapse Density (1/μm²)
−5
0
5
−2.5
0.0
2.5
200
400
0
2
4
0.2
0.4
0.6
0.8

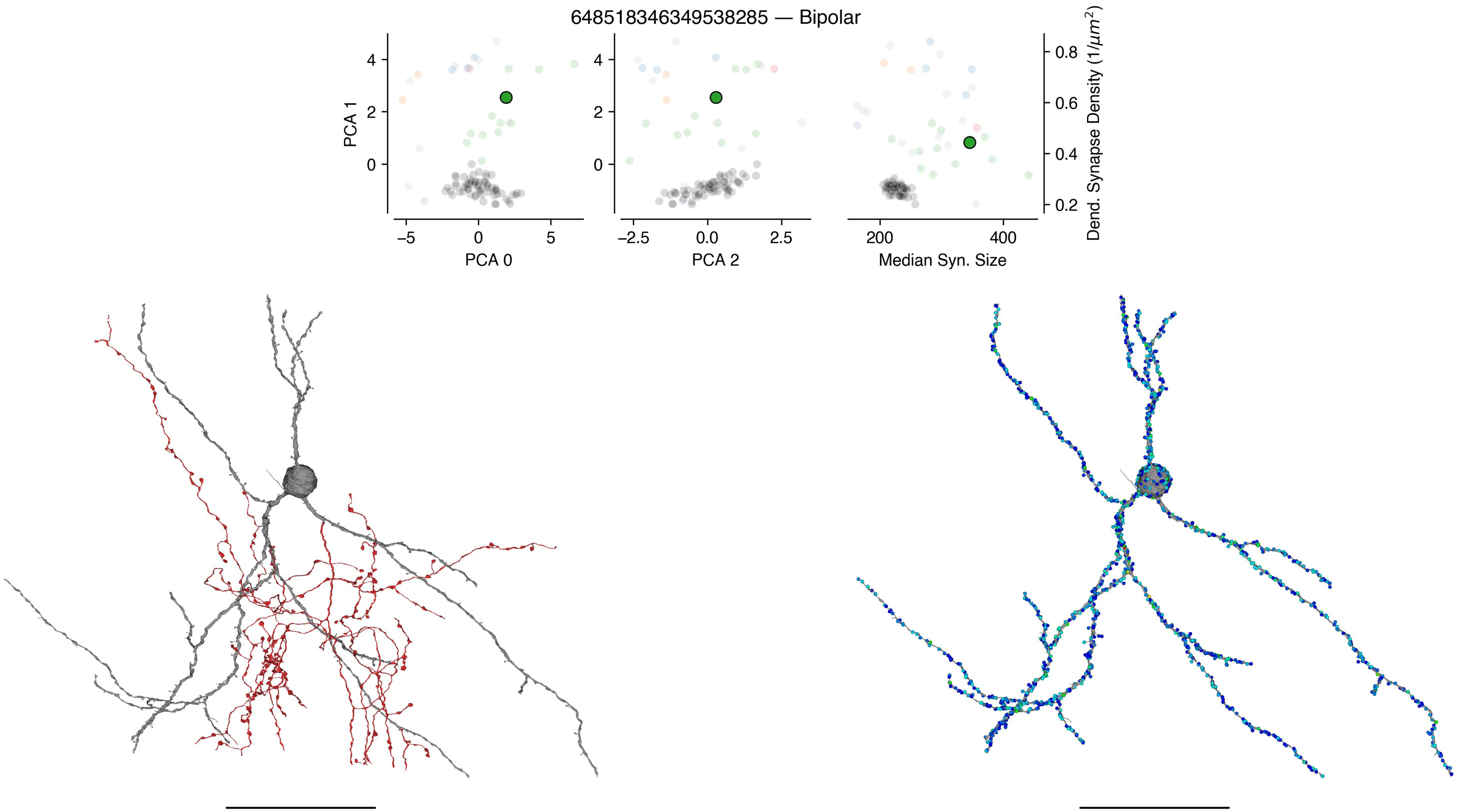
648518346349538285 — Bipolar
PCA 1
4
2
0
–5
0
5
PCA 0
4
2
0
–2.5
0.0
2.5
PCA 2
200
400
Median Syn. Size
0.8
0.6
0.4
0.2
Dend. Synapse Density ($1/\mu m^2$)

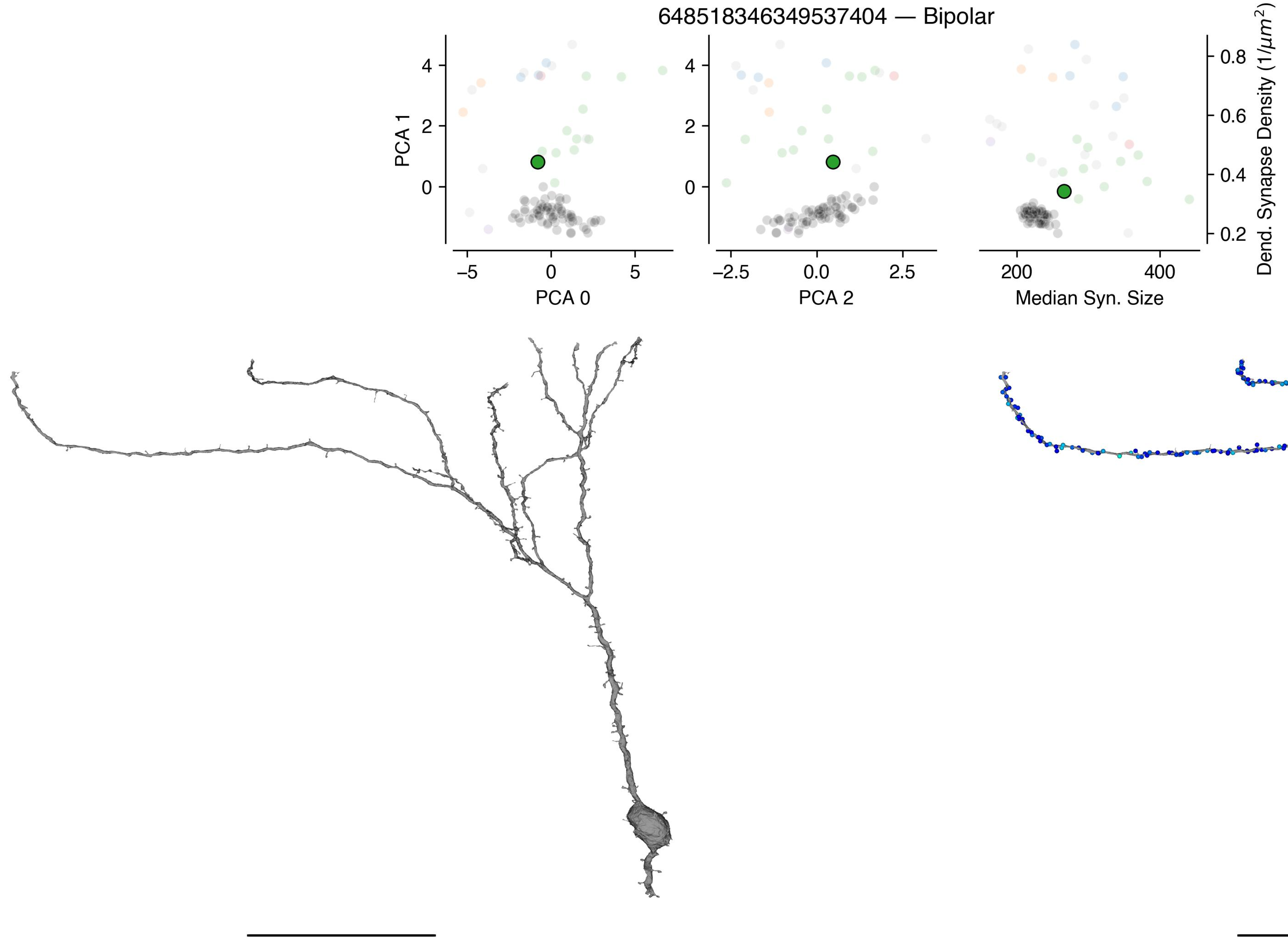

648518346349537404 — Bipolar
PCA 1
PCA 0
PCA 2
Median Syn. Size
Dend. Synapse Density (1/μm²)
4
2
0
–5
0
5
–2.5
0.0
2.5
200
400
0.8
0.6
0.4
0.2

648518346349537389 — Bipolar

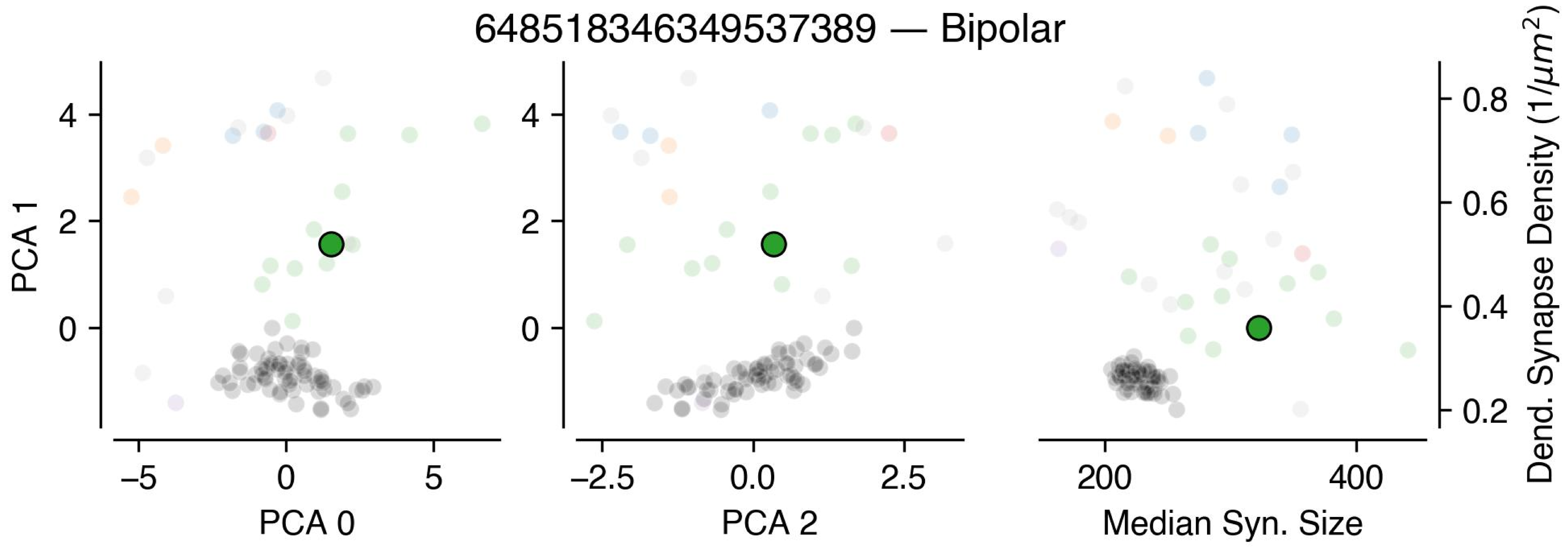


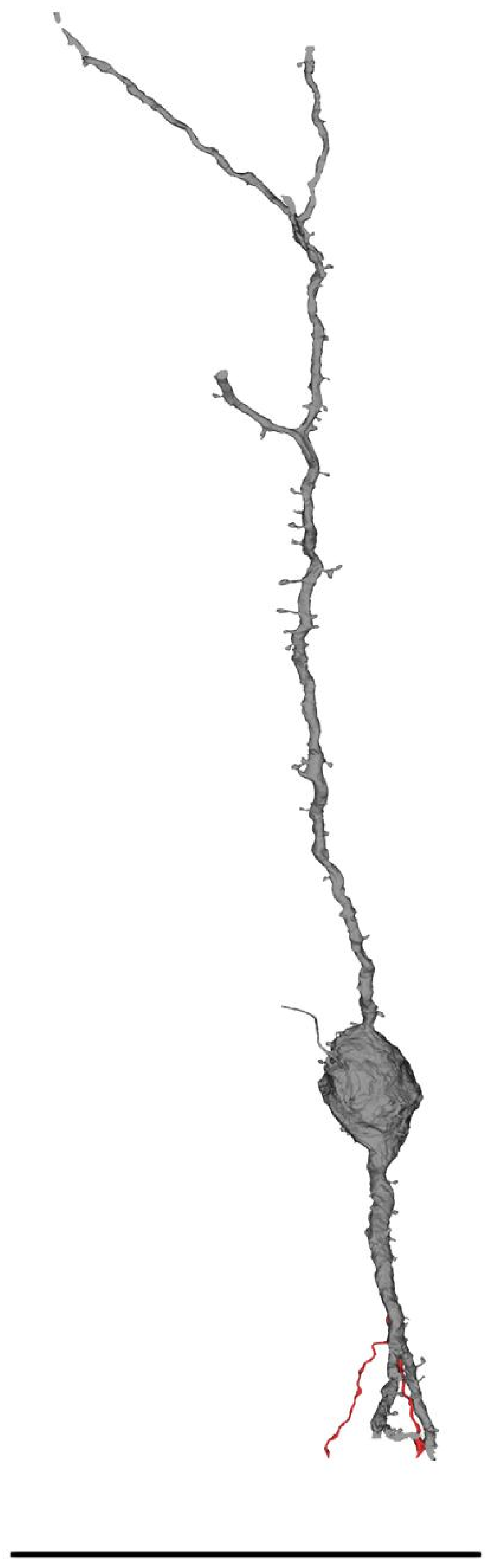

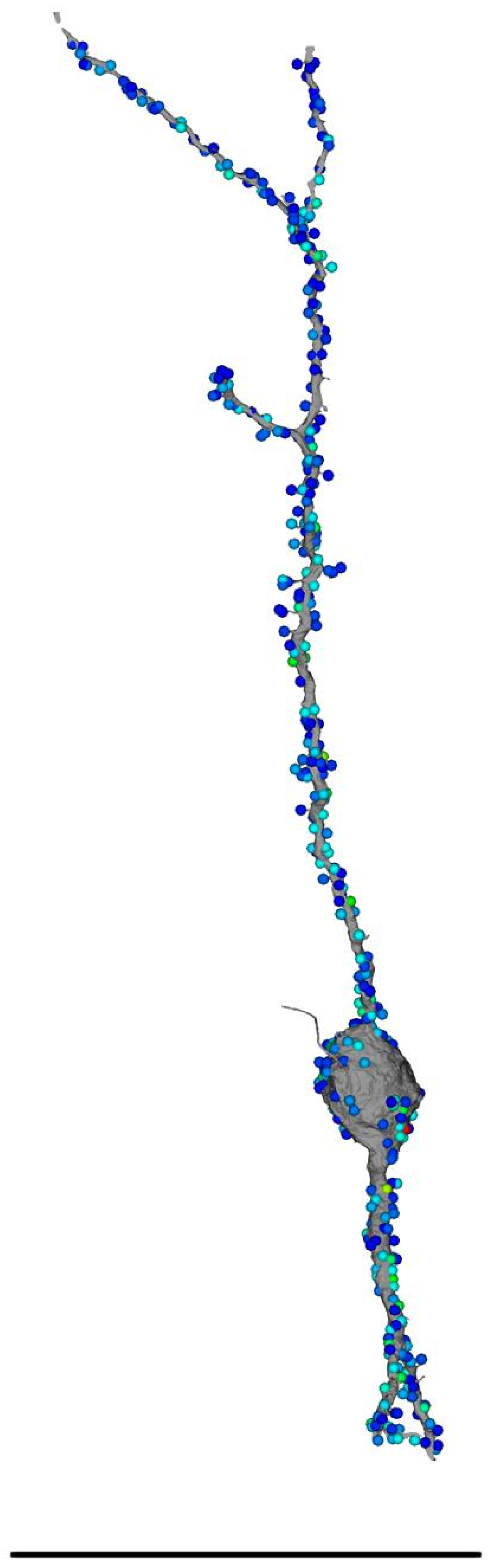

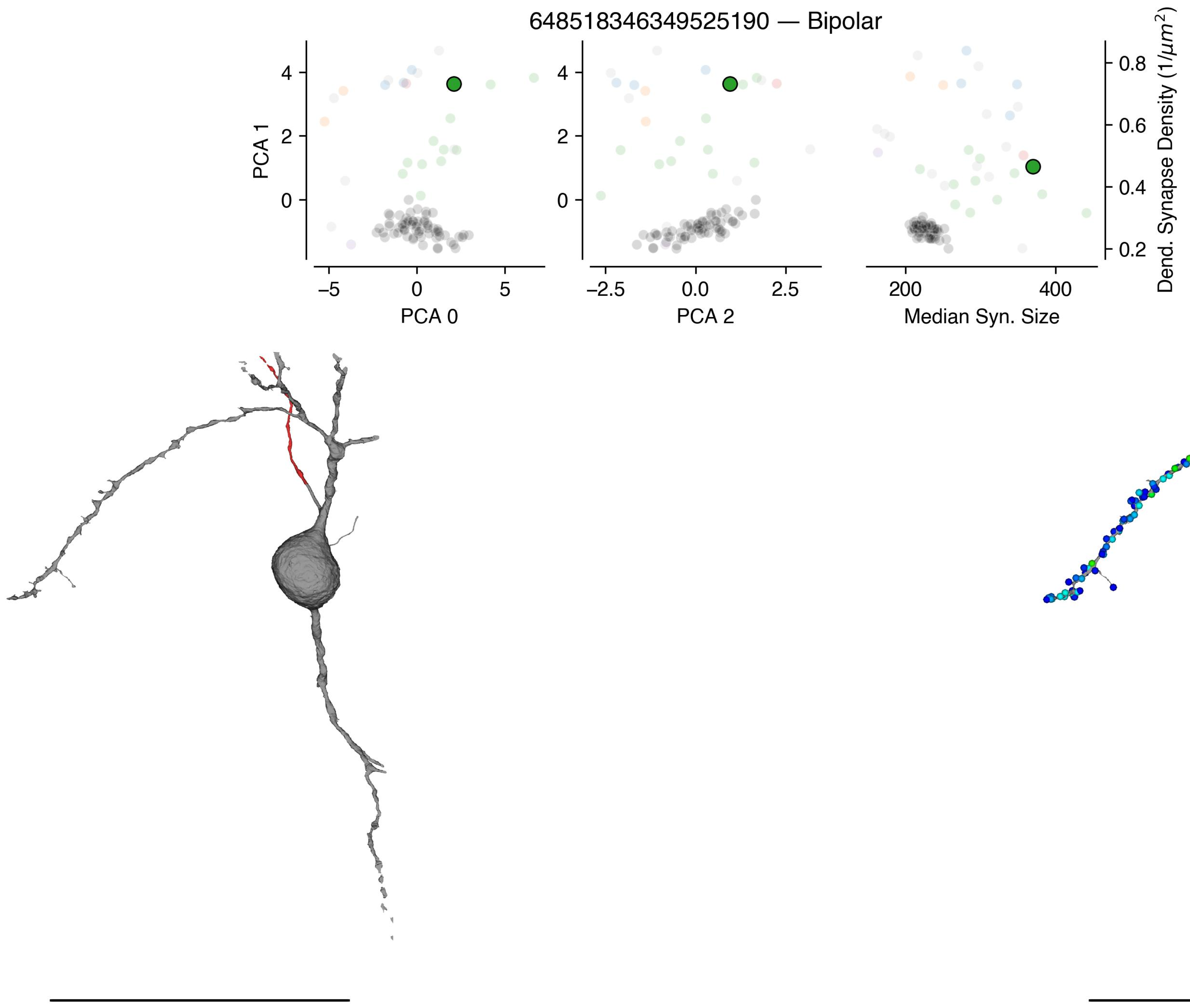
648518346349525190 — Bipolar
PCA 1
PCA 0
PCA 2
Median Syn. Size
Dend. Synapse Density (1/μm²)

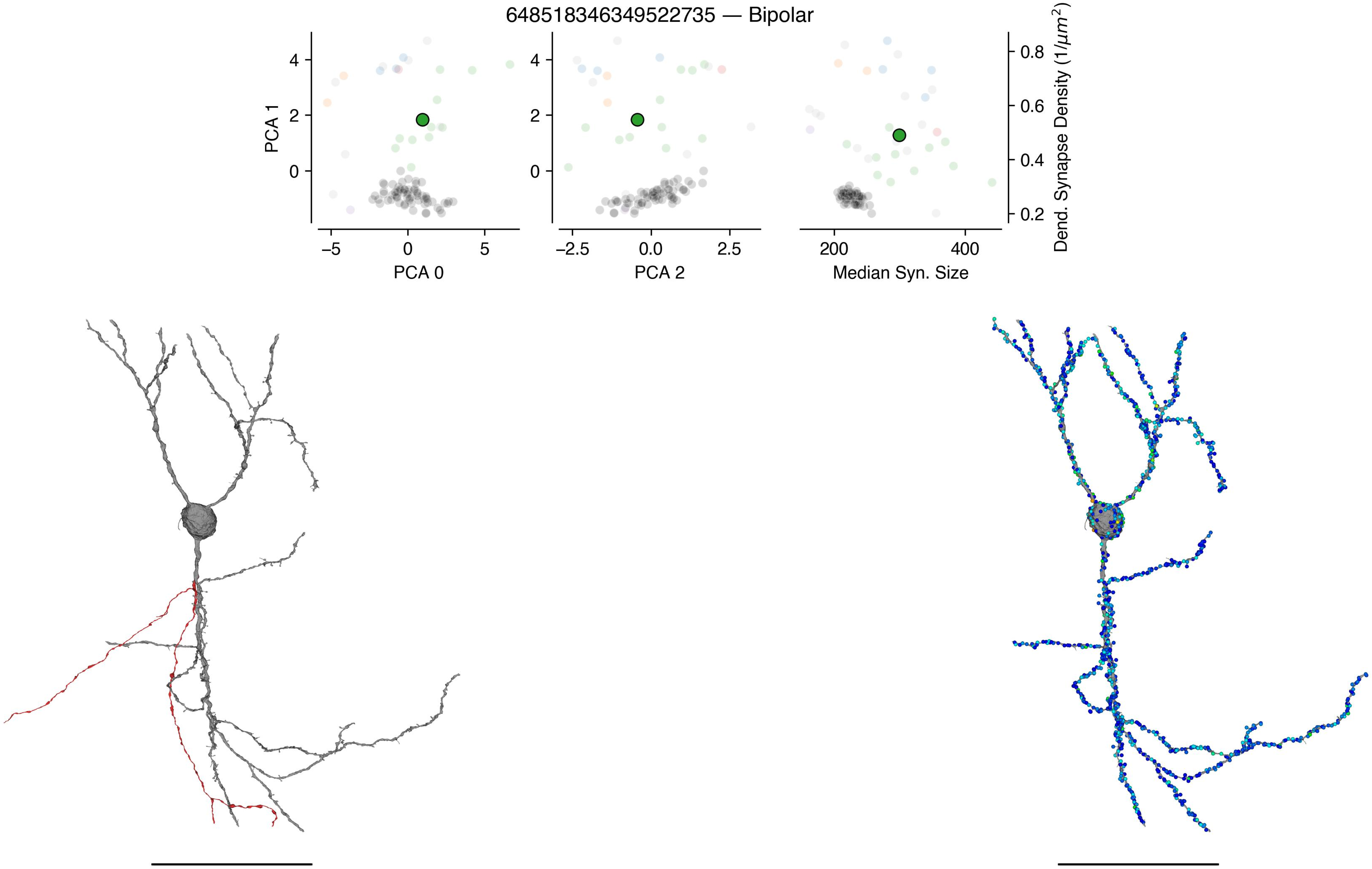
648518346349522735 — Bipolar
PCA 1
PCA 0
PCA 2
Median Syn. Size
Dend. Synapse Density ($1/\mu m^2$)

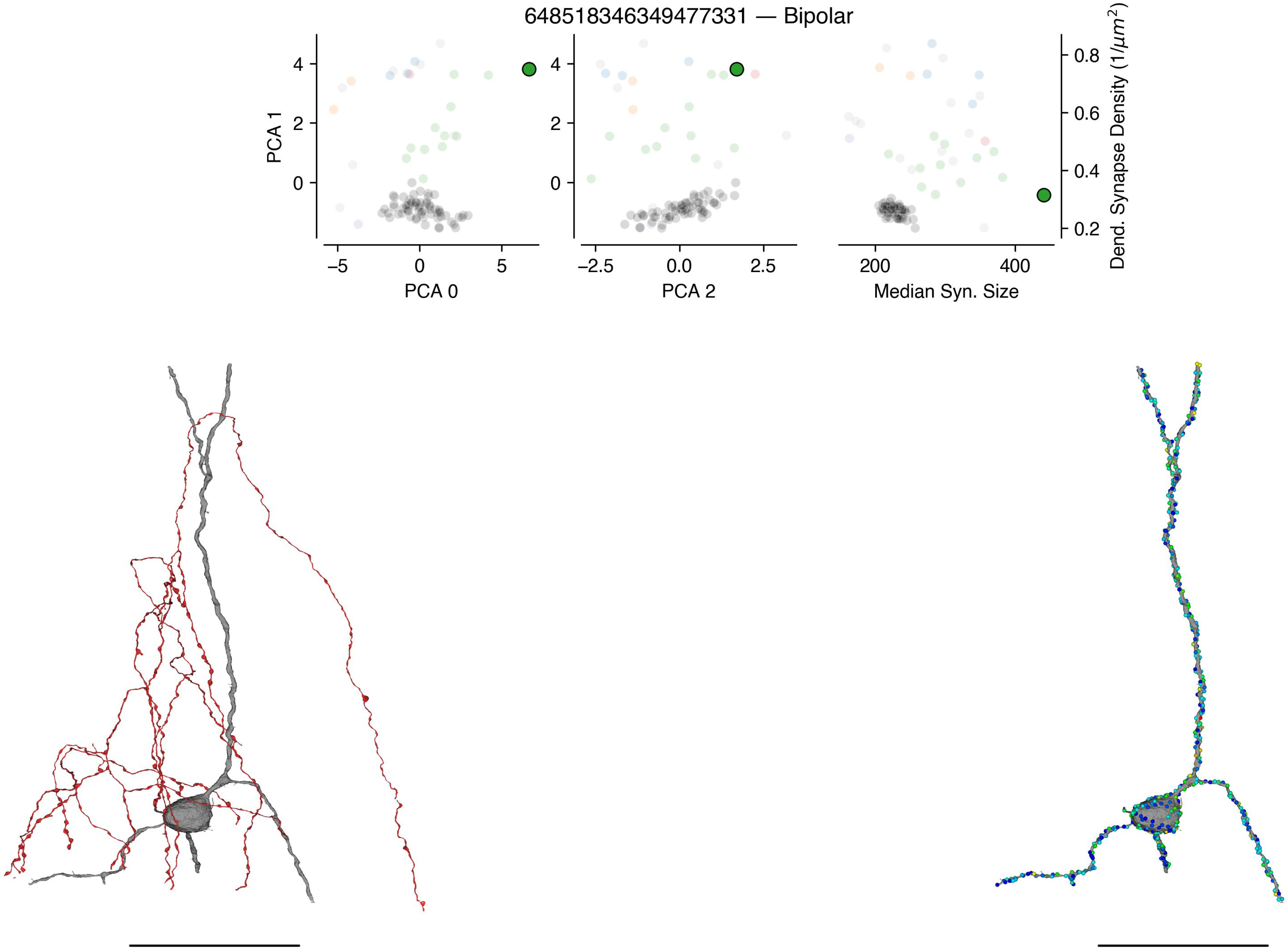
648518346349477331 — Bipolar
PCA 1
PCA 0
PCA 2
Median Syn. Size
Dend. Synapse Density (1/μm²)
4
2
0
−5
0
5
−2.5
0.0
2.5
200
400
0.8
0.6
0.4
0.2

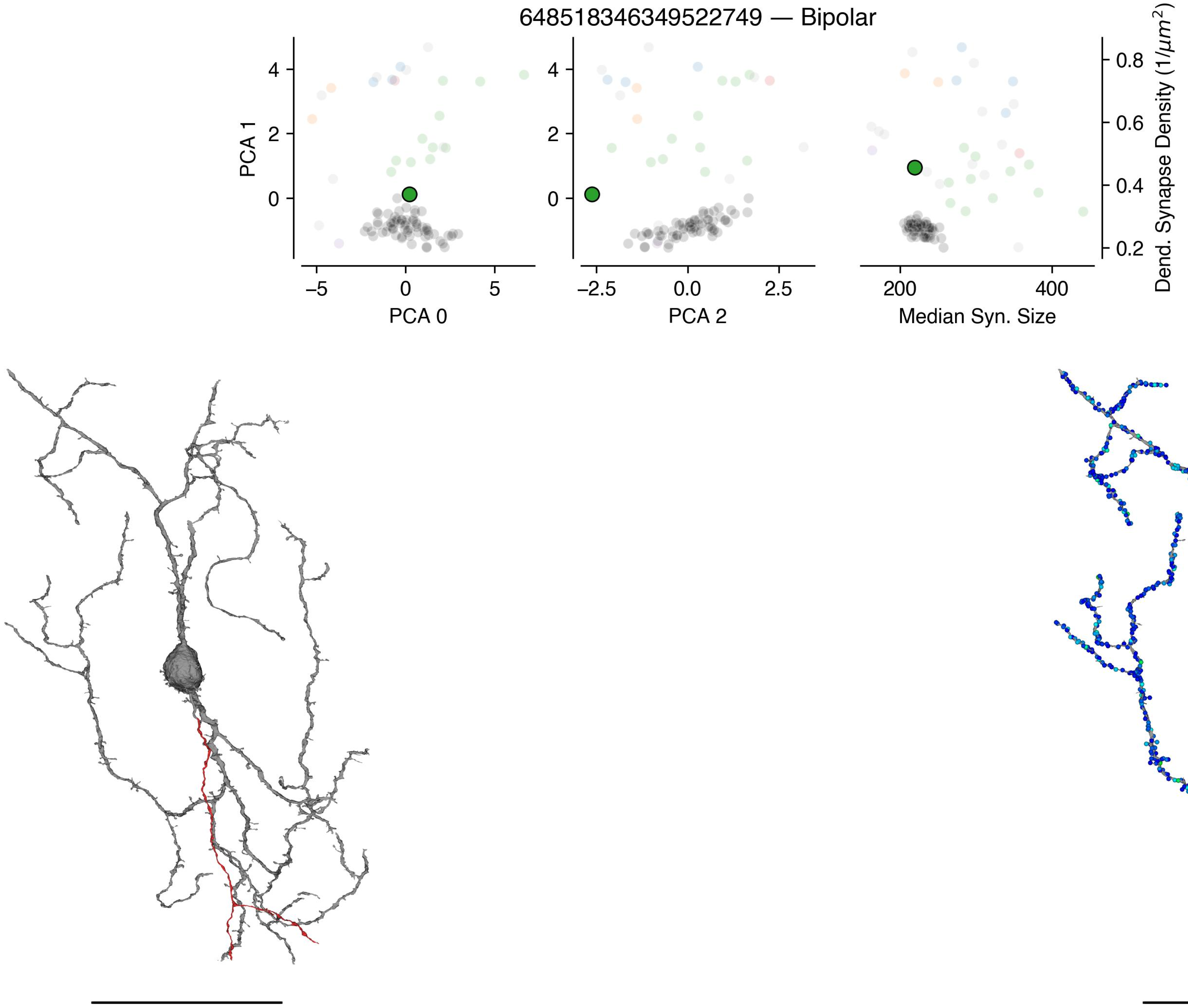

648518346349522749 — Bipolar
PCA 1
PCA 0
PCA 2
Median Syn. Size
Dend. Synapse Density ($1/\mu m^2$)

648518346349487432 — Bipolar

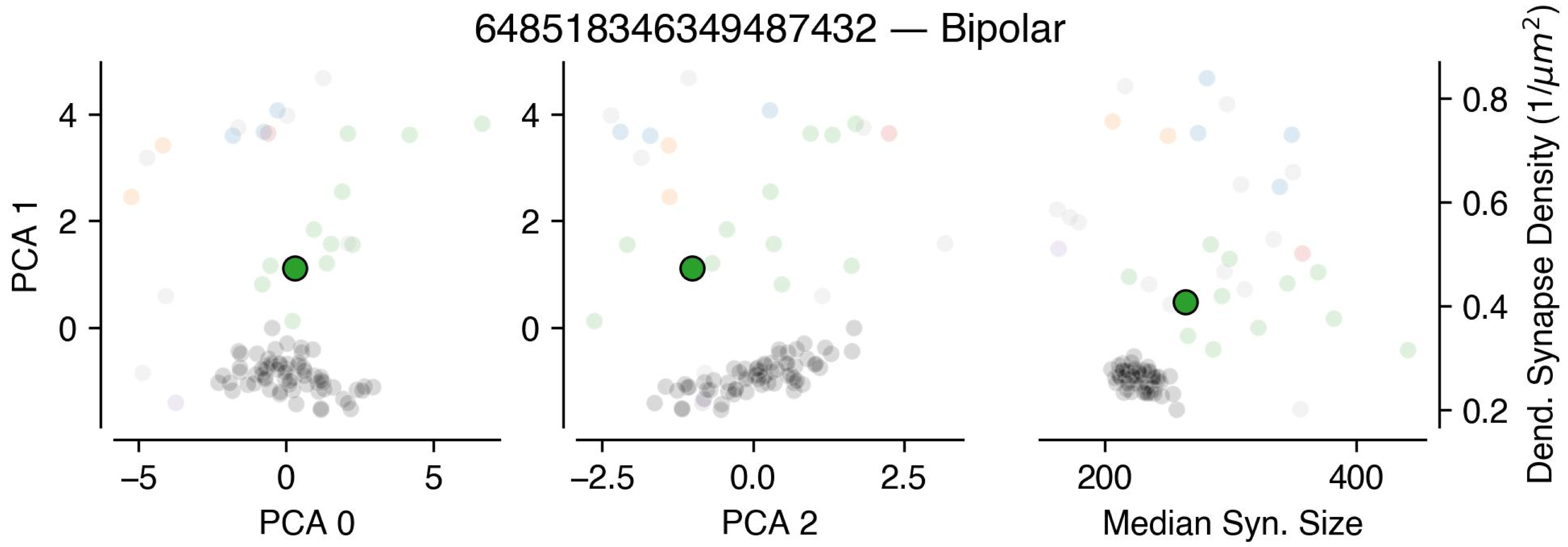


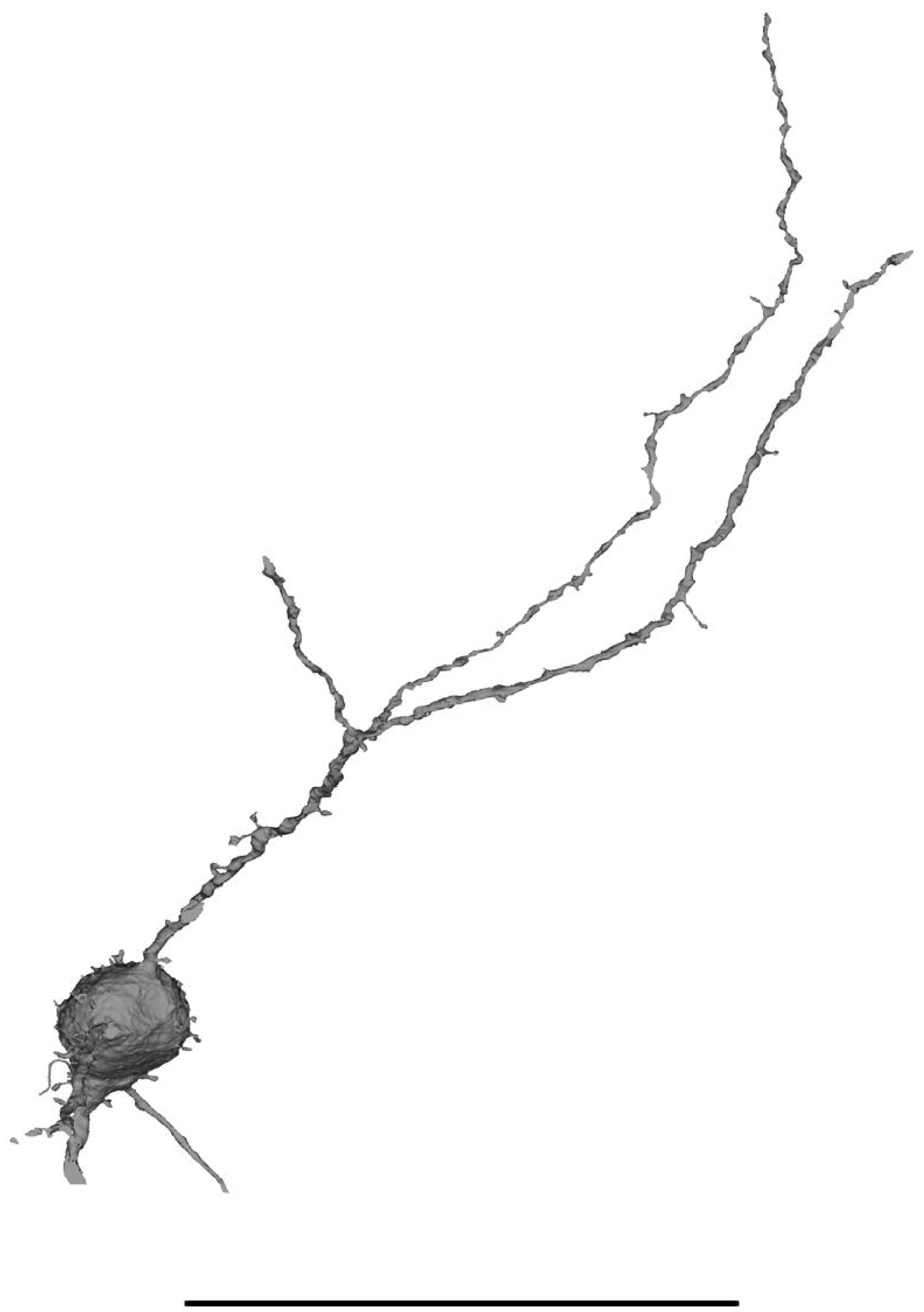

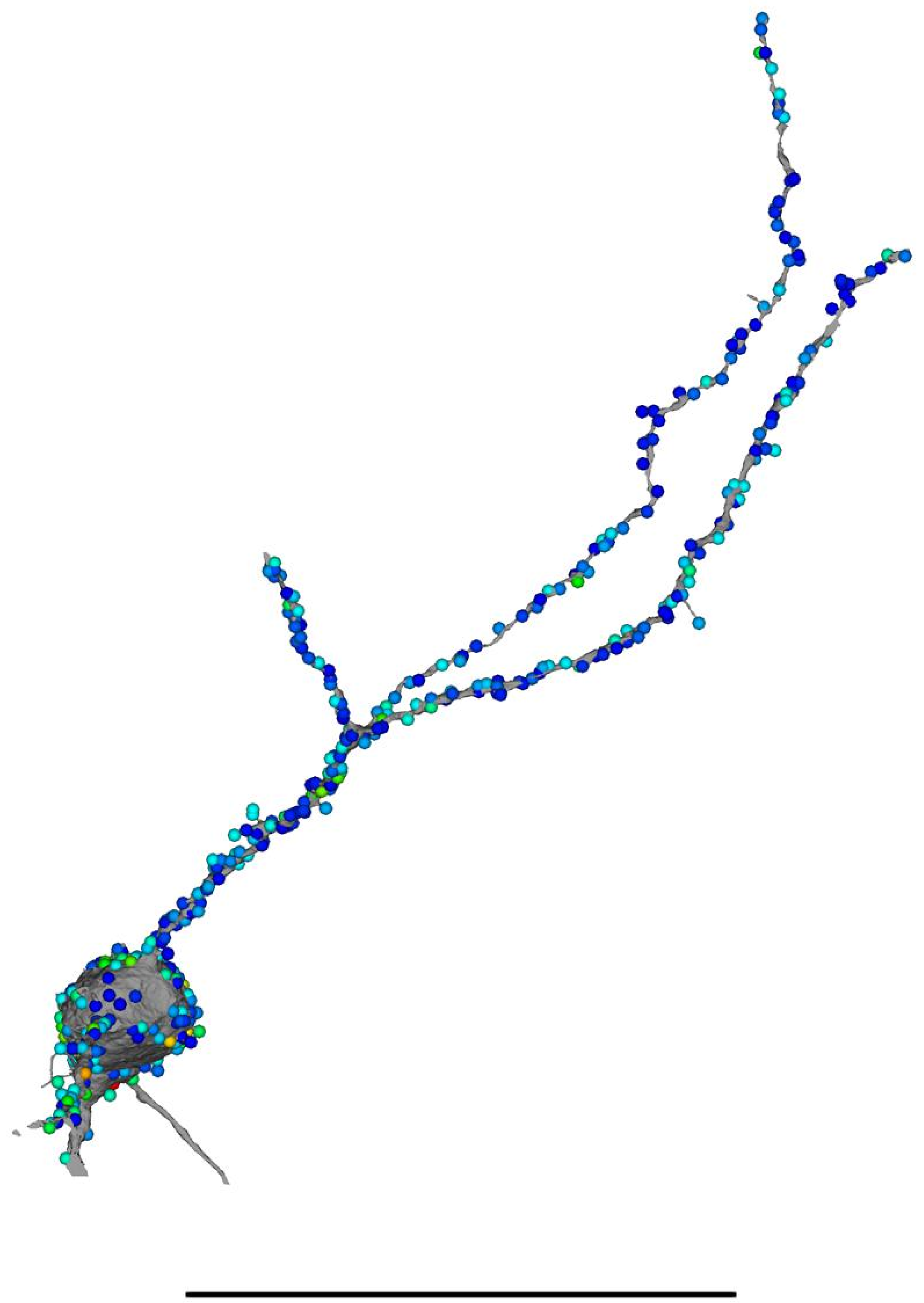

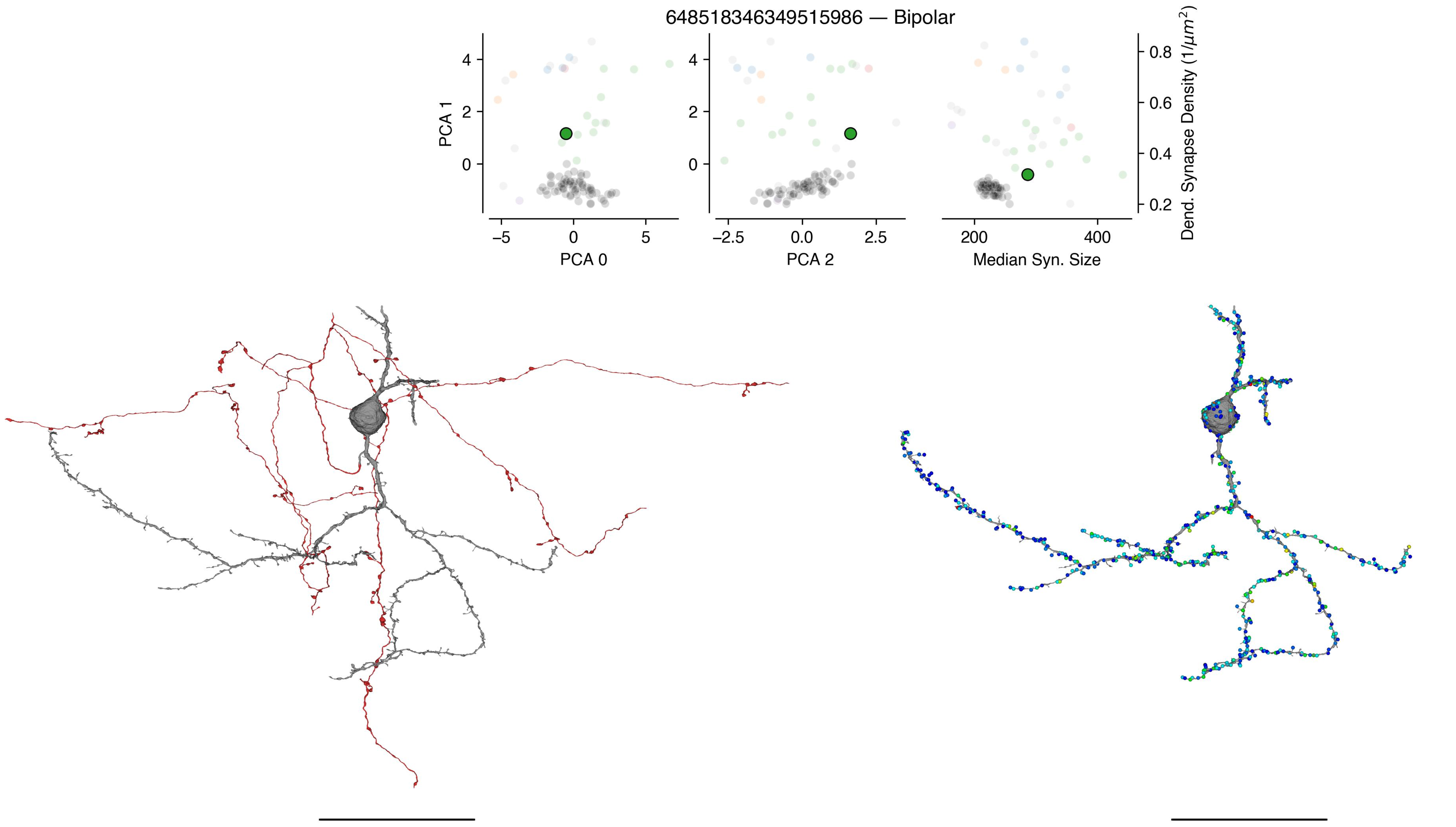

648518346349515986 — Bipolar
PCA 1
PCA 0
PCA 2
Median Syn. Size
Dend. Synapse Density (1/μm²)

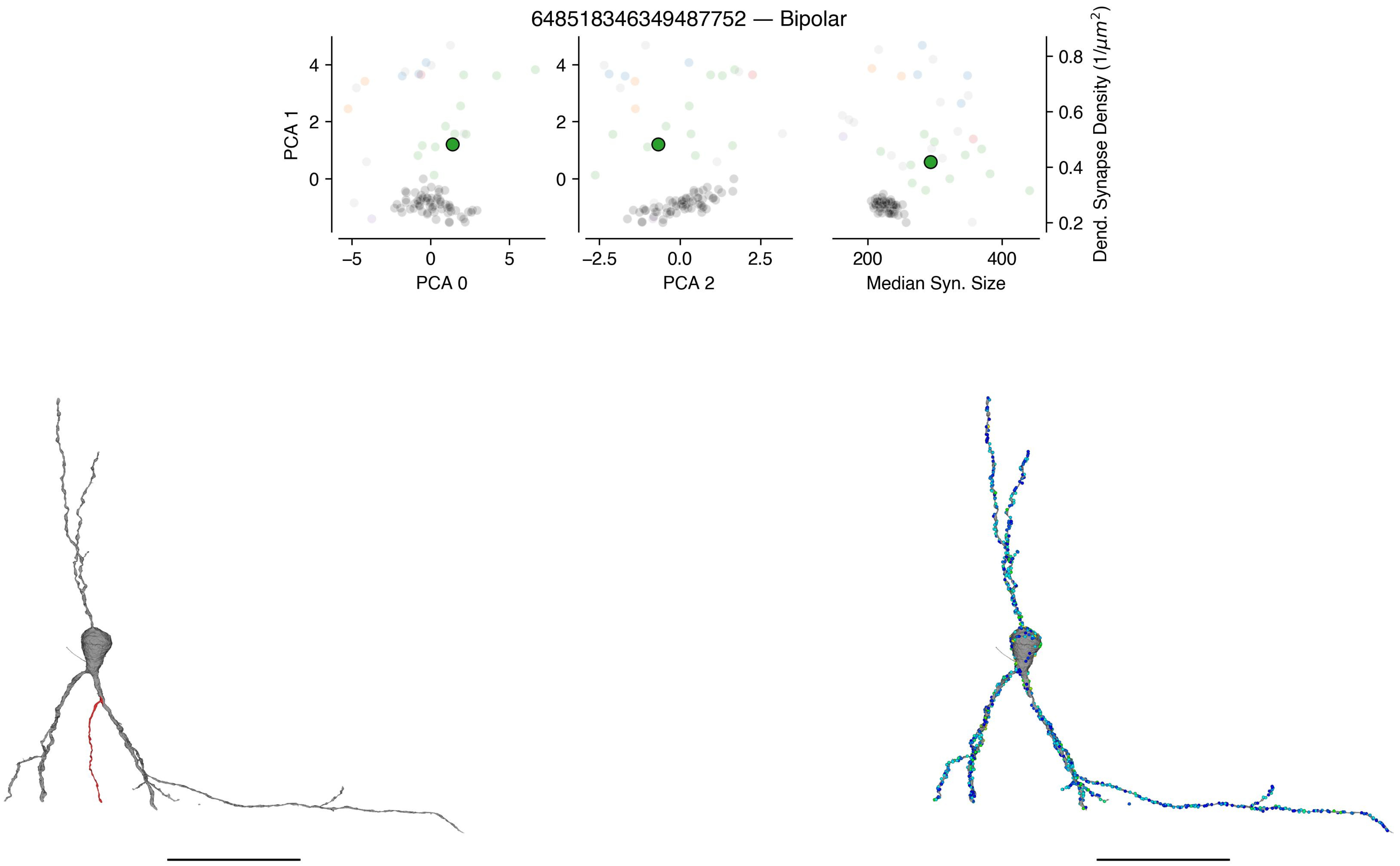
648518346349487752 — Bipolar
PCA 1
PCA 0
PCA 2
Median Syn. Size
Dend. Synapse Density (1/μm²)

648518346349489861 — Bipolar

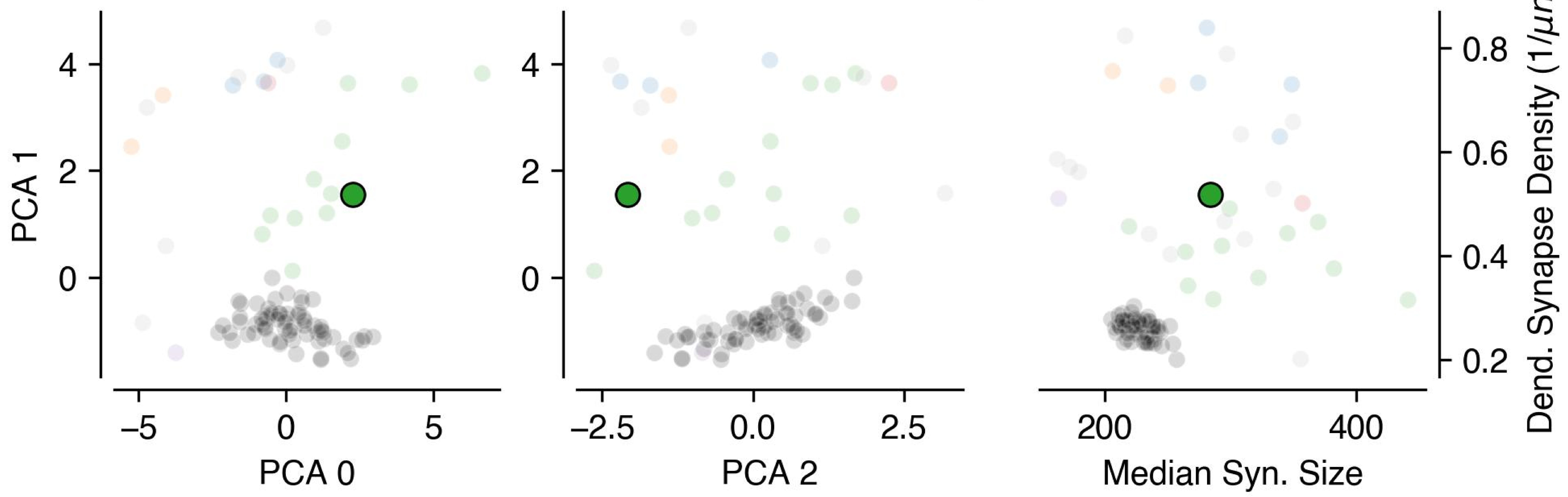


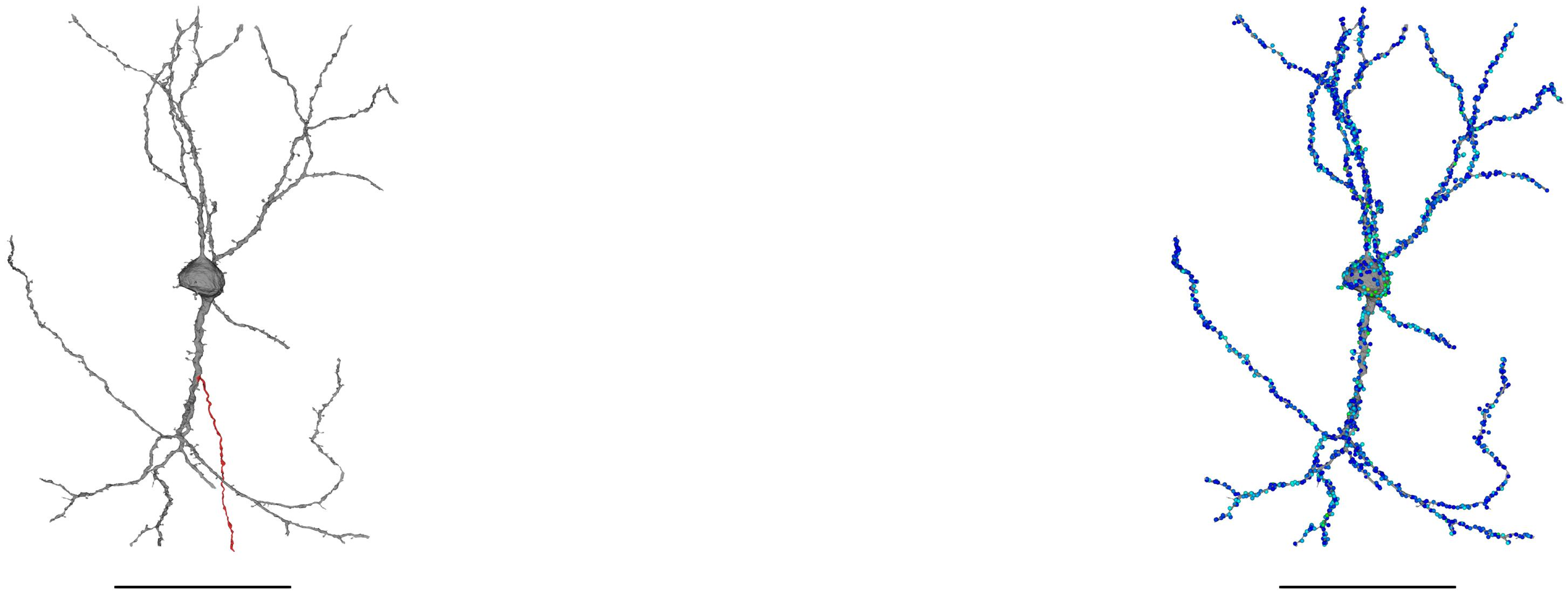

648518346349491816 — Bipolar

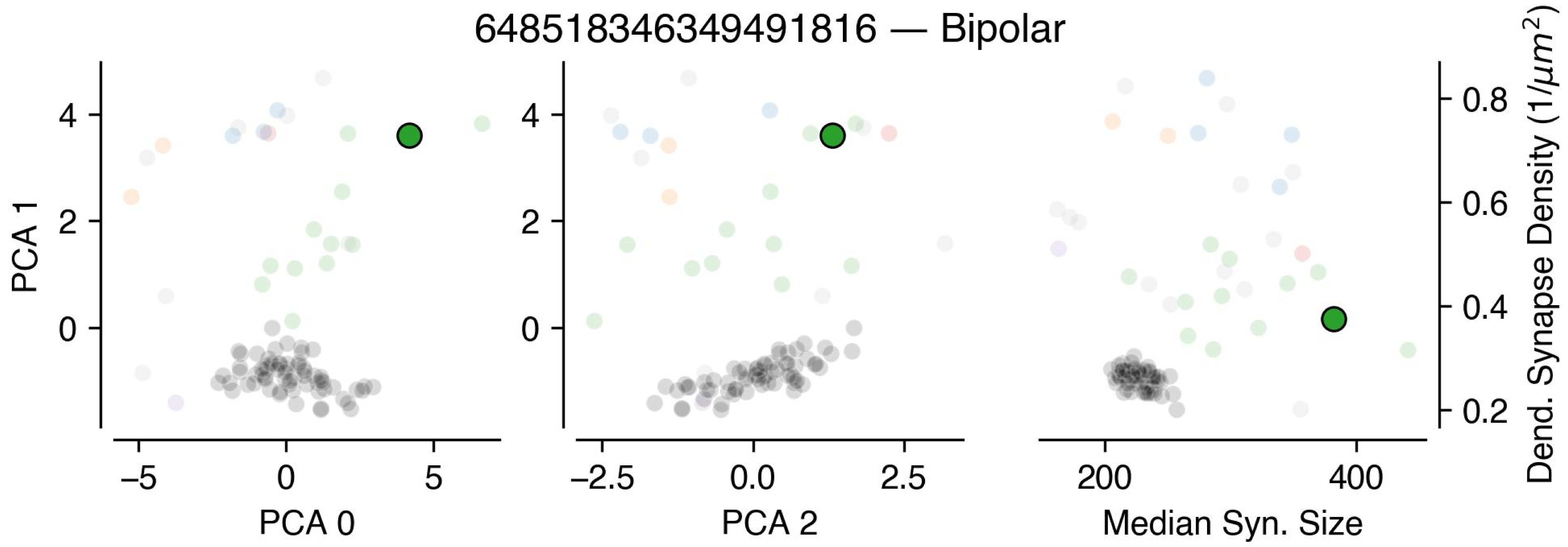


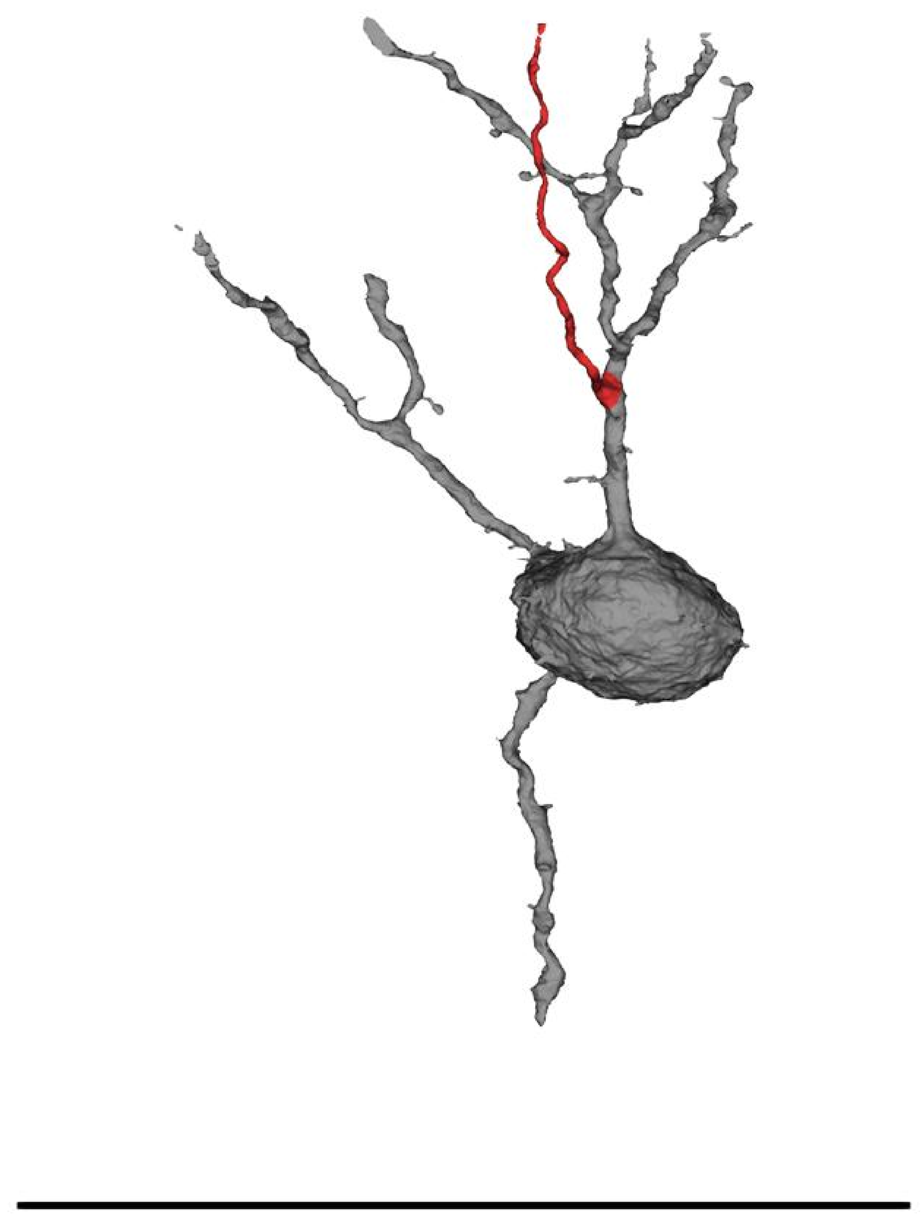

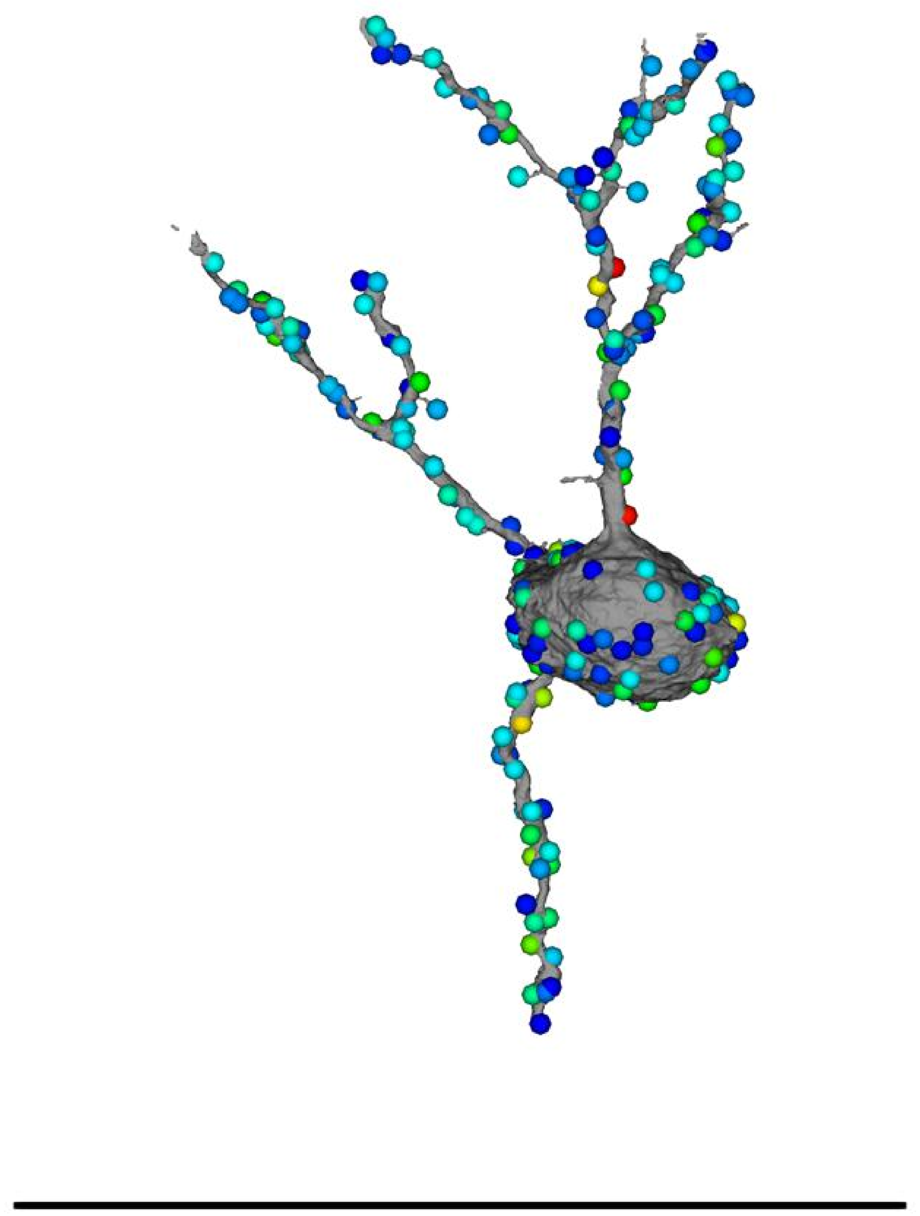

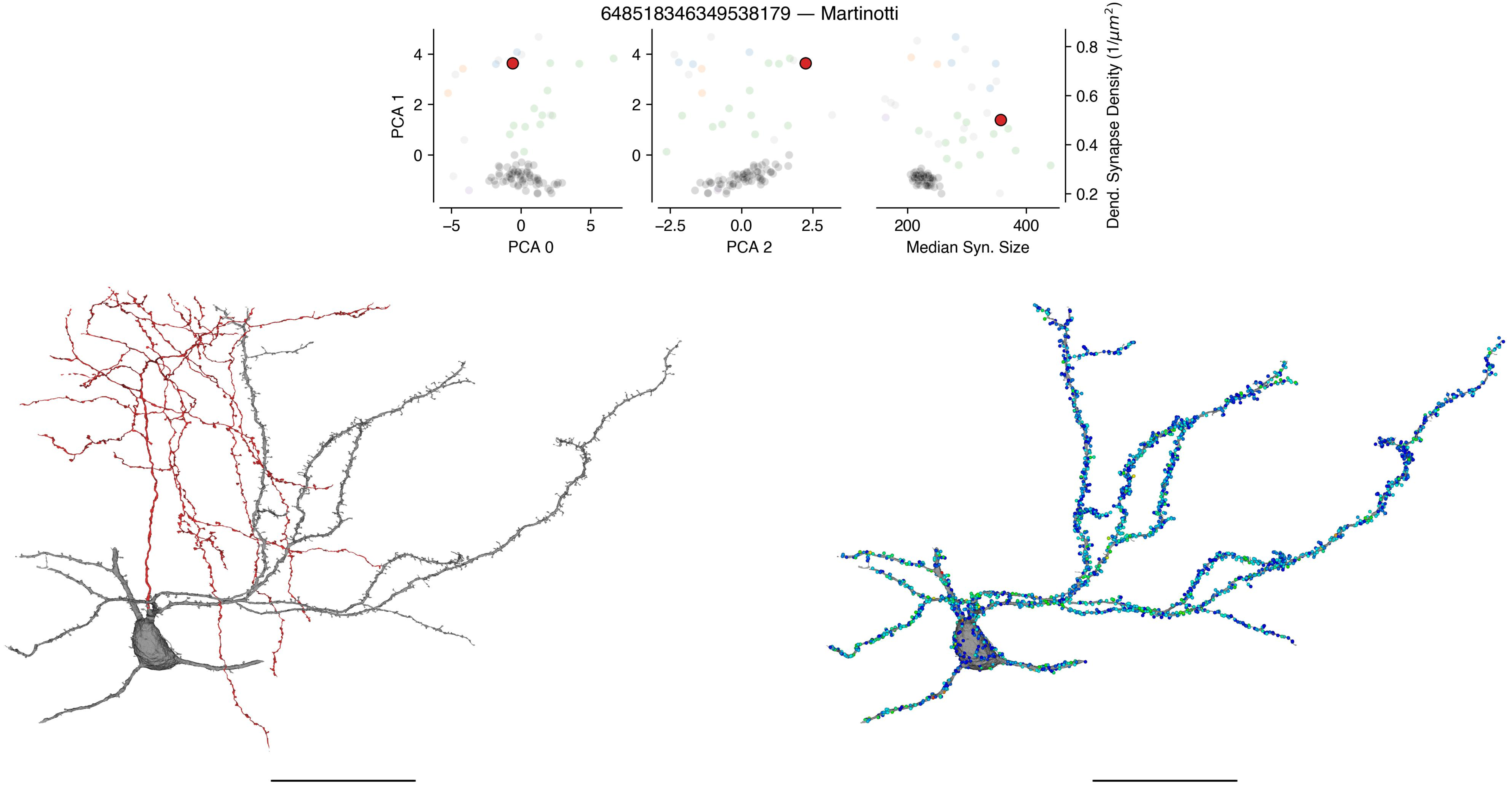
648518346349538179 — Martinotti
PCA 1
PCA 0
PCA 2
Median Syn. Size
Dend. Synapse Density (1/$\mu m^2$)
–5
0
5
–2.5
0.0
2.5
200
400
0
2
4
0.2
0.4
0.6
0.8

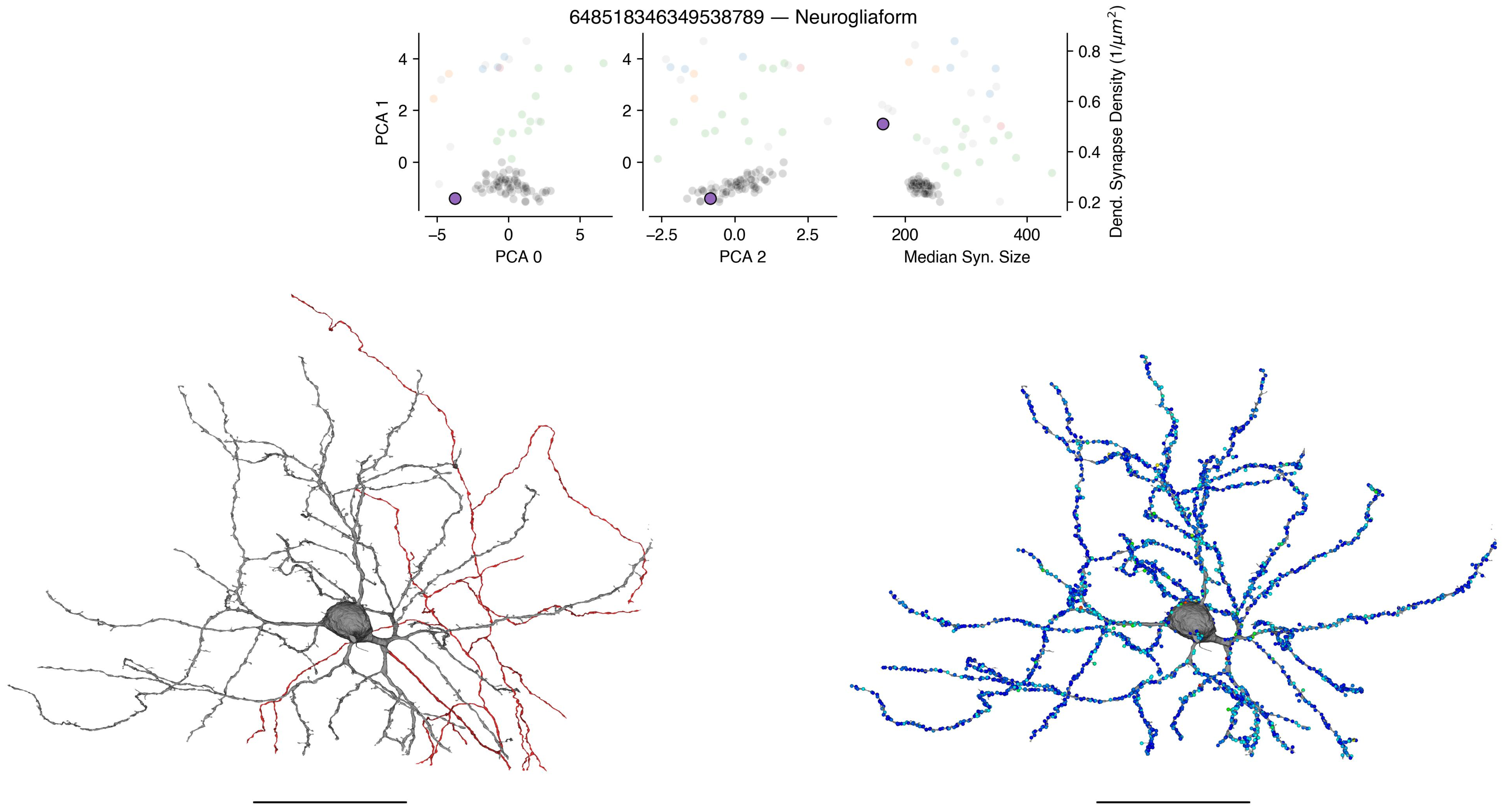
648518346349538789 — Neurogliaform
PCA 1
PCA 0
PCA 2
Median Syn. Size
Dend. Synapse Density (1/μm²)

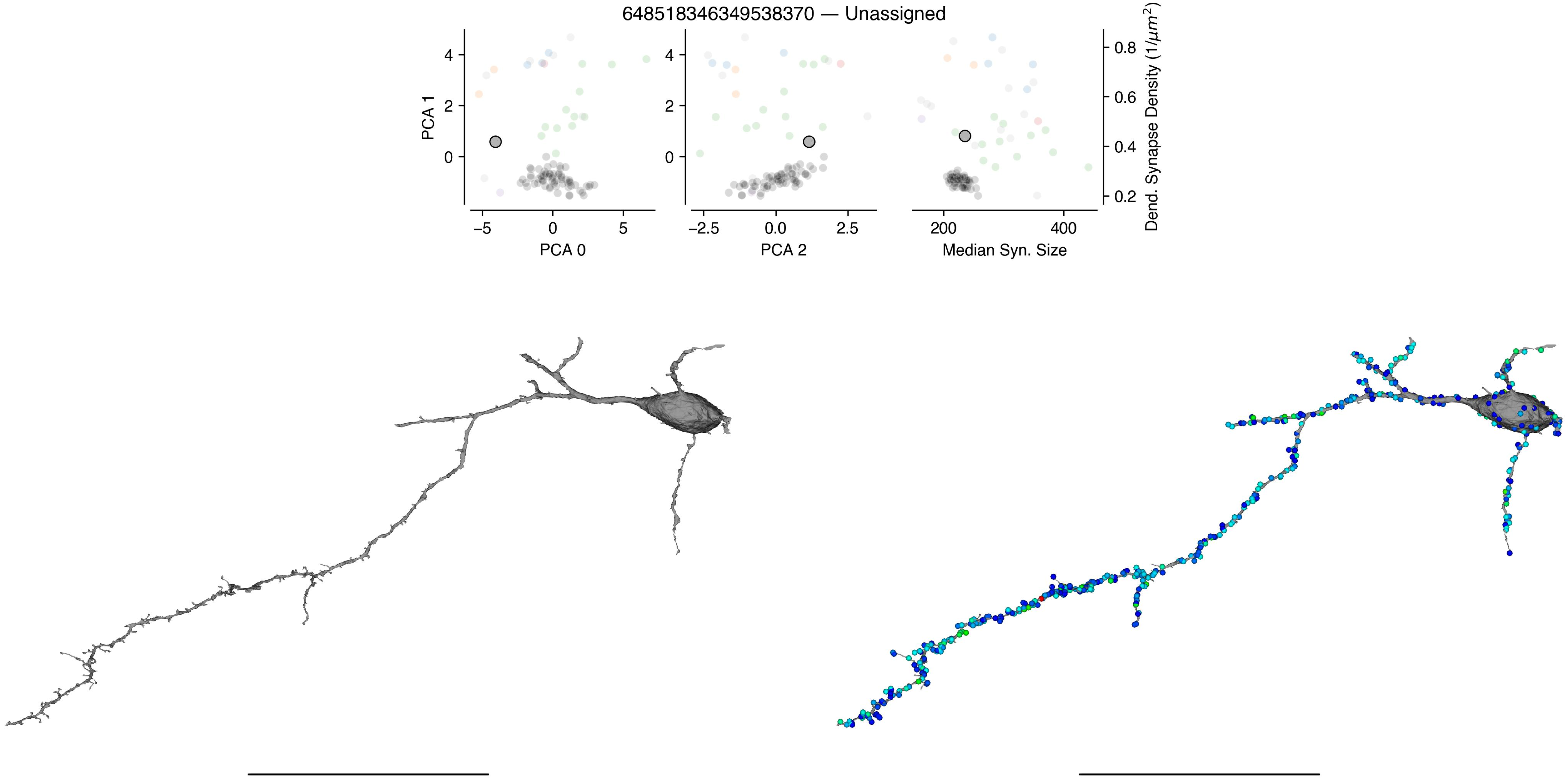

648518346349538370 — Unassigned
PCA 1
PCA 0
PCA 2
Median Syn. Size
Dend. Synapse Density ($1/\mu m^2$)

648518346349488919 — Unassigned

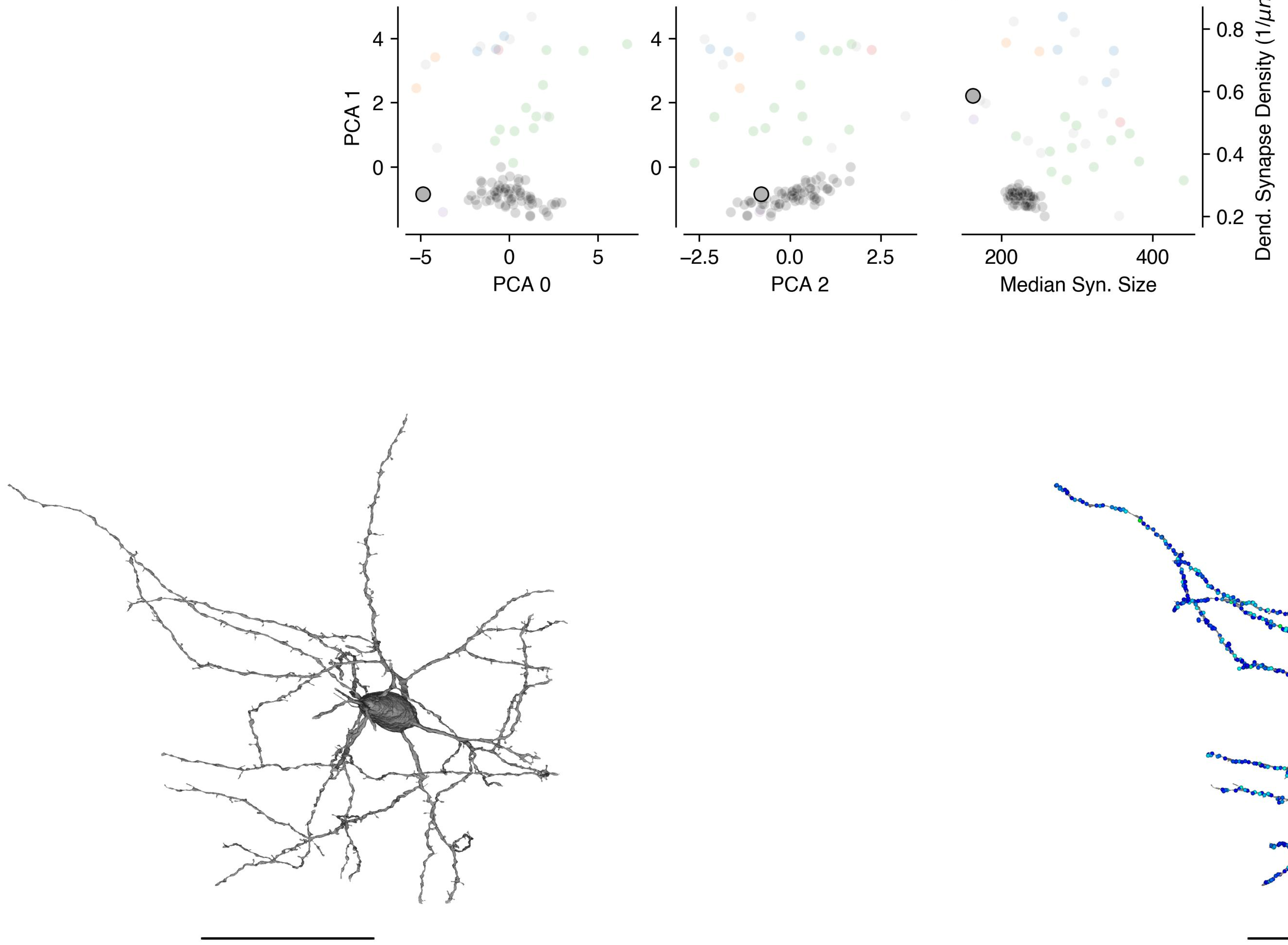

648518346349489985 — Unassigned

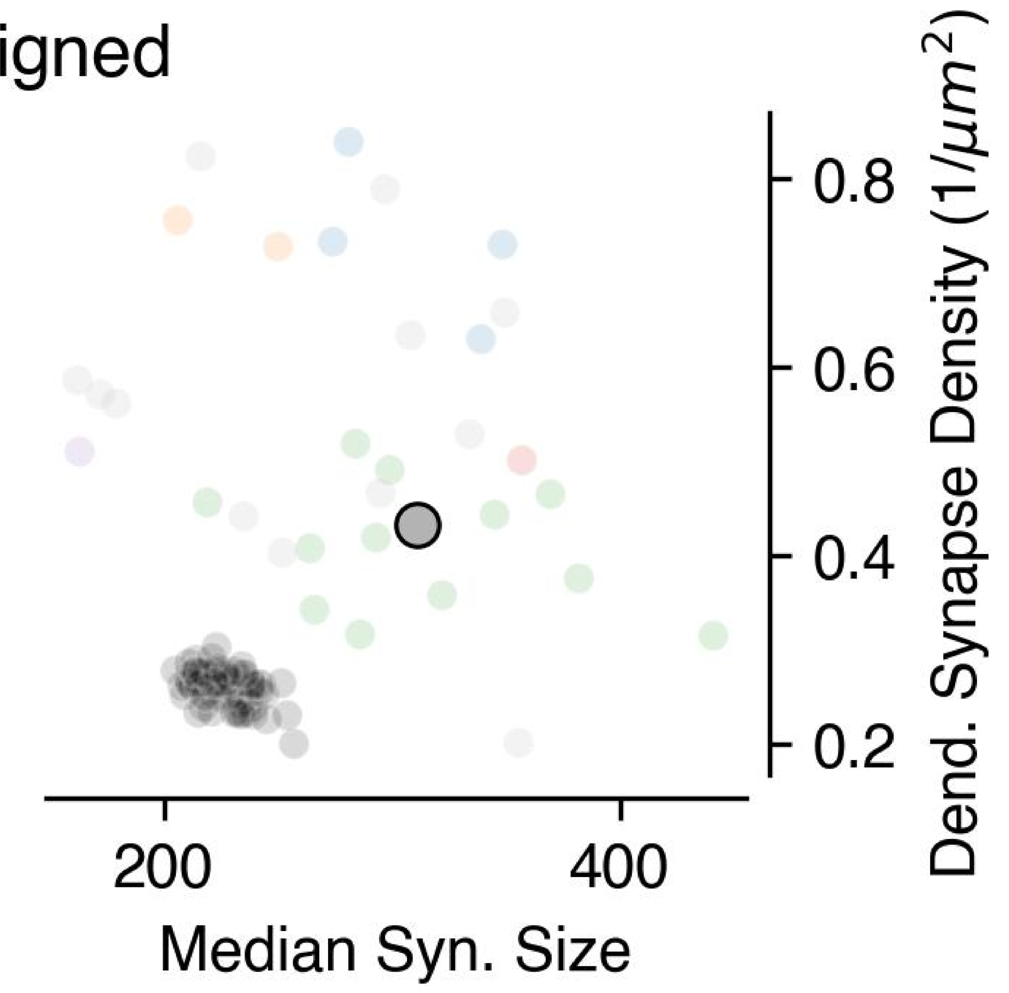


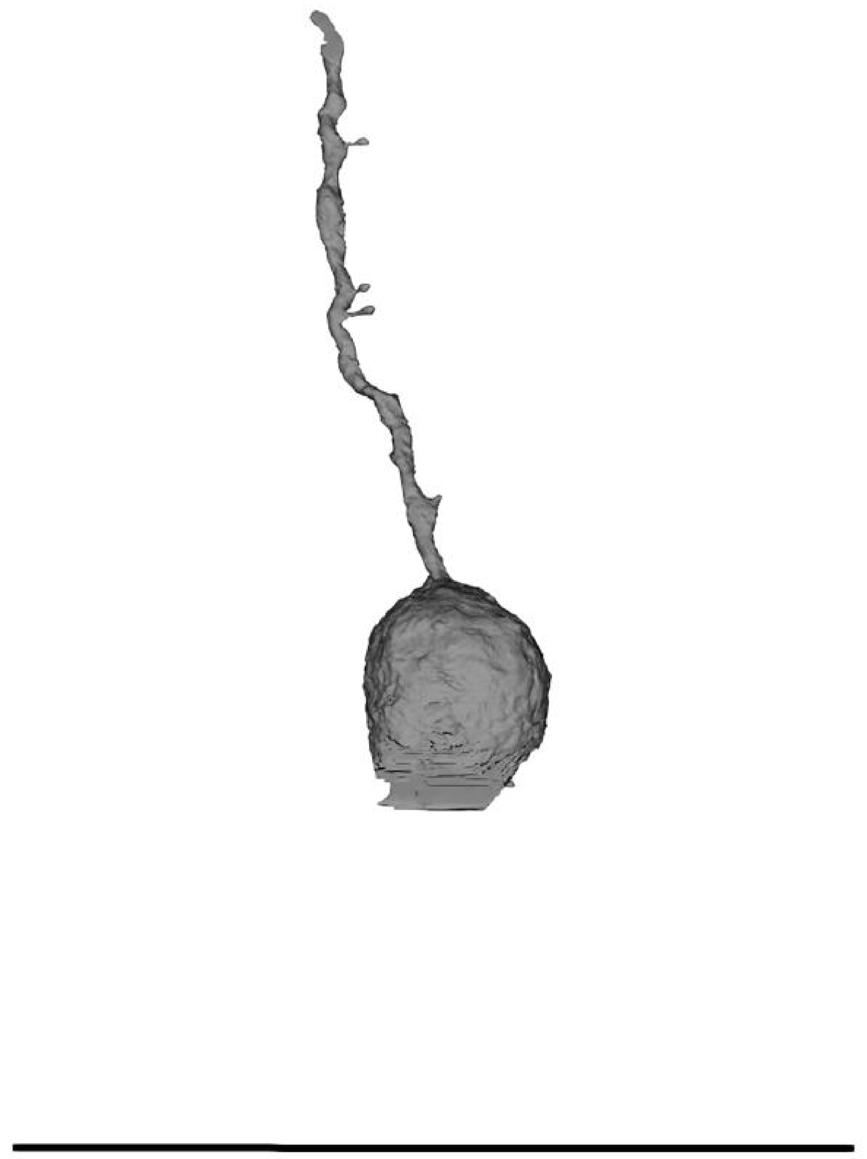

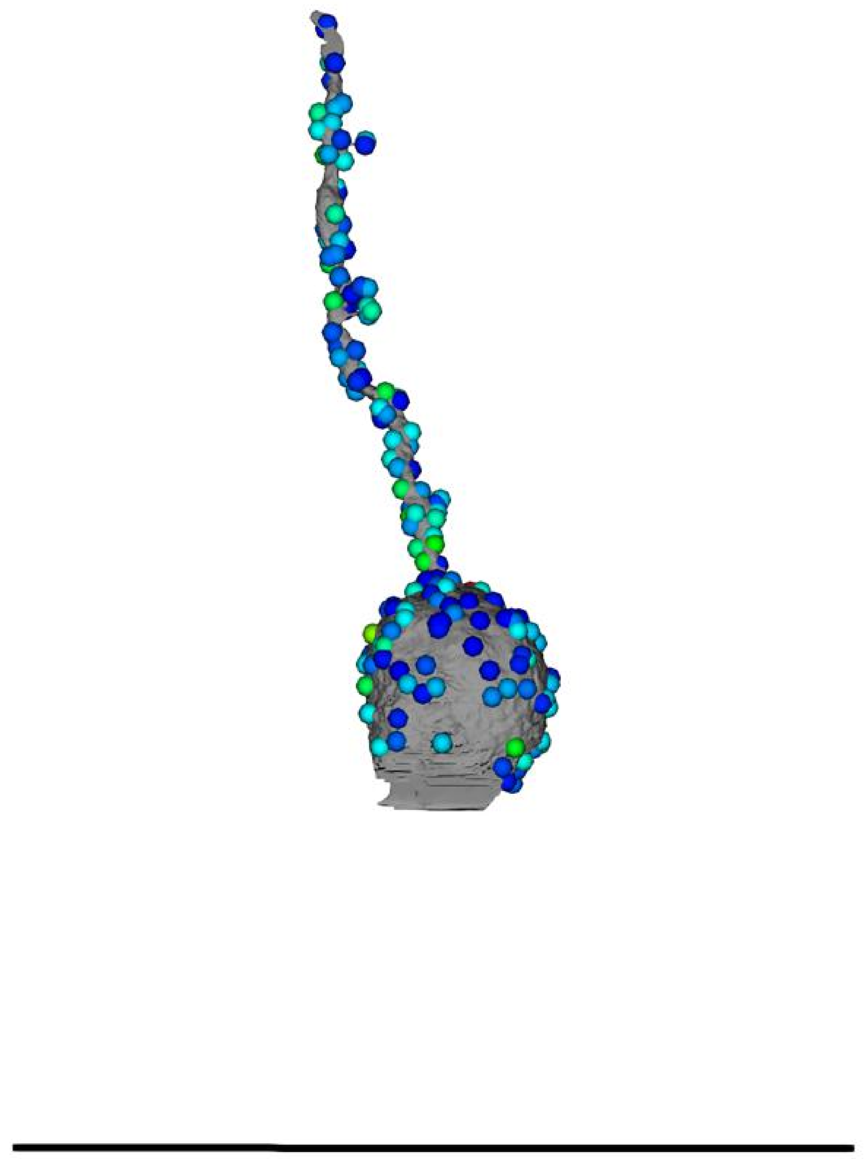

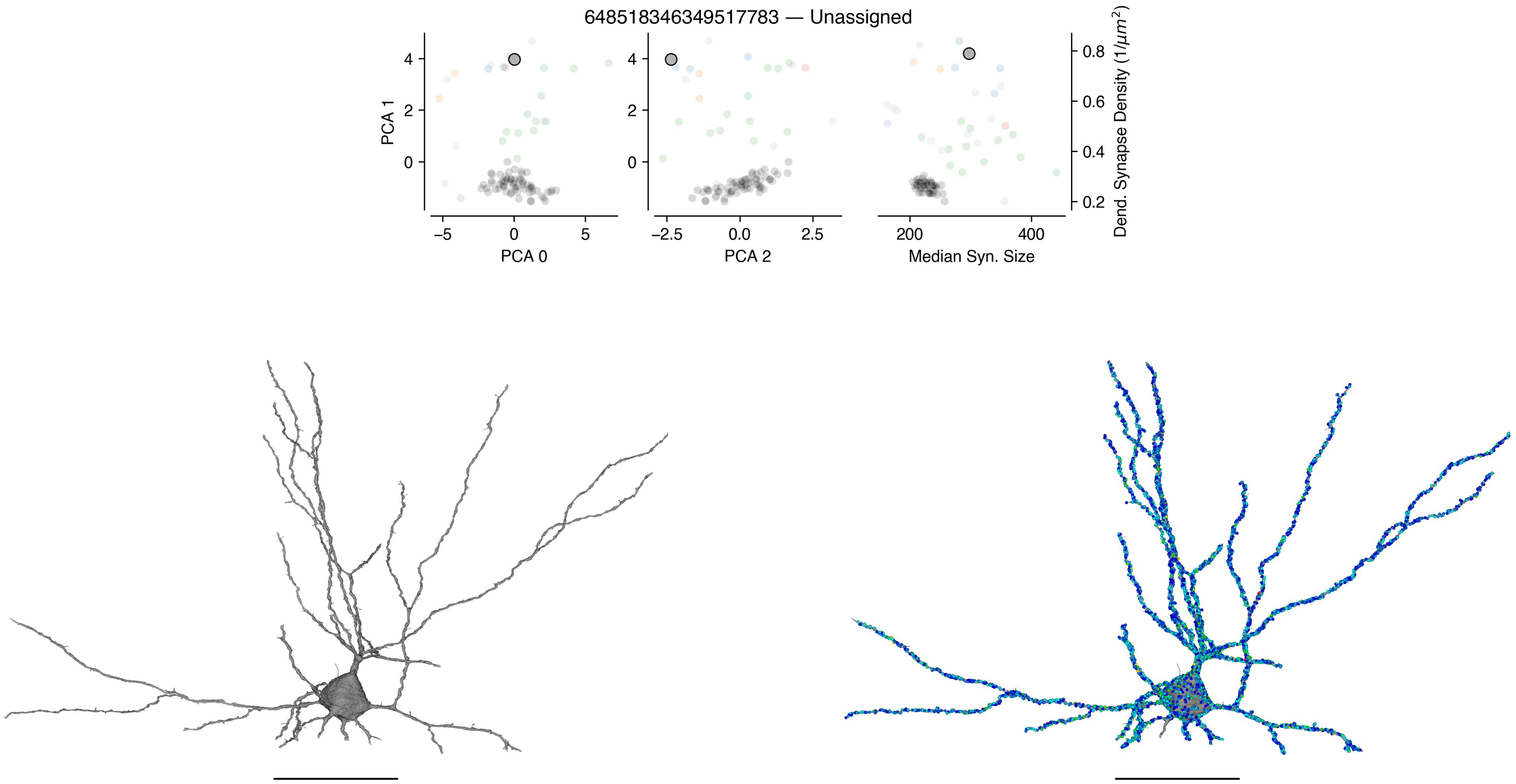
648518346349517783 — Unassigned
PCA 1
PCA 0
PCA 2
Median Syn. Size
Dend. Synapse Density (1/$\mu m^2$)

648518346349526691 — Unassigned

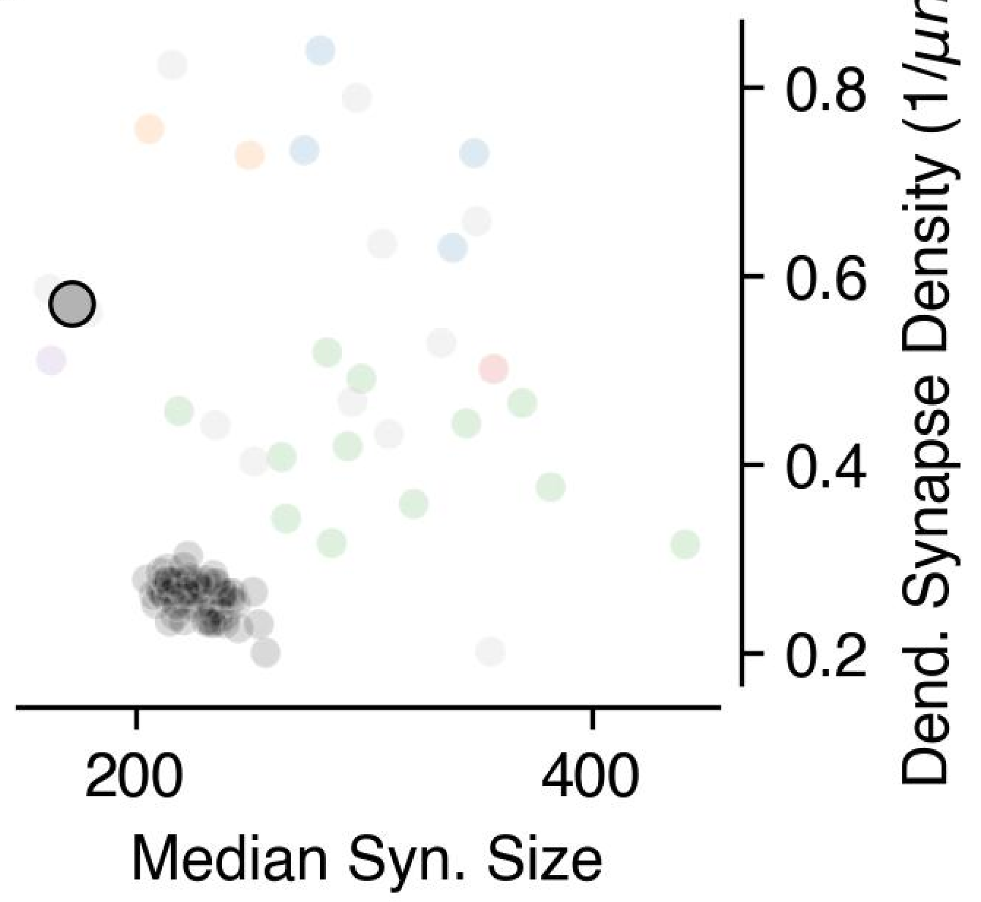


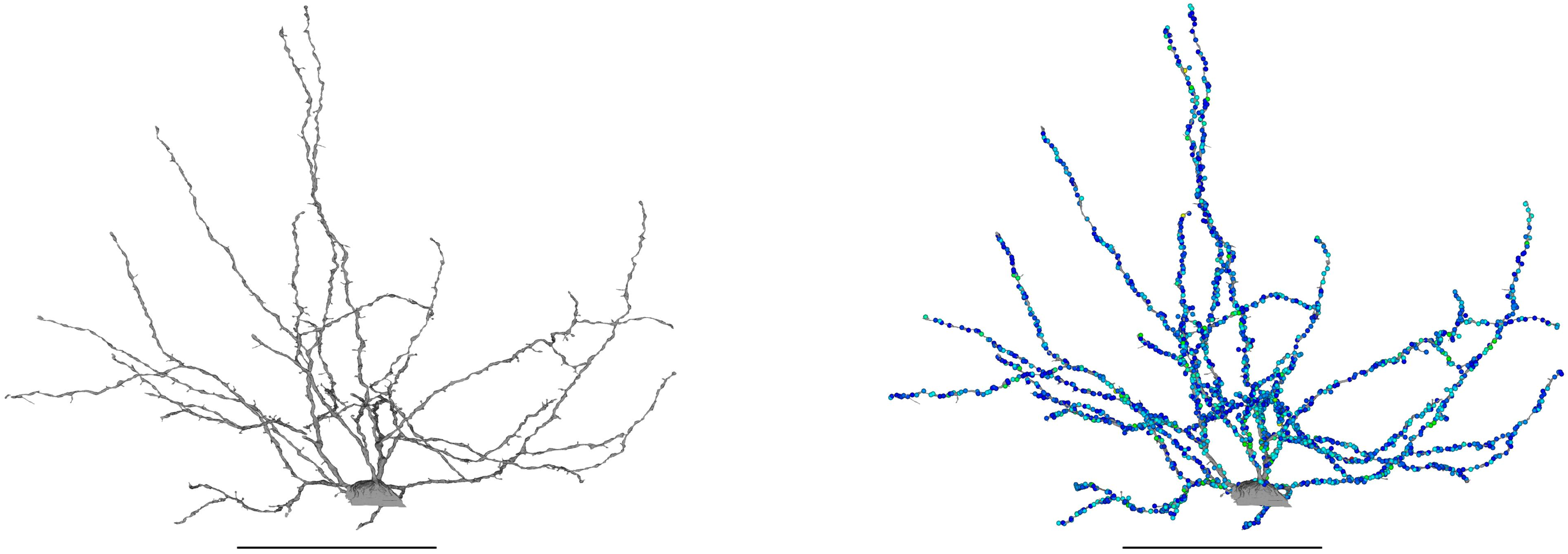

648518346349493894 — Unassigned

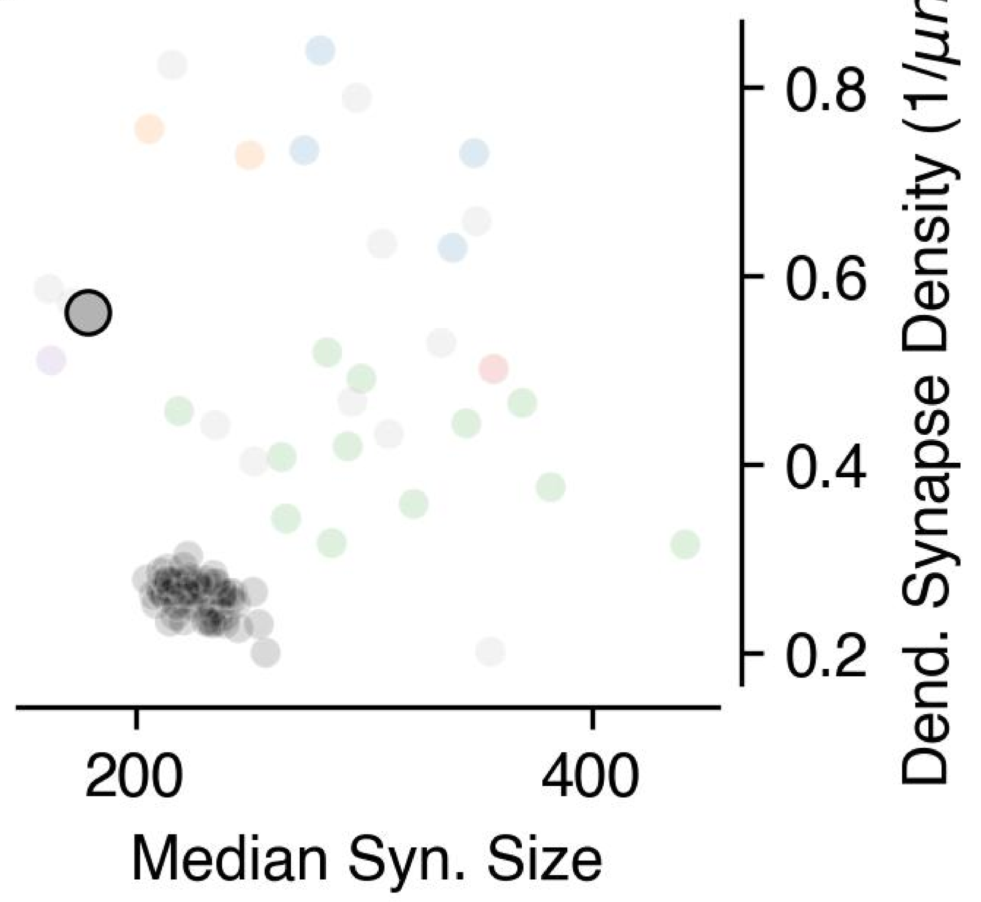


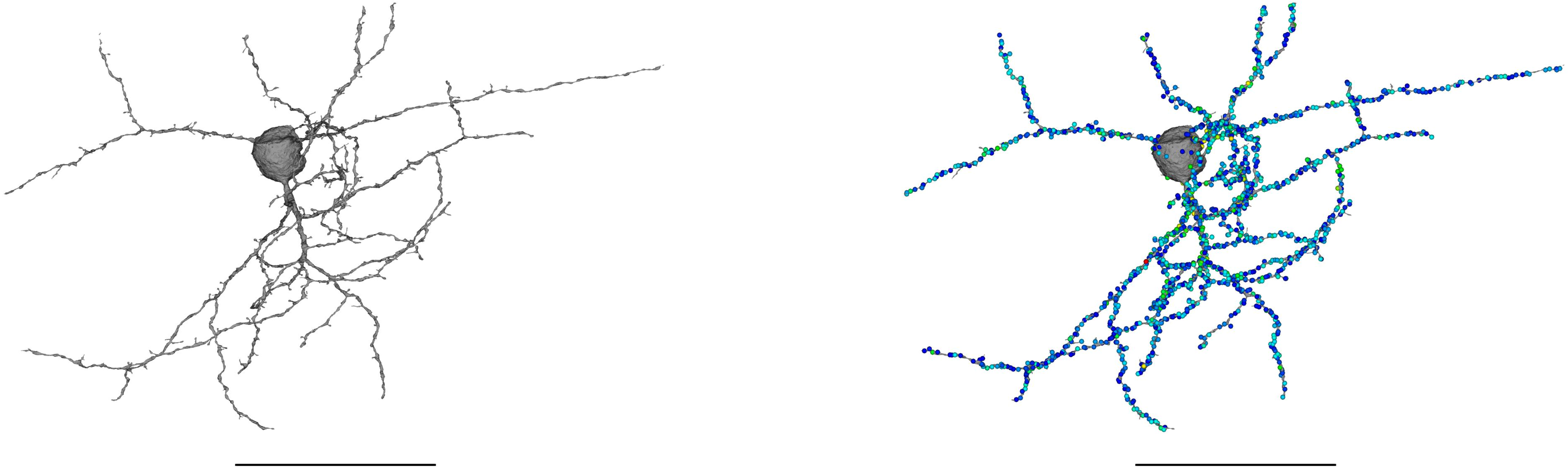

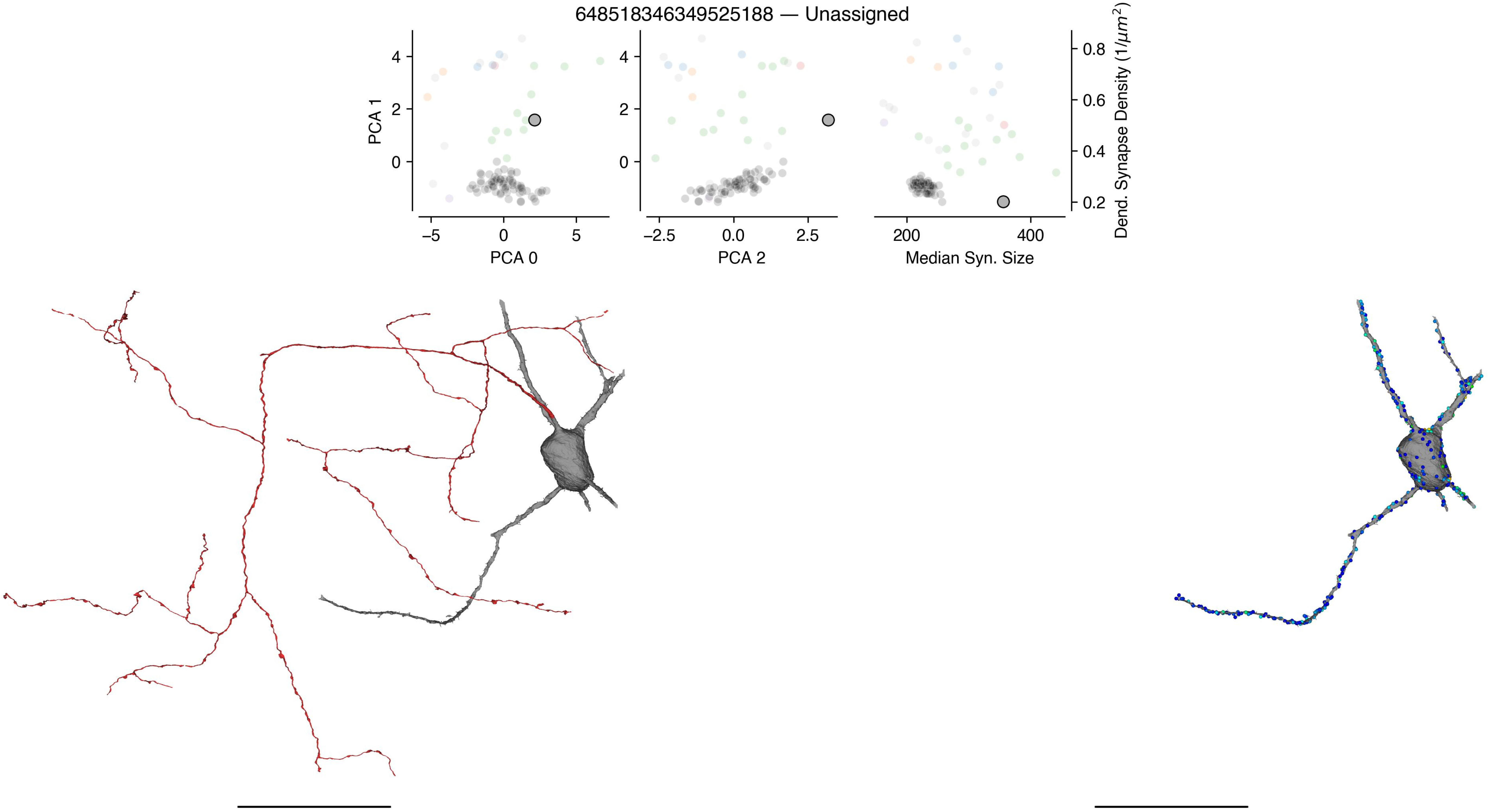

648518346349525188 — Unassigned
PCA 1
PCA 0
PCA 2
Median Syn. Size
Dend. Synapse Density (1/μm²)

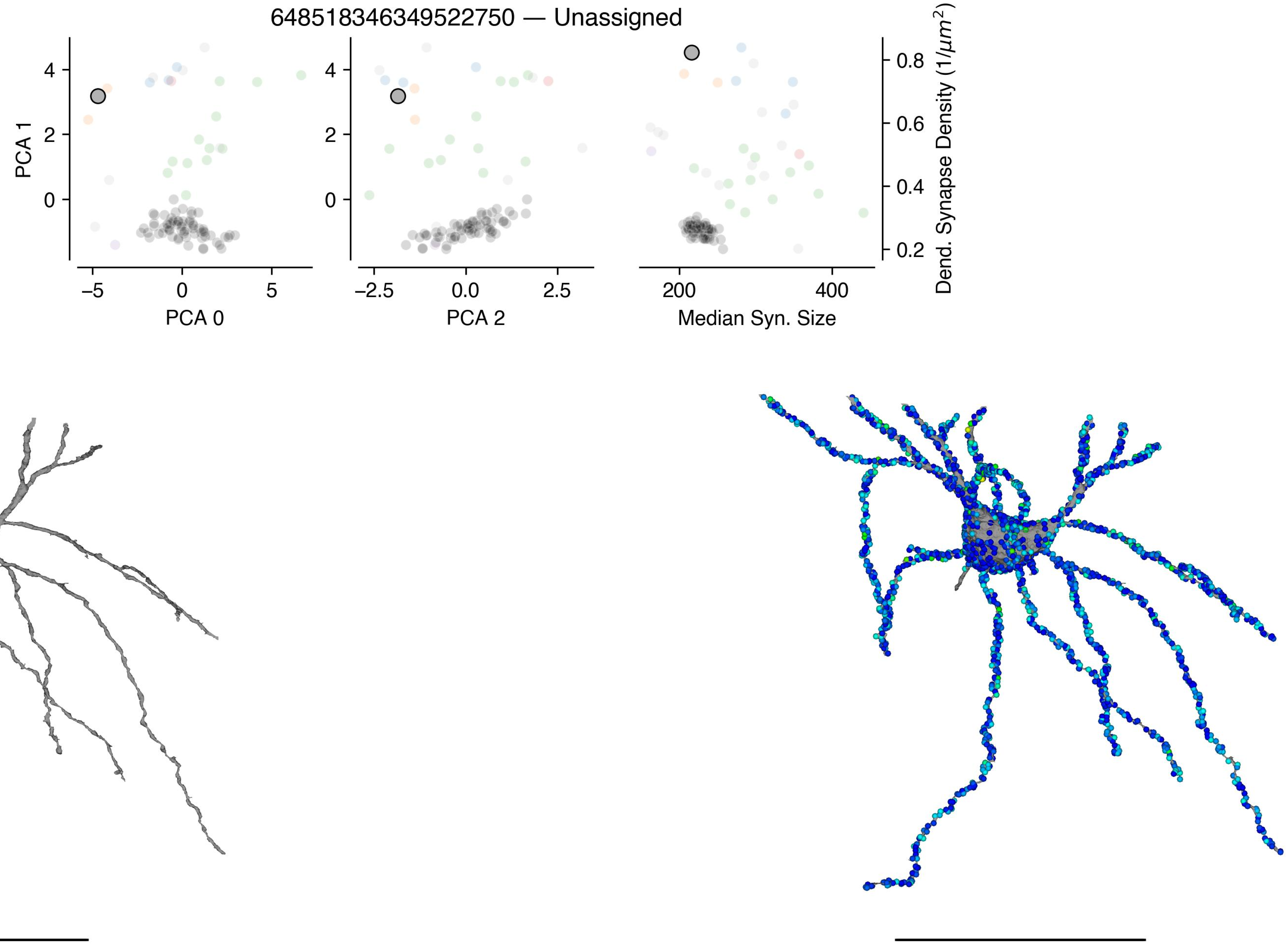
648518346349522750 — Unassigned
PCA 1
PCA 0
PCA 2
Median Syn. Size
Dend. Synapse Density (1/μm²)

648518346349515985 — Unassigned

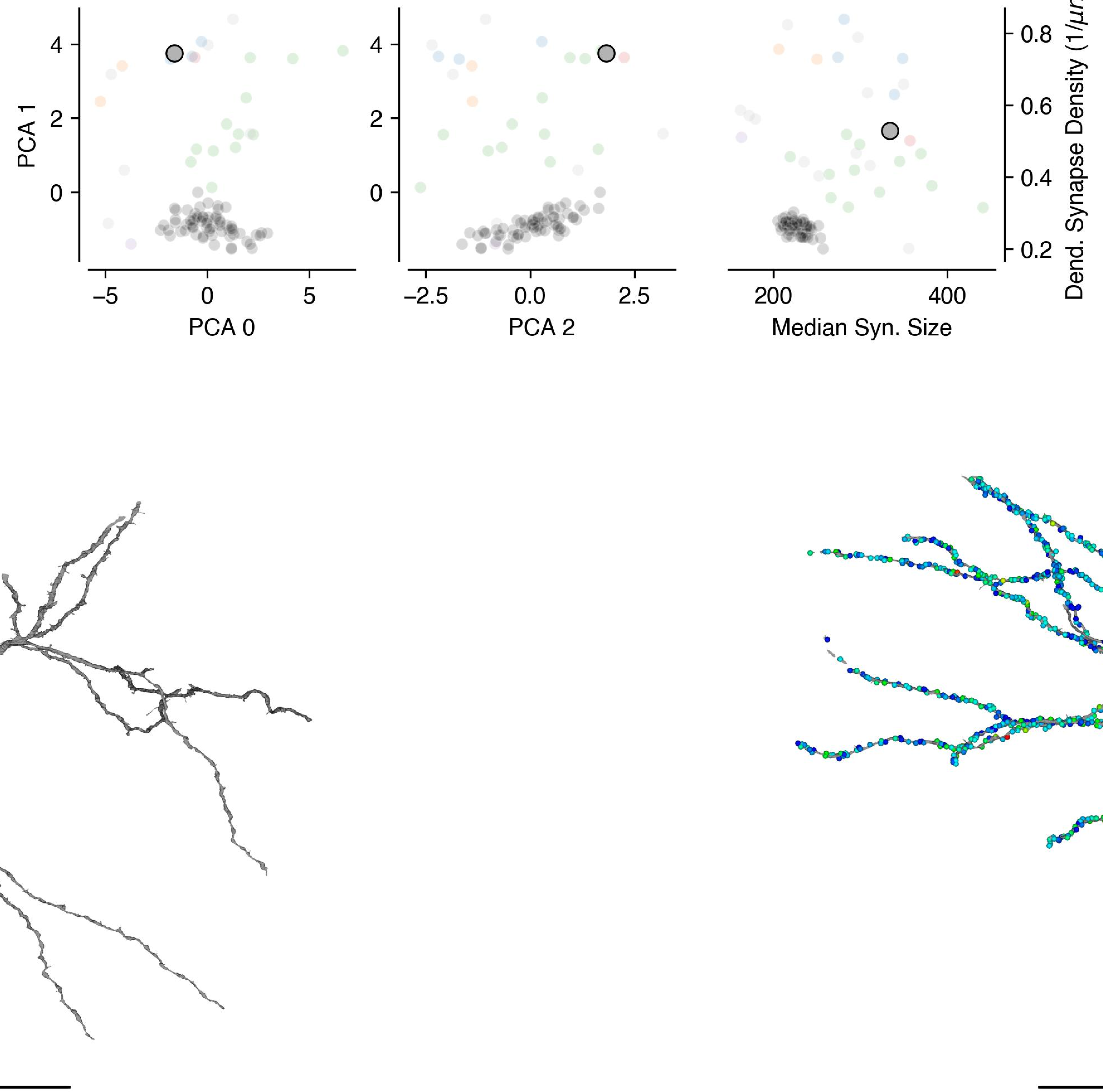

648518346349522740 — Unassigned

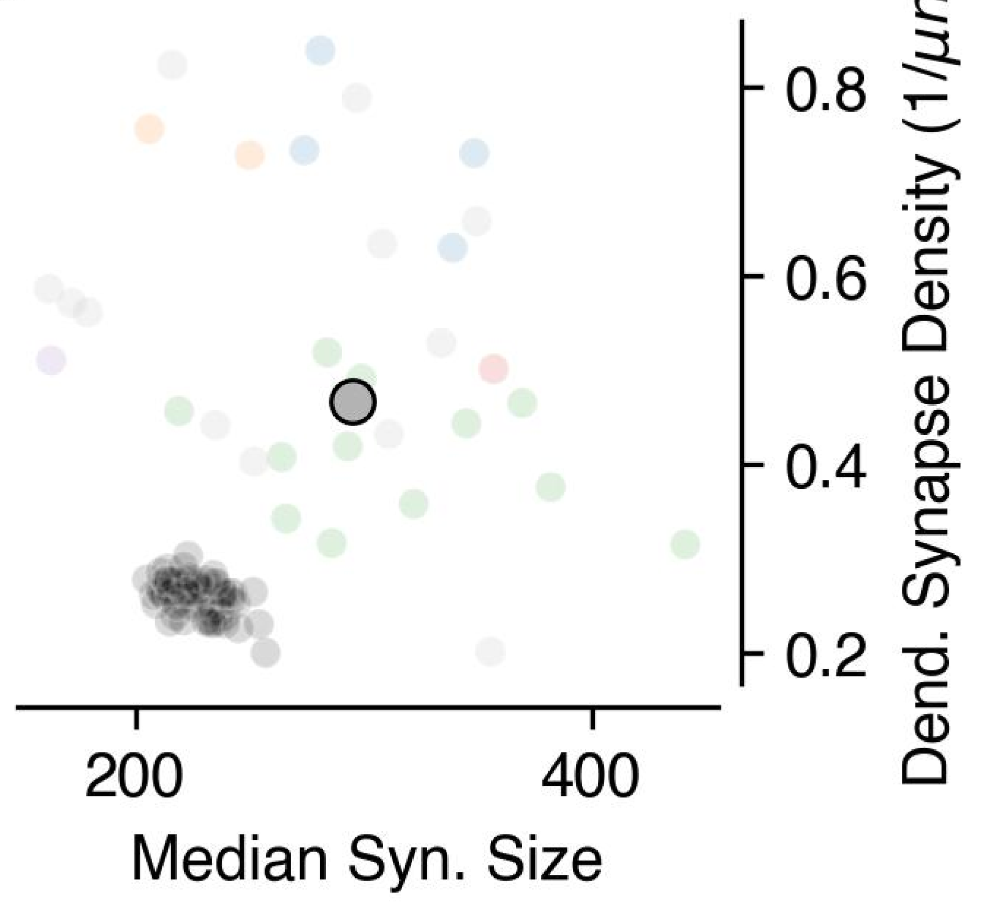


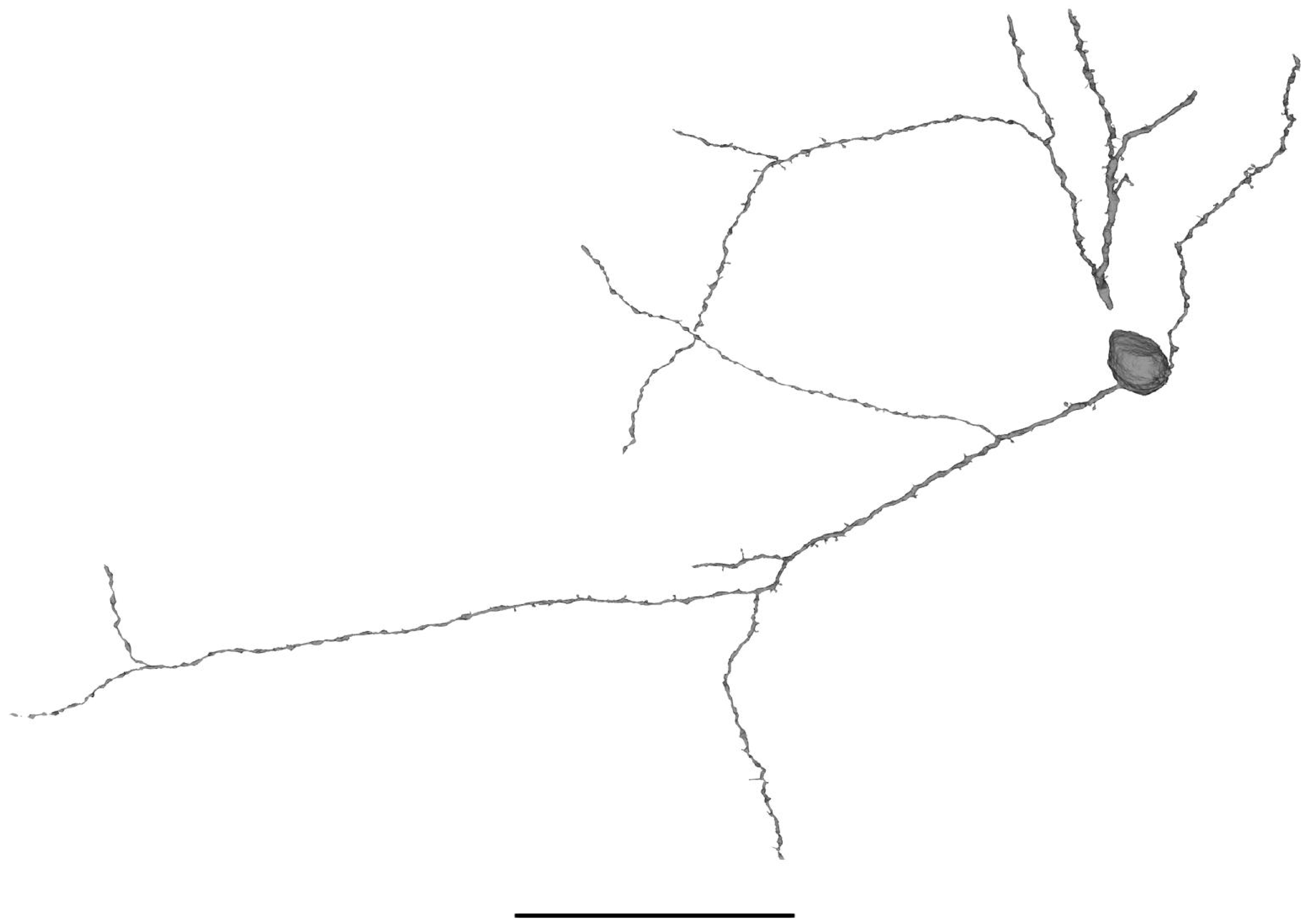

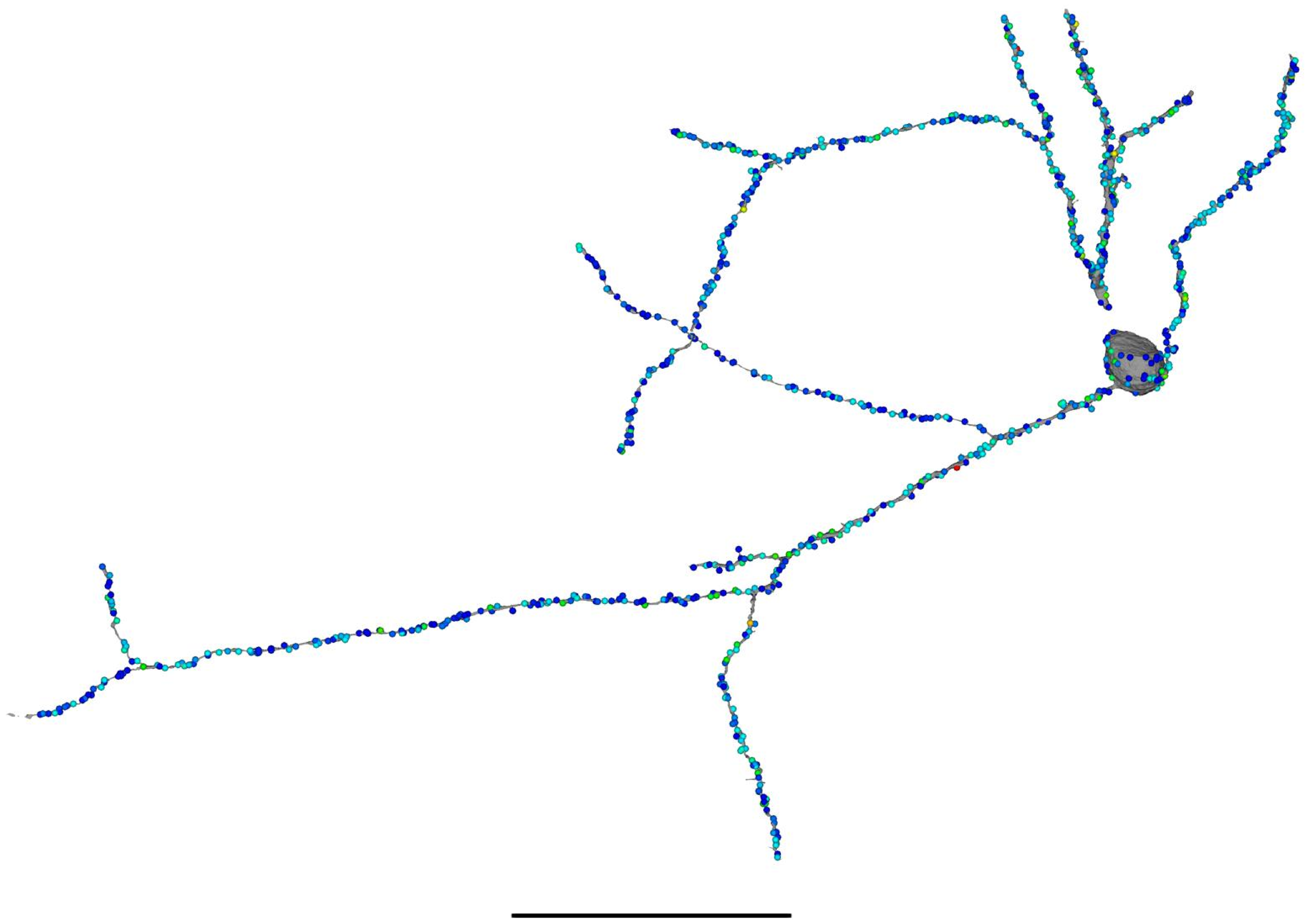

648518346349518096 — Unassigned

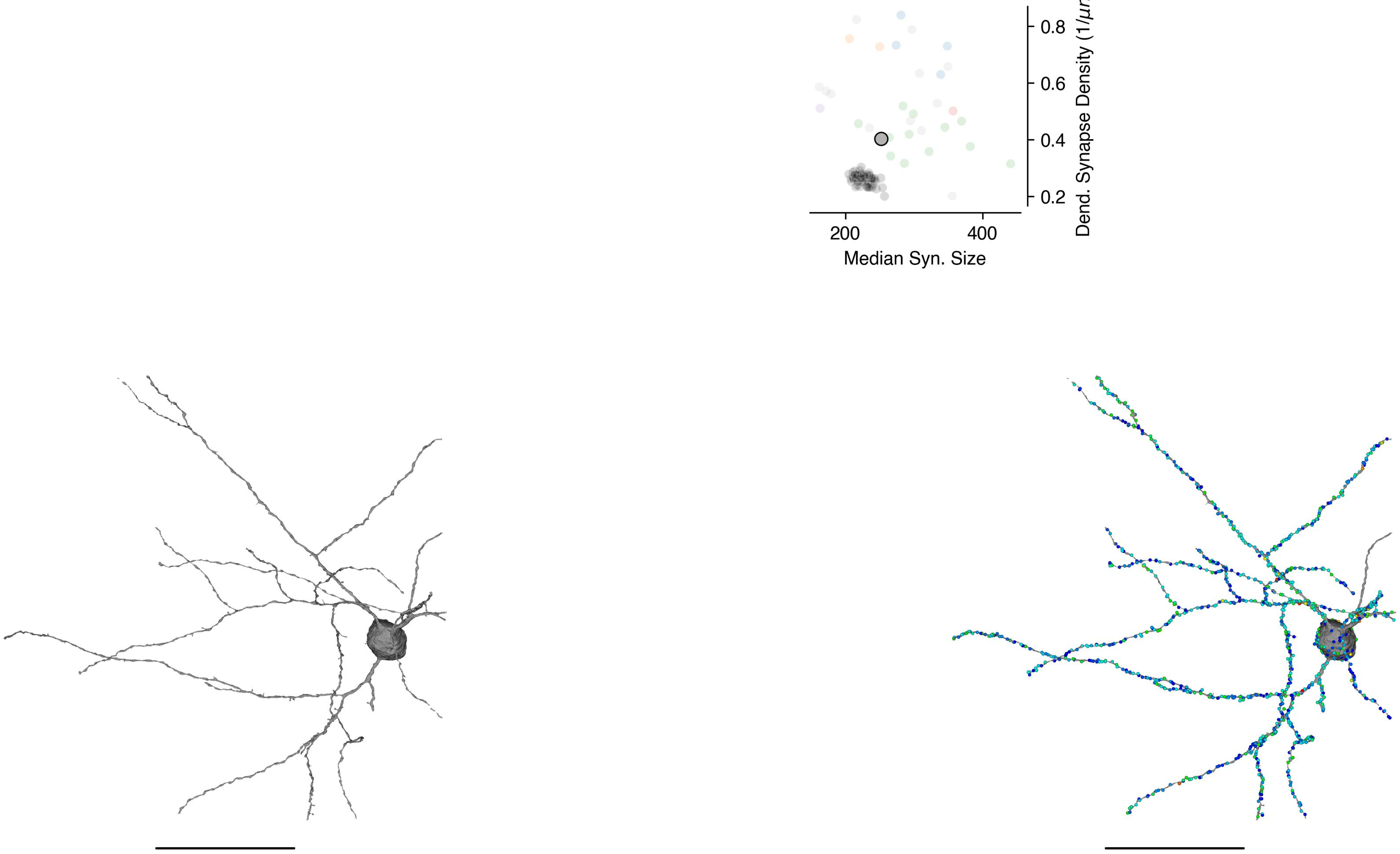

648518346349516055 — Unassigned

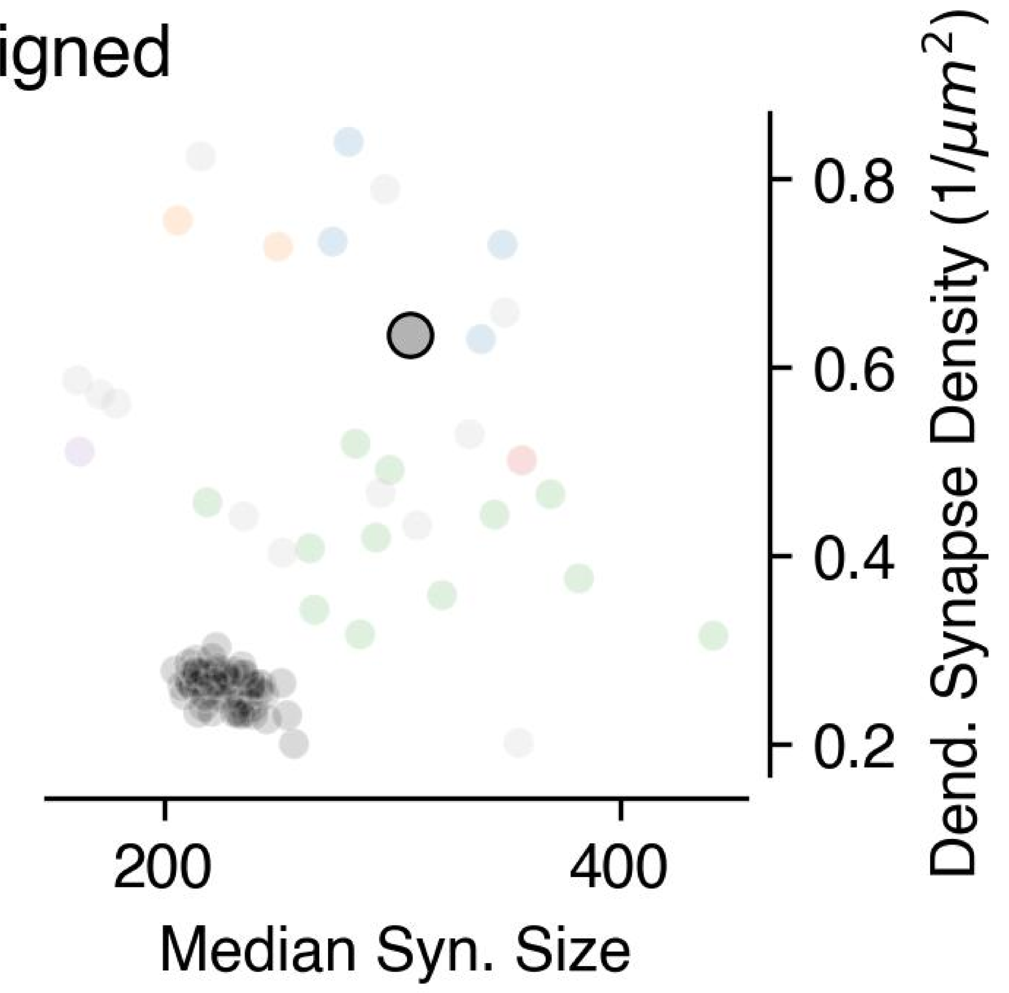


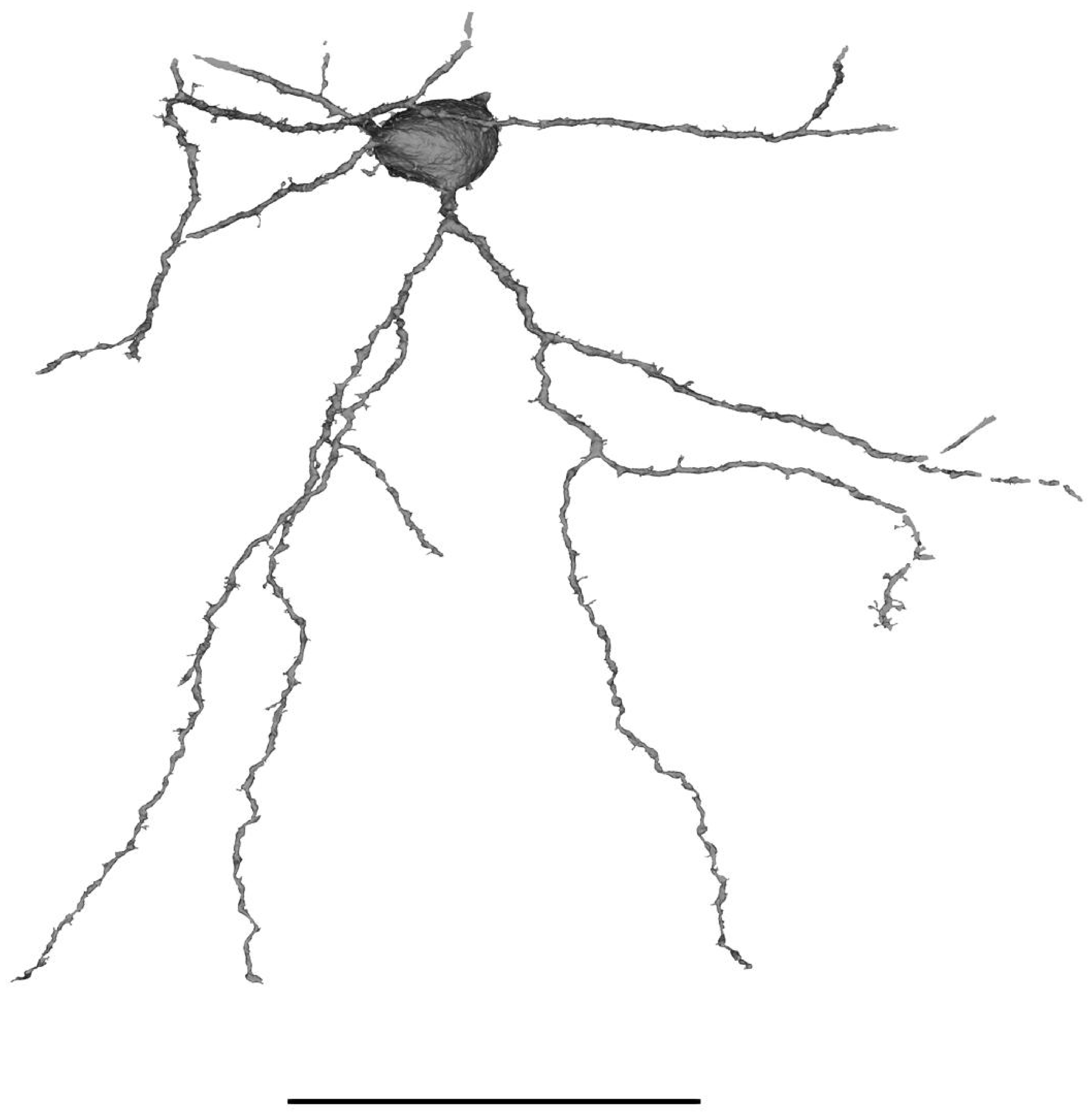

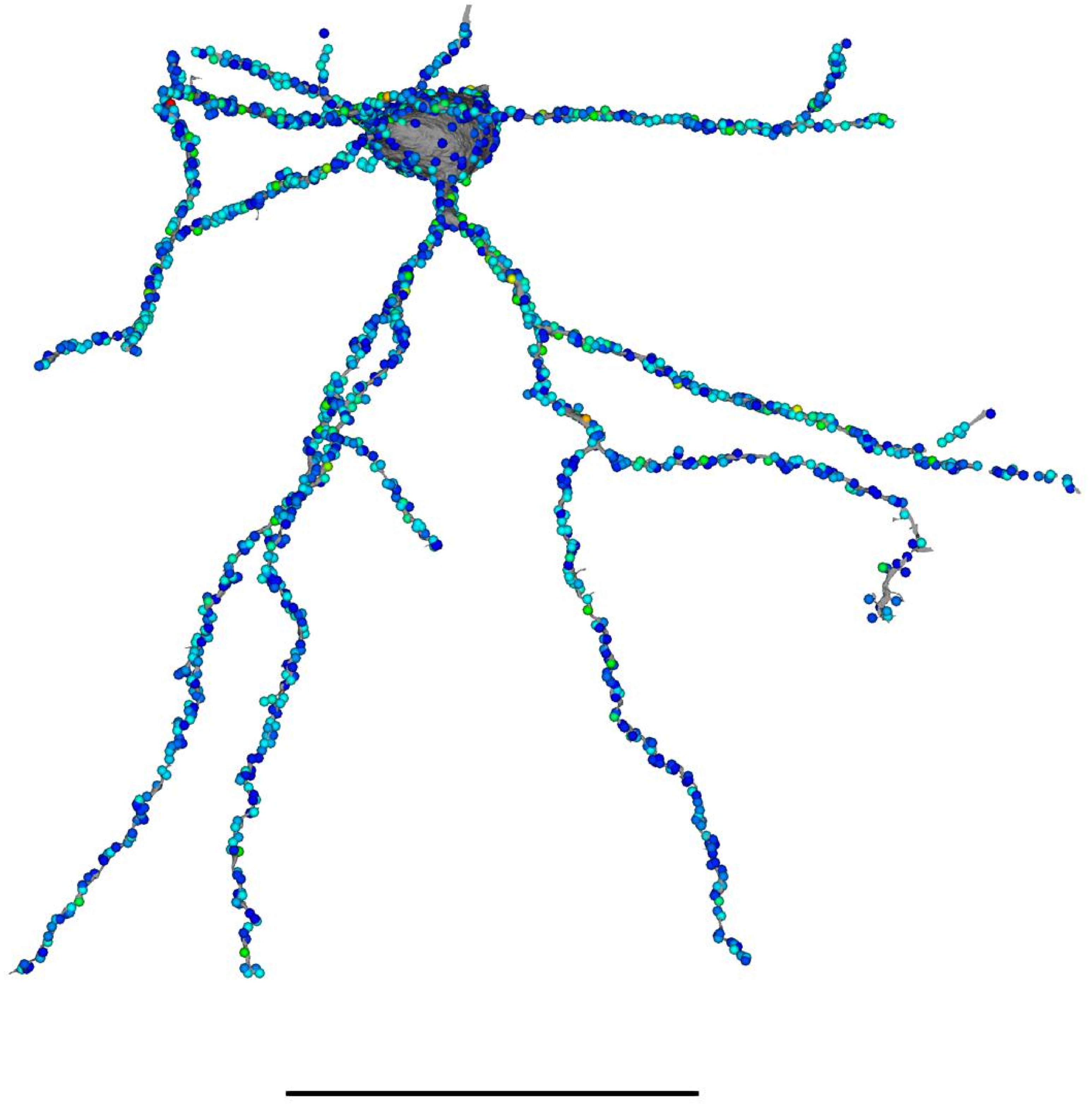

648518346349536849 — Unassigned

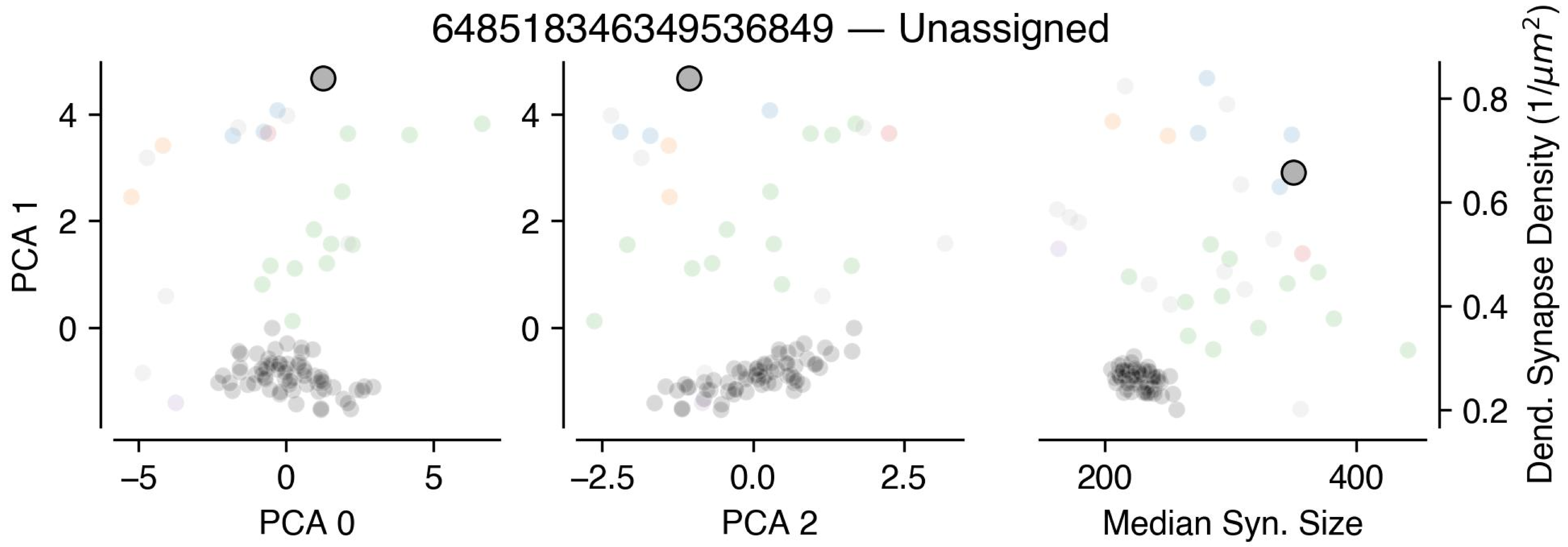


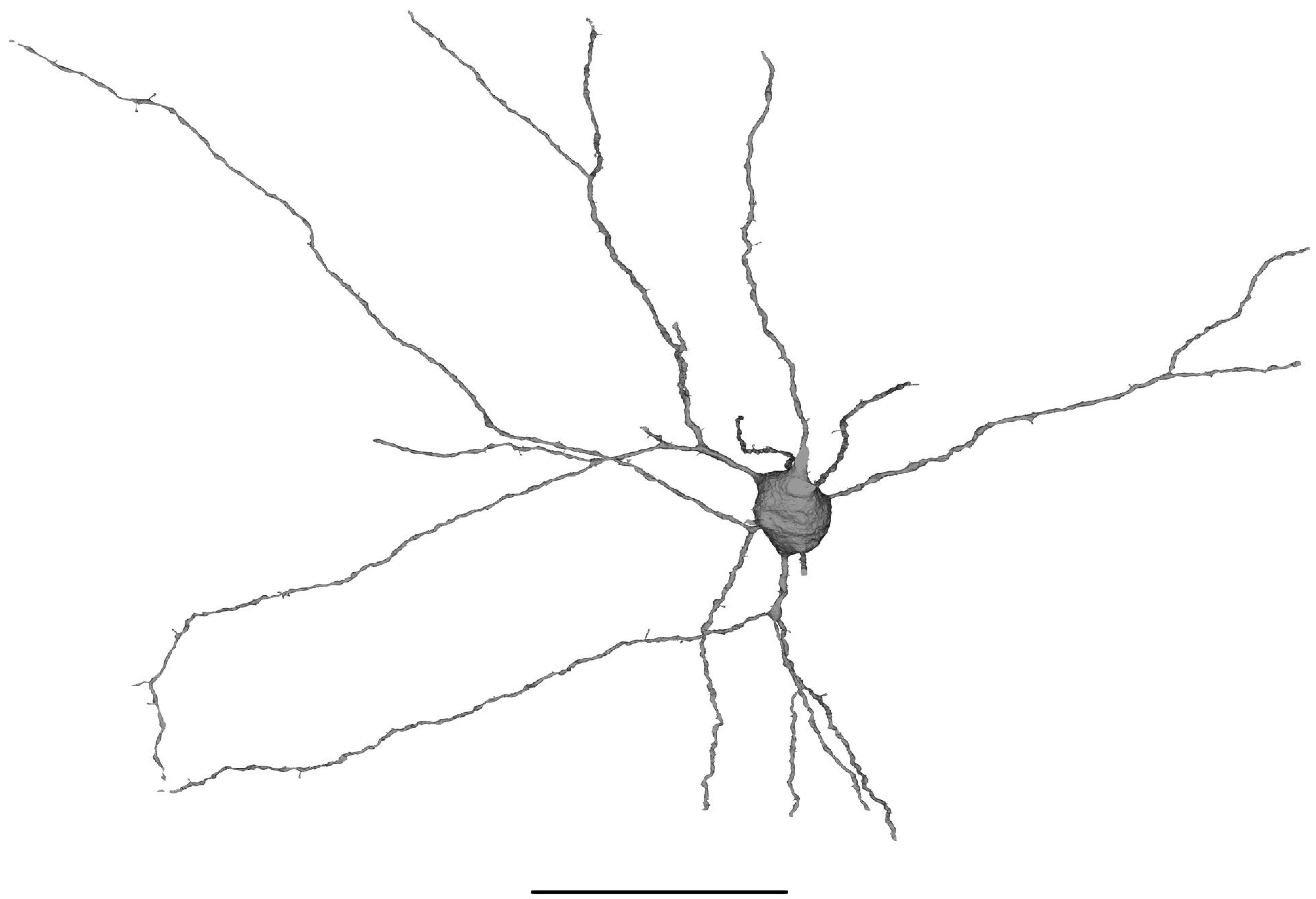

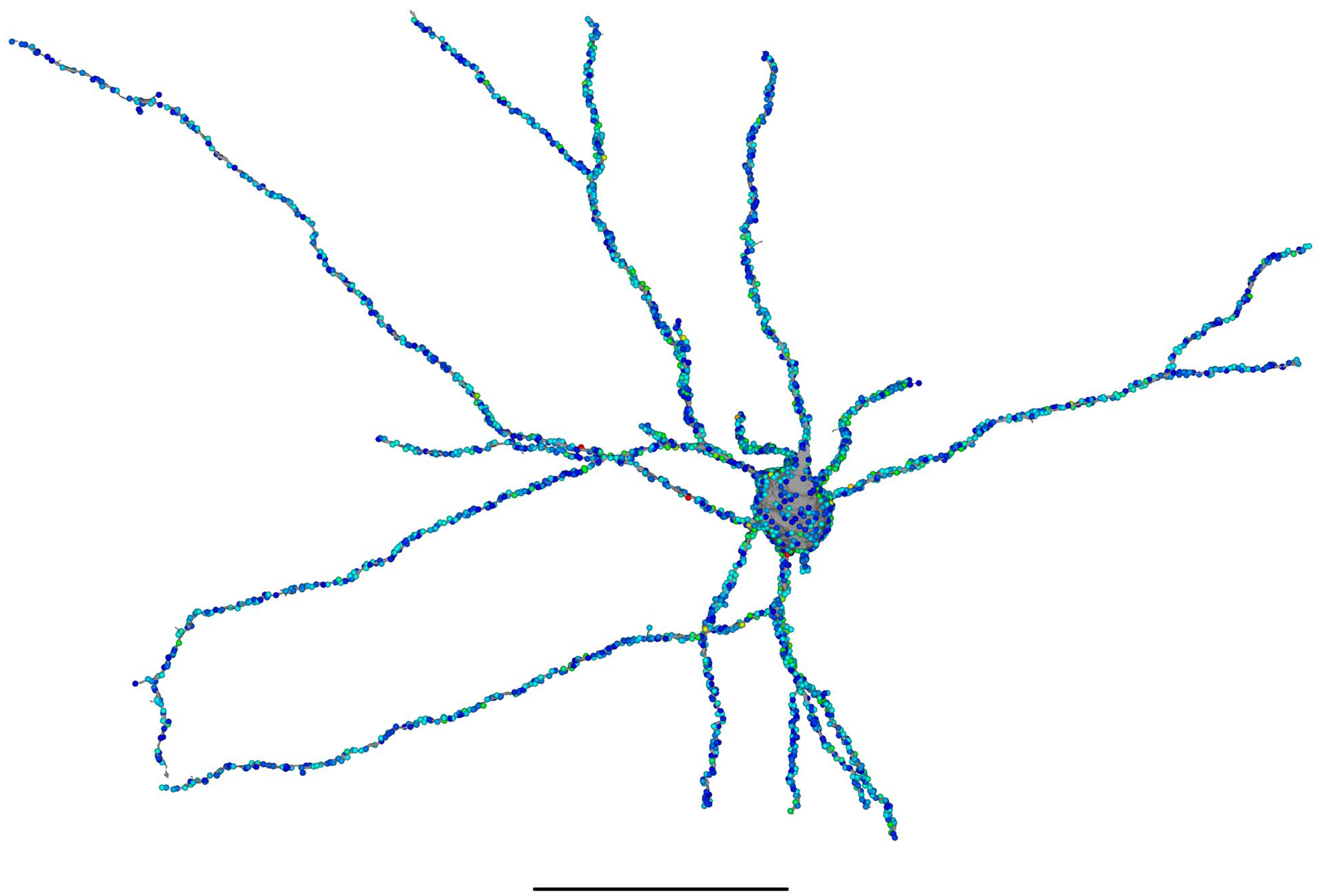
